# Adipogenic activity of chemicals used in plastic consumer products

**DOI:** 10.1101/2021.07.29.454199

**Authors:** Johannes Völker, Felicity Ashcroft, Åsa Vedøy, Lisa Zimmermann, Martin Wagner

**Affiliations:** Department of Biology, Norwegian University of Science and Technology (NTNU), 7491 Trondheim, Norway; Department of Aquatic Ecotoxicology, Goethe University Frankfurt am Main, 60438 Frankfurt am Main, Germany

## Abstract

Bisphenols and phthalates, chemicals frequently used in plastic products, promote obesity in cell and animal models. However, these well-known metabolism disrupting chemicals (MDCs) represent only a minute fraction of all compounds found in plastics. To gain a comprehensive understanding of plastics as a source of exposure to MDCs, we characterized all chemicals present in 34 everyday products using nontarget high-resolution mass spectrometry and analyzed their joint adipogenic activities by high-content imaging. We detected 55,300 chemical features and tentatively identified 629 unique compounds, including 11 known MDCs. Importantly, chemicals that induced proliferation, growth, and triglyceride accumulation in 3T3-L1 adipocytes were found in one third of the products. Since the majority did not target peroxisome proliferator-activated receptor γ, the effects are likely to be caused by unknown MDCs. Our study demonstrates that daily-use plastics contain potent mixtures of MDCs and can, therefore, be a relevant yet underestimated environmental factor contributing to obesity.

**Teaser:** Plastics contain a potent mixture of chemicals promoting adipogenesis, a key process in developing obesity.

## Introduction

Obesity is a global pandemic that generates a considerable burden of disease, in particular considering comorbidities such as type 2 diabetes, cardiovascular diseases, hypertension, non-alcoholic fatty liver, stroke, and certain types of cancer (*1*). Worldwide, the number of obese people has nearly tripled since 1975, and more than 41 million children under the age of five were overweight or obese in 2016 (*2*). This is problematic since a high body mass index (BMI) is one of the top risk factors for deaths (*3*), and overweight in childhood or adolescence is a good predictor of adult obesity (*4*). Accordingly, a high BMI contributed to four million deaths globally in 2015 with cardiovascular diseases as leading cause of death followed by diabetes, chronic kidney diseases and cancer (*5*).

This public health problem has mainly been attributed to genetic background and changes in lifestyle, such as diet, exercise, sleep deficiency, and aging. However, epidemiological evidence suggests that these factors insufficiently explain the magnitude and speed of the obesity pandemic (*6*). For instance, even when normalizing for caloric intake and exercise, the BMI of US adults increased by 2.3 kg m^−2^ in 2006 compared to 1998 (*6*). Consequently, identifying and understanding other environmental factors than lifestyle is crucial to manage obesity (*7*). Given that the endocrine system controls appetite, satiety, metabolism, and weight, exposure to endocrine disrupting chemicals is one such factor (*8*). In addition to prominent endocrine disruptors, such as the biocide tributyltin and the pesticide dichlorodiphenyltrichloroethane (DDT), plastic chemicals such as bisphenols or phthalates disrupt metabolic functions and promote obesity in cell and animal experiments (*8*). This is further supported by epidemiological studies that have linked weight gain in humans to bisphenol A (BPA) exposure (*9*), while contradicting outcomes have been reported regarding a link to phthalate exposure (*10–12*).

Considering the chemical complexity of plastic consumer products, bisphenols and phthalates represent only the tip of the iceberg. A final article often consists of one or more polymers, multiple intentionally added substances, such as fillers or additives, as well as non-intentionally added substances, for instance residues from the manufacturing (*13*). Based on regulatory inventories, over 4000 substances are associated with plastic food packaging alone (*13*) and, overall, 10,547 chemicals are known to be used in plastics (*14*). Moreover, empirical data suggests that plastics contain more chemicals than currently known. Using nontarget chemical analysis, we detected hundreds to thousands of chemicals in plastic consumer products, most of these remaining unknown (*15*). Importantly, the totality of plastic chemicals in a product was toxic *in vitro*, inducing baseline toxicity, oxidative stress, cytotoxicity, and endocrine effects.

Building on these results and the fact that bisphenols and phthalates are known metabolism disrupting chemicals (MDCs; *8, 16, 17*), we hypothesized that MDCs are present in plastic consumer products and that metabolic disruption might represent a common but understudied toxicological property of plastic chemicals. To explore this, we used the same plastic consumer products we have extensively characterized previously (*15*), and investigated the extracts’ adipogenic activity in differentiation experiments with murine 3T3-L1 cells. Following exposure to MDCs, 3T3-L1 preadipocyte differentiate into adipocytes, accumulate triglycerides until they finally resemble mature white fat cells (*18*). The bioassay targets the induction of adipogenesis at the cellular level and represents a well-established *in vitro* model for metabolic disruption *in vivo* (*19*). We performed optimization experiments and applied high-content fluorescence microscopy combined with an automated image processing to increase the sensitivity and throughput. We also investigated the underlying mechanisms of the adipogenic response and tested whether the extracted plastic chemicals activate the human peroxisome proliferator-activated receptor gamma (PPARγ) or glucocorticoid receptor (GR). We selected PPARγ as a key regulator of adipogenesis (*20*), and included GR because glucocorticoids are important regulators of lipid metabolism (*21*). Accordingly, an excess of agonists for these receptors is associated with obesogenic effects in animal models and humans (e.g., weight gain; *19*). Moreover, we performed nontarget, ultra-high performance liquid chromatography coupled to a quadrupole of flight spectrometer (LC-QTOF-MS/MS) to characterize the chemicals present in plastics and compare these with a list of compounds known to induce adipogenesis.

## Results

### Adipogenic activity of plastic consumer products

To exclude cytotoxic effects masking the adipogenic response, we used nuclei count data to assess cytotoxicity. Most extracts were not cytotoxic up to the maximum concentration tested (3 mg plastic well^−1^), except for PP 4, PUR 3 and PUR 4. The latter two were the most cytotoxic samples with the highest noncytotoxic concentration (HNC) of 0.75 mg plastic well^−1^. The HNC for PP 4 was 1.5 mg plastic well^−1^ (Fig. S1). To assess the induction of adipogenesis by the plastic extracts, we present the numbers of adipocytes and mature adipocytes in the cell populations and the total lipid droplet count per image for the HNC of each sample. Data were compared to both vehicle and rosiglitazone–treated controls (Fig. 1). Further specifications on the endpoints of the automated image processing, dose-response relationships and example images can be found in the supplementary material (Fig. S2–S17).

**Fig. 1.**
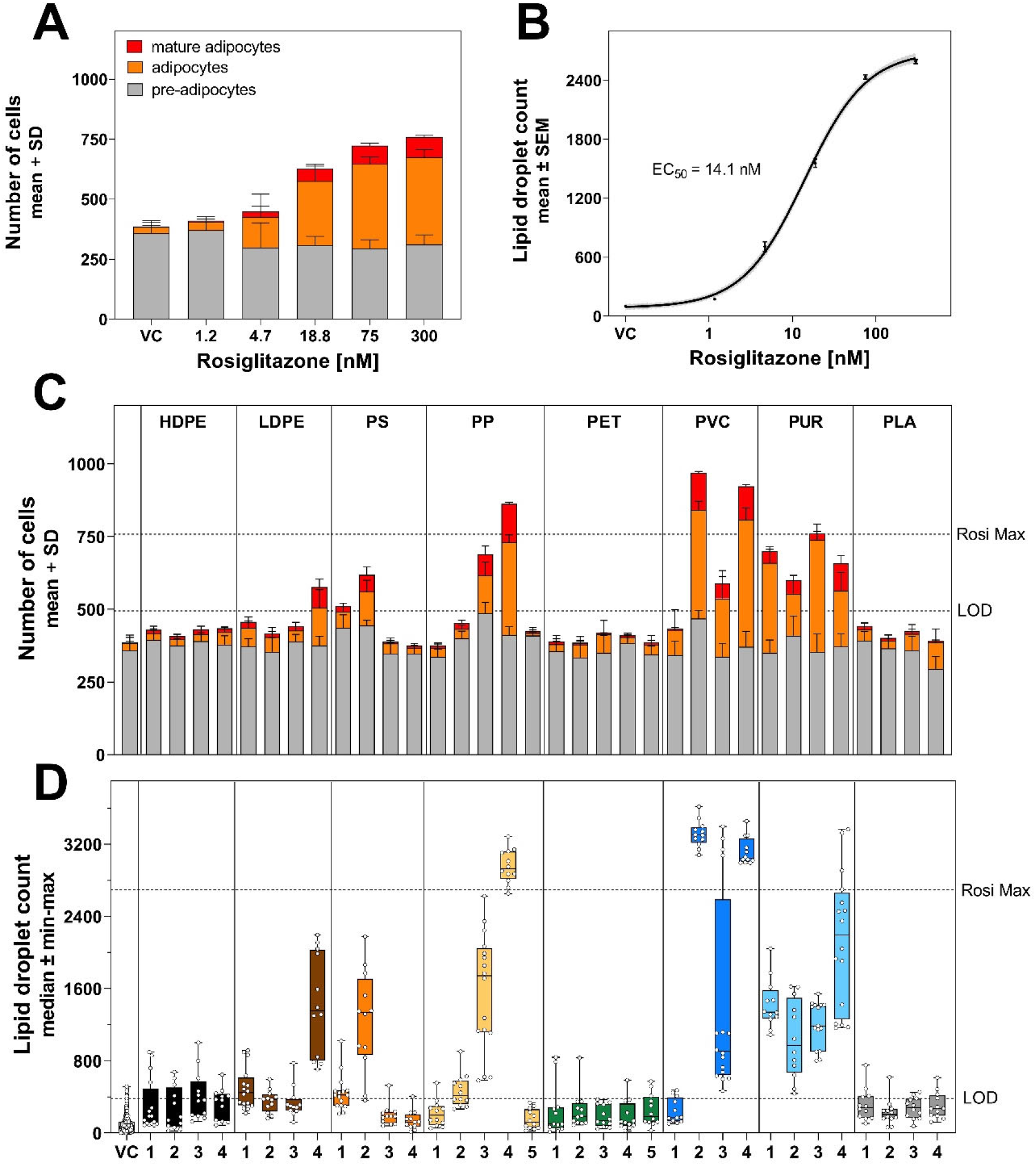
Effect of rosiglitazone on (A) the adipocyte population and (B) the lipid droplet count (pooled data from four experiments). Effect of plastic extracts on (C) the adipocyte population and (D) the lipid droplet count in the highest noncytotoxic concentration. The highest noncytotoxic concentration (HNC) was 3 mg plastics well^−1^ except for PP 4 (1.5 mg plastic well^−1^) as well as PUR 2 and PUR 3 (0.75 mg plastic well^−1^). VC = vehicle control, LOD = limit of the detection, Rosi Max = maximal response of rosiglitazone.

The extracts of eleven plastic consumer products induced adipogenesis with four samples having an equal or stronger effect than the maximal response of cells exposed to rosiglitazone (PVC 2 and 4, PP 4, PUR 3). Similarly to rosiglitazone (Fig. 1A), the proliferative effects induced by plastic extracts was driven by an increase in the numbers of adipocytes and mature adipocytes, while the number of preadipocytes remained stable (Fig. 1C). Regarding the polymer type, the most potent extracts were PUR and PVC, with seven out of eight samples inducing adipogenesis, whereas for PP, PS and LDPE only specific samples induced adipogenic responses. In contrast, PET, HDPE, and PLA samples were consistently inactive. The same pattern is reflected by the lipid droplet count data (Fig. 1D). Here, however, some additional samples induced a slight increase in lipid droplets (LDPE 1, PS 1, PP 2).

Given the propensity of environmental pollutants to promote unhealthy adipogenesis, we used the single-cell data to explore a size shift towards hypertrophy and increased accumulation of triglycerides in comparison to rosiglitazone (Fig. 2). We here present the results of one out of four experiments due to the large size of the dataset (results of the other experiments can be found in Fig. S18). Exposure to rosiglitazone dose-dependently increased the lipid content of adipocytes with a median lipid droplet area of 137 pixels cell^−1^ in the lowest concentration to 290 pixels cell^−1^ in the highest concentration (Fig. 2A). The average fluorescence intensity of the lipid droplets remained stable (Fig. 2B), thus the adipocytes increased in size in response to rosiglitazone but triglyceride accumulation within the droplets remained constant. Compared with the maximal response to rosiglitazone, adipocytes exposed to many of the active plastic extracts had a higher lipid content, resulting both from an increase in adipocyte size and from the amount of triglycerides contained within the lipid droplets. The lipid droplet area per adipocyte was greater in nine out of the eleven active samples with a median increase of 21.6–114% (PS 2–LDPE 4). In line with that, the average fluorescence intensity was higher in ten out of the eleven active samples with a median increase of 25.1–60.4% (PS 2–PVC 4). These effects are consistent over all experiments, except for PVC 3 (Fig. S18).

**Fig. 2.**
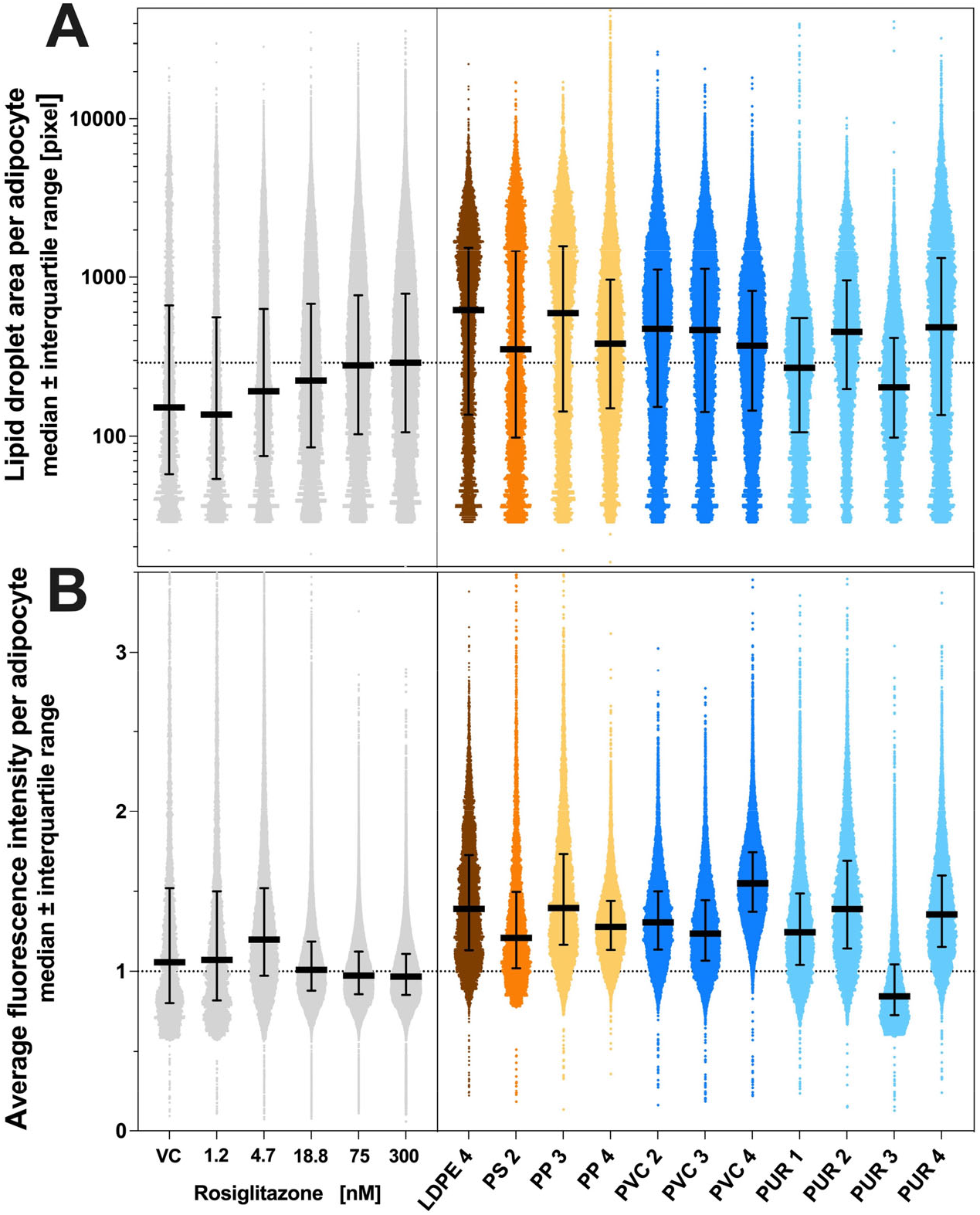
(A) Size distribution of adipocyte population and (B) accumulation of triglyceride per adipocyte in cells exposed to rosiglitazone (left) or the highest noncytotoxic concentration of the eleven active plastic extracts (right). Single-cell data from one experiment. Intensity data is normalized on the mean of the highest rosiglitazone concentration (300 nM). VC = vehicle control.

### Reporter gene assays

We observed a higher cytotoxicity of the extracts on the U2OS cells compared to the 3T3-L1 cells with five samples being cytotoxic. The most cytotoxic sample was PP 4 with a HNC of 0.19 mg plastic well^−1^, followed by PS 2, PP 3 and PLA 1 as well as PVC 2 with a HNC of 0.38 and 0.75 mg plastic well^−1^, respectively (Tab. S1).

None of the samples activated GR (Fig. S19) and five extracts activated PPARγ (Fig. 3A). PLA1 was the most potent sample with a median receptor activity of 34.7%, followed by PS 2 (24.4%), PVC 2 (10.3%), LDPE 2 (8.4%) and PVC 1 (7.3%). Accordingly, the PPARγ activity of plastic chemicals is a poor predictor of their adipogenic activity (Fig. 3B), except for PVC 2 and PS 2 which induced both PPARγ and adipogenesis at similar effect concentrations. Moreover, three out of the five samples activating PPARγ did not induce adipogenesis in 3T3-L1 cells (PLA 1, LDPE 2, PVC 1).

**Fig. 3.**
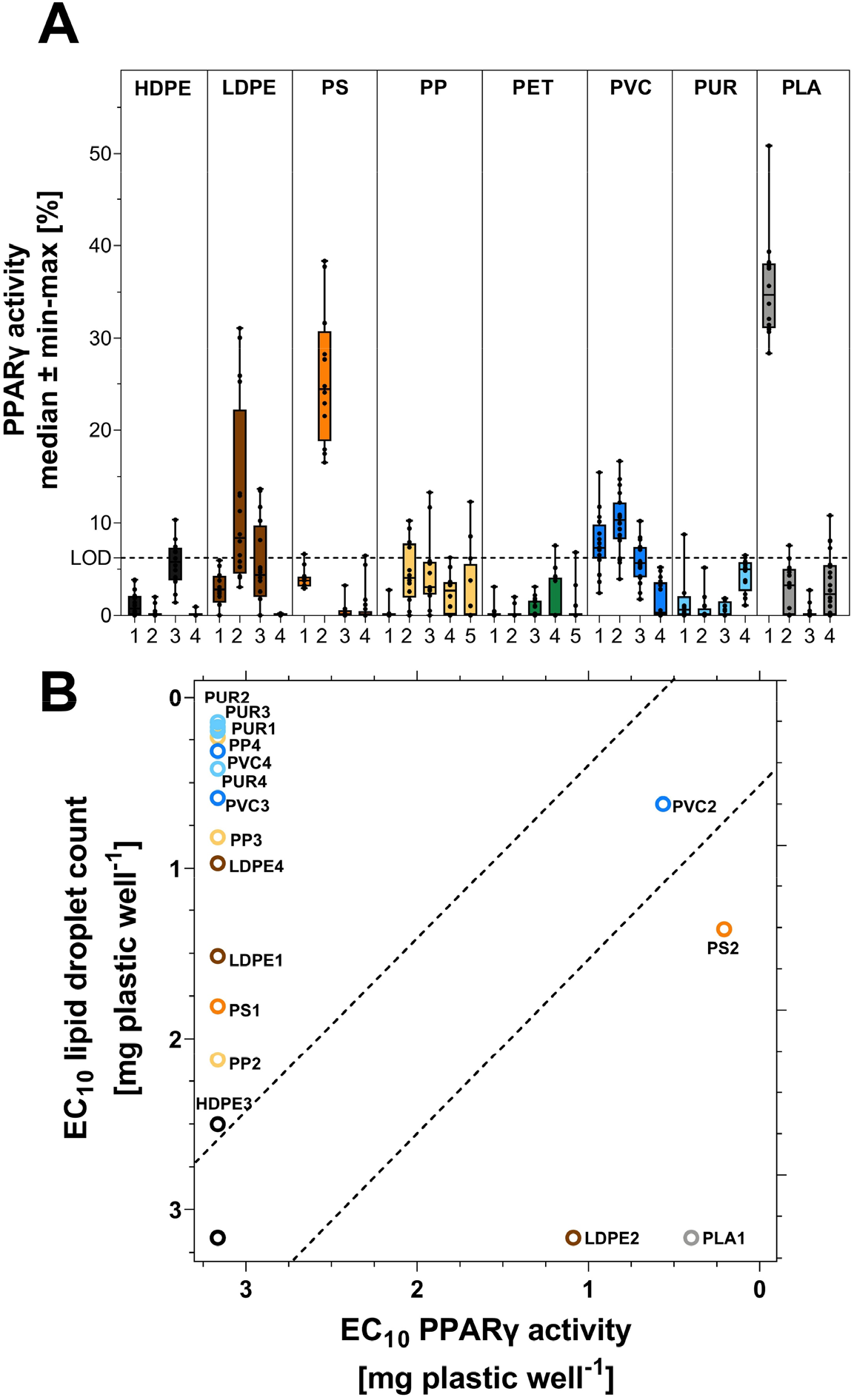
(A) PPARγ activity induced by plastic extracts at the highest noncytotoxic concentration and (B) correlation of the EC10 of the PPARγ activity and lipid droplet count. The highest noncytotoxic concentration (HCN) was 1.5 mg plastic well^−1^ except for PP 4 (0.19 mg plastic well^−1^), PS 2 and PP 3 (0.38 mg plastic well^−1^) as well as PLA 1 and PVC 2 (0.75 mg plastic well^−1^).

### Chemicals tentatively identified in plastics

Using nontarget GC-QTOF-MS/MS, we previously identified 260 unique chemicals in extracts of the same plastic products (*15*). This corresponds to 231 tentatively identified chemicals with 227 unique PubChem CIDs in the samples used in this study (Tab. S2). In the nontarget LC-QTOF-MS/MS analysis, we detected in total 55,300 features (i.e., unidentified chemicals) across all samples that were only present in samples or had a >10-fold higher abundance compared to the blanks. Here, the number of features in individual samples ranged from 107 (HDPE 2) to 6242 (PUR 3, Tab. 1). In total, 5500 features had spectral MS/MS data we could use for compound identification, out of which we detected between 30 (PS 4) and 2117 features (PUR 3) per sample.

**Table 1.**
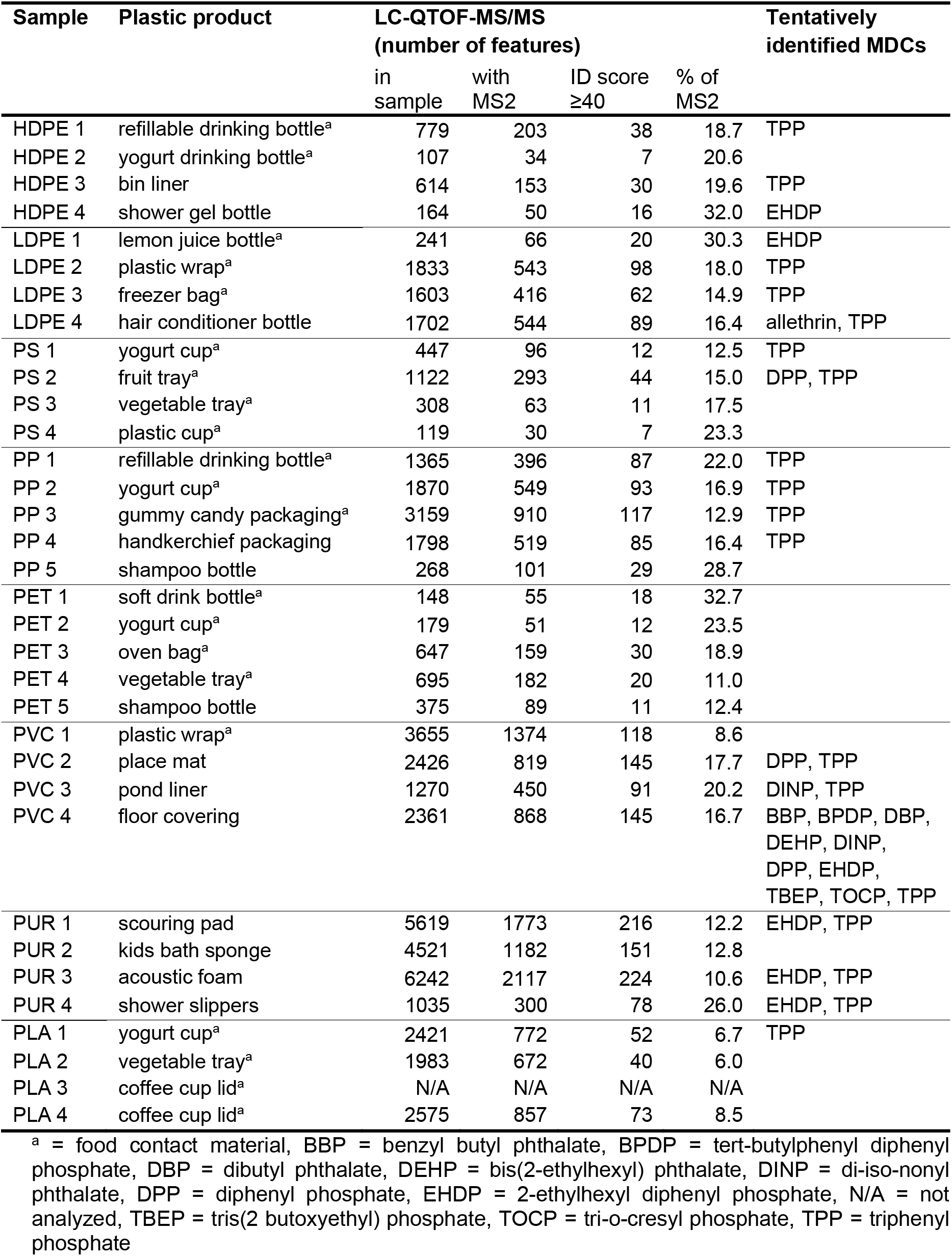
Plastic products analyzed in this study, results of the nontarget chemical analysis and the tentatively identified metabolic disrupting chemicals (MDCs).

| Sample | Plastic product | LC-QTOF-MS/MS<br>(number of features) |  |  |  | Tentatively<br>identified MDCs |
| --- | --- | --- | --- | --- | --- | --- |
|  |  | in<br>sample | with<br>MS2 | ID score<br>≥40 | % of<br>MS2 |  |
| HDPE 1 | refillable drinking bottle<sup>a</sup> | 779 | 203 | 38 | 18.7 | TPP |
| HDPE 2 | yogurt drinking bottle<sup>a</sup> | 107 | 34 | 7 | 20.6 |  |
| HDPE 3 | bin liner | 614 | 153 | 30 | 19.6 | TPP |
| HDPE 4 | shower gel bottle | 164 | 50 | 16 | 32.0 | EHDP |
| LDPE 1 | lemon juice bottle<sup>a</sup> | 241 | 66 | 20 | 30.3 | EHDP |
| LDPE 2 | plastic wrap<sup>a</sup> | 1833 | 543 | 98 | 18.0 | TPP |
| LDPE 3 | freezer bag<sup>a</sup> | 1603 | 416 | 62 | 14.9 | TPP |
| LDPE 4 | hair conditioner bottle | 1702 | 544 | 89 | 16.4 | allethrin, TPP |
| PS 1 | yogurt cup<sup>a</sup> | 447 | 96 | 12 | 12.5 | TPP |
| PS 2 | fruit tray<sup>a</sup> | 1122 | 293 | 44 | 15.0 | DPP, TPP |
| PS 3 | vegetable tray<sup>a</sup> | 308 | 63 | 11 | 17.5 |  |
| PS 4 | plastic cup<sup>a</sup> | 119 | 30 | 7 | 23.3 |  |
| PP 1 | refillable drinking bottle<sup>a</sup> | 1365 | 396 | 87 | 22.0 | TPP |
| PP 2 | yogurt cup<sup>a</sup> | 1870 | 549 | 93 | 16.9 | TPP |
| PP 3 | gummy candy packaging<sup>a</sup> | 3159 | 910 | 117 | 12.9 | TPP |
| PP 4 | handkerchief packaging | 1798 | 519 | 85 | 16.4 | TPP |
| PP 5 | shampoo bottle | 268 | 101 | 29 | 28.7 |  |
| PET 1 | soft drink bottle<sup>a</sup> | 148 | 55 | 18 | 32.7 |  |
| PET 2 | yogurt cup<sup>a</sup> | 179 | 51 | 12 | 23.5 |  |
| PET 3 | oven bag<sup>a</sup> | 647 | 159 | 30 | 18.9 |  |
| PET 4 | vegetable tray<sup>a</sup> | 695 | 182 | 20 | 11.0 |  |
| PET 5 | shampoo bottle | 375 | 89 | 11 | 12.4 |  |
| PVC 1 | plastic wrap<sup>a</sup> | 3655 | 1374 | 118 | 8.6 |  |
| PVC 2 | place mat | 2426 | 819 | 145 | 17.7 | DPP, TPP |
| PVC 3 | pond liner | 1270 | 450 | 91 | 20.2 | DINP, TPP |
| PVC 4 | floor covering | 2361 | 868 | 145 | 16.7 | BBP, BPDP, DBP,<br>DEHP, DINP,<br>DPP, EHDP,<br>TBEP, TOCP, TPP |
| PUR 1 | scouring pad | 5619 | 1773 | 216 | 12.2 | EHDP, TPP |
| PUR 2 | kids bath sponge | 4521 | 1182 | 151 | 12.8 |  |
| PUR 3 | acoustic foam | 6242 | 2117 | 224 | 10.6 | EHDP, TPP |
| PUR 4 | shower slippers | 1035 | 300 | 78 | 26.0 | EHDP, TPP |
| PLA 1 | yogurt cup<sup>a</sup> | 2421 | 772 | 52 | 6.7 | TPP |
| PLA 2 | vegetable tray<sup>a</sup> | 1983 | 672 | 40 | 6.0 |  |
| PLA 3 | coffee cup lid<sup>a</sup> | N/A | N/A | N/A | N/A |  |
| PLA 4 | coffee cup lid<sup>a</sup> | 2575 | 857 | 73 | 8.5 |  |

<sup>a</sup> = food contact material, BBP = benzyl butyl phthalate, BPDP = tert-butylphenyl diphenyl phosphate, DBP = dibutyl phthalate, DEHP = bis(2-ethylhexyl) phthalate, DINP = di-iso-nonyl phthalate, DPP = diphenyl phosphate, EHDP = 2-ethylhexyl diphenyl phosphate, N/A = not analyzed, TBEP = tris(2 butoxyethyl) phosphate, TOCP = tri-o-cresyl phosphate, TPP = triphenyl phosphate

For tentatively identifying the plastic chemicals, we used the MS/MS data with the MassBank library (14,788 compounds) and three *in silico* fragmented databases of chemicals potentially used in plastics or (pre-)registered for authorization on the European market (in total 75,510 compounds; *22*). These queries resulted in a successful identification of 2364 features across all samples, corresponding to 629 unique chemicals (SM Excel Tab. S1). Accordingly, between 6 (PLA 2) and 33 % (PET 1) of the features in each sample were tentatively identified. For the 25 compounds with the highest identification scores (≥ 50) and abundance in the samples, we confirmed the plausibility of the identification by checking whether the compounds are known to be used in plastics (SM Excel Tab. S2). We found that 14 out of 25 compounds are used in plastics, including five plasticizers (e.g., acetyl tributyl citrate), four flame retardants (e.g., tris(2-butoxyethyl) phosphate, tris(3-methylphenyl) phosphate) and multiple processing aids, such as the lubricant 2-nonyl-N-(2-nonylphenyl) aniline, the hardener 4-methylphthalic anhydride and the slip additive (Z)-docos-13-enamide. We also identified compounds that likely migrated from the packed content into the packaging (two octadecanamides used in cosmetics) and one compound that was implausible (the veterinary drug febarbamate).

When cross-referencing the chemicals tentatively identified in plastics and a list of known MDCs inducing adipogenesis in 3T3-L1 cells (Tab. S3), we found eleven compounds known to induce adipogenesis *in vitro*. The MDCs present in our samples include four phthalates and six organophosphates (Tab. 1). Benzyl butyl phthalate (BBP), di-butyl phthalate (DBP) and di(2-ethylhexyl) phthalate (DEHP) were present in PVC 4. Di-iso-nonyl phthalate (DINP) was detected in PVC 3 and 4. Diphenyl phosphate (DPP), 2-ethylhexyl diphenyl phosphate (EHDP) and triphenyl phosphate (TPP) were detected in multiple samples. When using raw abundance as a proxy for concentration, high levels of TPP, DPP and EHDP (the MDCs present in at least three samples) were detected in two to three active PVC samples (Tab. S4). In contrast, the other active samples contain very low levels of these chemicals (PS 2, PP 4, PUR 2, PUR 3). Interestingly, we did not detect organotin compounds or bisphenols (SM Excel Table S1) despite these are known MDCs and thought of as being common in PVC and other plastics (*16, 17*).

## Discussion

### Adipogenic activity of plastic consumer products

We hypothesized that plastic products contain MDCs based on the fact that a small set of compounds used in plastics is known to induce adipogenesis *in vitro* and *in vivo* (*8, 16, 17*). However, the current focus on few, individual plastic chemicals neglects the chemical complexity of plastic materials given that we know that thousands of compounds are either used in plastics or non-intentionally present (*13, 14*). Thus, we decided to characterize the adipogenic activity of all compounds extractable from plastic consumer products.

Eleven out of 34 products contain chemicals that induce adipogenesis and are, thus, MDCs *in vitro* (Fig. 1). The chemicals extracted from some plastics are not only quite potent but also trigger effects that are similar to or higher than those induced by the reference compound rosiglitazone (PVC 2 and 4, PP 4). Supramaximal efficacies have previously been reported for single compounds, such as dibutyl phthalate and tert-butyl phenyl diphenyl phosphate (*23*) but only at concentrations ≥10 μM, illustrating the potency of the extracted mixtures.

Products with multiple applications, including two FCMs (PS 2, PP 3) and nine non-FCMs, contain adipogenic chemicals. While chemicals migrating from packaging into food represent an obvious source of human exposure (*24*), compounds released from non-FCMs can also contribute via dermal uptake (e.g., PUR 4 shower slippers) or inhalation. For instance, dust contains chemical mixtures that induce adipogenesis (*23*). Here, we show that plastic flooring (e.g., PVC 4) contains MDCs that may contribute to human exposure if they partition into dust. Given the potency of the extracted mixtures and considering our close and constant contact with plastics, it appears plausible that plastic chemicals contribute to an obesogenic environment and, thus, the obesity pandemic.

The chemicals present in PVC and PUR products most consistently induce potent adipogenic responses, while compounds extracted from PET, HDPE, and PLA products were inactive. Apart from the PLA samples, this is in line with our previous findings for other toxicity endpoints (*15*). This suggests that PVC and PUR are more likely to contain MDCs compared to other polymers. However, the chemicals extracted from some PP, PS, and LDPE products also induce adipogenesis (Fig. 1). This supports the idea that caution is needed when trying to generalize the occurrence of toxic chemicals based on polymer type (*15*).

Unhealthy or dysfunctional adipocytes are part of the obesity phenotype. They are larger in size, have an impaired glucose uptake and insulin signaling, an elevated inflammatory response and decreased respiration (*25*). While we did not investigate the latter characteristics, adipocytes exposed to plastic chemicals were larger and contained more triglycerides compared to those treated with rosiglitazone (Fig. 2). Since rosiglitazone promotes the development of healthy white adipocytes (*26, 27*), these results suggest that exposure to plastic chemicals could shift adipocytes towards an unhealthy phenotype. Similar trends have been reported for a range of MDCs, including BPA (*28*), organotin compounds (*29, 30*) and DEHP (*31*), which we detected in PVC 4 (Tab. 1). Hence, it will be interesting to investigate whether plastic chemicals also trigger the other hallmarks of unhealthy, dysfunctional adipocytes.

### Plastic chemicals and adipogenesis

Using nontarget high resolution mass spectrometry, we show that plastic products contain hundreds to thousands of extractable chemicals; few of those identifiable using spectral libraries and *in silico* tools. This is in line with our previous research (*15, 22*) and points towards the presence of unknown chemicals in plastics (e.g., non-intentionally added substances). Accordingly, the relatively low identification performance in our study is a result of the limited coverage of chemical databases. These limitations notwithstanding, we tentatively identified a range of known plastic chemicals providing confidence in the accuracy of the identifications.

Plastic products contain known MDCs, including four phthalates (only in PVC 3 and 4) and six organophosphates (Tab. 1). Biomonitoring data suggests that humans are commonly exposed to some of these compounds (*32–34*). As an example, the phthalates DBP and DEHP as well as the flame retardants TPP and TBEP we found in plastics were recently detected in matched maternal and cord blood samples (*35*). Accordingly, plastic products can be one source of exposure to these MDCs.

Known MDCs may explain the adipogenic response to chemicals extracted from some but not all plastic samples. Most active samples contain at least one MDC with TPP, DPP and EHDP being present in multiple samples. Interestingly, the floor covering (PVC 4) contained ten known MDCs. While the active PVC samples contain high levels of TPP, DPP and EHDP, the abundance of these chemicals was very low in the other active samples (Tab. S4). This suggests that other than the known MDCs contribute to the adipogenesis induced by plastic chemicals.

### Underlying mechanisms

PPARγ is a key regulator of adipogenesis (*20*), and many MDCs that induce adipogenesis also activate PPARγ (*17*). Despite the common idea that PPARγ activation is a main mechanism via which anthropogenic chemicals trigger adipogenesis, most of the adipogenic plastic samples in fact do not activate this receptor (Fig. 3). Only in two cases (PVC 2, PS 2) does a high PPARγ activity correspond to a strong induction of lipid droplet formation. Moreover, three samples (PLA 1, LDPE 2, PVC 1) activated PPARγ but were inactive in the adipogenesis assay. Thus, the adipogenic effects of the plastic extracts is not necessarily dependent on direct activation of PPARγ and other mechanisms must be involved.

GR is another important nuclear receptor that participates in adipogenesis, and various MDCs activate GR (*36*). In particular, glucocorticoids are essential in inducing adipocyte differentiation (Fig. S20). However, none of the plastic extracts activated GR rendering this an unlikely mechanism of action in this case.

Elucidating the mechanism by which plastic chemicals induce adipogenesis is complex since we are dealing with two black boxes, namely the complex chemical mixtures present in plastics and the multitude of potential mechanism of actions involved in adipogenic responses in 3T3-L1 cells (*19*). In addition to PPARγ and GR, (ant)agonists of multiple other nuclear receptors, such as the retinoid x receptor α, estrogen receptor, androgen receptor, liver x receptor, thyroid receptor β, have been demonstrated or are discussed to contribute to an adipogenic response (*37*). In the light of the diversity of compounds we detected in plastics, it appears probable that these act via multiple mechanisms that are in most cases PPARγ and GR independent. Although more work needs to be done to elucidate the underlying mechanisms, our results underline the importance of using integrative methods, such as the adipogenesis assay to identify MDCs triggering cellular responses rather than assessing (anta)agonism at selected nuclear receptors.

### Limitations and future directions

To the best of our knowledge, this is the first study investigating the adipogenic activity of chemicals extractable from plastic consumer products. Considering the diversity of plastic products and their chemical composition, the sample set is certainly not representative of all plastic chemicals, humans will be exposed to. While it is challenging to comprehensively characterize the human exposure to plastic chemicals from all types of products, given their ubiquity and diversity, a way forward is to prioritize polymer types that are likely to contain MDCs, such as PVC and PUR.

Given that our aim was to investigate whether MDCs are present in plastic products, we used methanol to extract the samples. This simulates a worst-case scenario. Thus, even though we demonstrated that potent (mixtures of) MDCs are present in consumer products, it remains to be investigated whether these will migrate under more realistic conditions into air, water or food, or can be taken up dermally. Using the same samples as in the present study, we recently demonstrated that a significant number of chemicals and *in vitro* toxicity, such as antiandrogenic compounds, migrate into water (*22*). However, it remains unknown if this is also the case for the present MDCs.

Moreover, we analyzed plastic packaging that contained foodstuff or personal care products because we aimed at investigating final products. Since chemical migration is not a one-way street, we cannot exclude that compounds from the contents migrated into the packaging. The detection of compounds used in cosmetics in its packaging underlines this limitation. Such compounds may contribute to the observed adipogenic activity or PPARγ activation and future research should cover unused final packaging.

The nontarget chemical analysis resulted in the tentative identification of several MDCs. However, many compounds remain unidentified and there is some likelihood of false-positive identifications. The challenge of a rather low identification success is well-known for environmental pollutants (*38*) and can be addressed by building more comprehensive spectral databases. Recent efforts to build specific databases for plastic chemicals are promising (*13, 14*) but must be complemented with spectral information and non-intentionally added substances. Moreover, we show that known MDCs only partially – if at all – contribute to the adipogenesis induced by plastic extracts. This points towards the presence of unidentified MDCs in plastics. To identify the compounds that are indeed causative for the observed responses, future research should apply effect-directed analysis. Moreover, our results indicate that plastic chemicals may promote a development towards unhealthy adipocytes. However, more evidence is needed to further support this hypothesis. For instance, one needs to extend the adipogenesis assay to cover later stages of adipocyte development and investigate biomarkers of inflammation and metabolic function (e.g., glucose uptake, insulin sensitivity).

Taken together, we demonstrated that plastic consumer products contain potent (mixtures of) MDCs inducing adipogenesis *in vitro* via mechanisms that are mostly not mediated via PPARγ and GR. Accordingly, plastic chemicals may contribute to an obesogenic environment considering our constant contact with a broad range of plastic products. Given that the plastic products containing MDCs also contained compounds triggering other toxicological endpoints (*15*), a shift towards chemically less complex plastics represents a way forward to a non-toxic environment.

## Materials and Methods

A list of used chemicals is provided in the supplementary material (Tab. S5).

### Sample selection and plastic extraction

We used the same 34 plastic samples (Tab. 1) as in Zimmermann *et al.* (*15*). The samples cover petroleum-based polymers with the highest market share (polypropylene (PP) > low density polyethylene (LDPE) > high density polyethylene (HDPE) > polyvinyl chloride (PVC) > polyurethane (PUR) > polyethylene terephthalate (PET) > polystyrene (PS); *39*), and polylactic acid (PLA) as a bio-based alternative. The samples include 21 products with and 13 products without food contact. Further specifications on the sample selection, collection and polymer identification are described in Zimmermann *et al.* (*15*). We extracted 3 g sample with including three procedure blanks (PB 1-3) with methanol and concentrated the extracts to a final volume of 200 μL using dimethyl sulfoxide as a keeper. To contextualize the bioassay results, we use “plastic equivalents” in such that “1 mg plastic” corresponds to the chemicals extracted from 1 mg of plastic. Accordingly, 1 μl extract corresponds to 15 mg plastic. See supplementary text for details.

### Bioassays

We performed differentiation assays with murine 3T3-L1 adipocytes (Zenbio Inc., SP-L1-F, lot 3T3L1062104) to examine the induction of adipogenesis as well as CALUX reporter gene assays (BioDetection Systems B.V., Amsterdam, The Netherlands) to investigate the agonistic activity at the human peroxisome proliferator-activated receptor γ (*40*) and glucocorticoid receptor (GR, *41*). All experiments were conducted with negative controls, vehicle controls, positive controls, and PB 1–3. Samples, controls, and blanks were diluted 1000-fold (adipogenesis assay) or 500-fold (reporter gene assays) with medium, resulting in a maximum final solvent concentration of 0.1% or 0.2% (v/v), respectively. Each sample was analyzed in serial dilutions of 1:2 with four replicates per concentration in at least three independent experiments per assay. Moreover, the respective reference compound was included on every microtiter plate to control for potential variations between plates, and the sample arrangement was randomized to exclude position effects. As negative and vehicle controls did not differ significantly, the results from both were pooled. Further, there was no contamination during sample extraction and analysis since none of the controls and blanks induced activity (Fig. S21 and S22). See supplementary text for details on the cell culture conditions.

### Adipogenesis assay

We performed the differentiation assay with 3T3-L1 cells in accordance with a previously described method (*42*). In brief, an experiment consists of three days pre-differentiation (one day seeding, two days allowing cells to enter the resting state) followed by an 8-d differentiation window (two days differentiation, six days maintenance). Subconfluent cells of passage 10 were trypsinized and counted with a flow cytometer (NovoCyte, Acea Biosciences). 15,000 cells well^−1^ were seeded in 200 μL preadipocyte medium (PAM: DMEM-high supplemented with 10% bovine calf serum and 1% penicillin/streptomycin) into 96-well black, clear bottom tissue culture plates (655090, Greiner Bio-One) and incubated at 37 ℃ and 5% CO_2_. After 24 h, we checked that the cells reached confluency, replaced the medium with 200 μL fresh PAM well^−1^, and cultured the cells for another 48 h to initiate growth arrest. We included on every plate a pre-adipocyte control (undifferentiated cells) which was kept in PAM, while the rest of the cells were differentiate as described below.

### Optimization experiments

Given that a systematic analysis of dexamethasone (DEX) effects on triglyceride accumulation and differentiation efficiency in 3T3-L1 cells was lacking, we conducted optimization experiments to identify a suitable DEX concentration to initiate adipocyte differentiation that results in the lowest baseline as well as the highest sensitivity and dynamic range when co-exposed to the reference compound rosiglitazone. Moreover, we compared two methods to quantify triglycerides based on Nile Red Staining. We determined the total NileRed fluorescence well^−1^ and compared it to an automated imaging and analysis platform to determine whether the latter improves the sensitivity and dynamic range for the screening of adipogenic activity.

Based on the results (Fig. S20) and in comparison with previous studies, we found that a rather low optimal DEX concentration (6.25 nM) was sufficient to initiate adipocyte differentiation without increasing the assay’s baseline. Compared to the fluorescence readout well^−1^, an automated imaging approach is more sensitive to assess proliferation, enhances the dynamic range of the assay (Fig. S20) and provides more information because it enables single-cell analysis for a more in-depth characterization of the adipocyte population (pre-adipocytes, adipocytes and mature adipocytes). Accordingly, we analyzed the effects of the plastic extracts using 6.25 nM DEX during the differentiation window and the automated imaging approach. See supplementary text for details.

### Dosing of samples

To initiate differentiation, we replaced the PAM medium with 200 μL differentiation medium well^−1^ (DM: DMEM-high supplemented with 10% FBS, 1% penicillin/streptomycin, 20 mM HEPES, 1 μg mL^−1^ insulin, 0.5 mM 3-isobutyl-1-methylxanthine (IBMX), and 6.25 nM DEX) containing 5 concentrations of samples serially diluted 1:2 (0.19–3 mg plastic well^−1^ equivalent to 0.94–15 mg plastic mL^−1^) or rosiglitazone (1.17–300 nM). Following the 48-h differentiation, the medium was replaced with 200 μL adipocyte maintenance medium well^−1^ (DM without IBMX and DEX) containing the respective controls, samples, or rosiglitazone and changed the medium every other day during the 6d maintenance period.

### Fixation and staining

After 11 d, cells were fixed with 2% paraformaldehyde and co-stained with NileRed and NucBlue. Imaging was carried out on the Cytation 5 Cell Imaging Multimode reader (BioTek). Three images per field (Brightfield, NucBlue and NileRed) and nine fields per well were captured. See supplementary text for details.

### Image analysis

Images were analyzed in the open-source software CellProfiler (*43*). A description of the image analysis protocol and the CellProfiler pipelines are available in the supplementary material. We quantified proliferation (nuclei count), number of lipid droplets (lipid droplet count), total area occupied by lipid droplets (total area), and the total intensity of the NileRed staining within the lipid droplets (total intensity) per image. We further quantified the total area occupied by lipid (lipid droplet area), and the total intensity of NileRed staining in the lipid droplet area per cell.

Analysis of the single-cell data was applied to quantify the numbers of adipocytes and mature adipocytes per image. An adipocyte was defined as a cell containing at least one lipid droplet and a mature adipocyte was defined as a cell containing a total lipid content equivalent to ≥ 8 average sized lipid droplets (1000 pixels).

Single-cell measurements of lipid droplet area, and NileRed staining were analyzed further to explore how triglyceride accumulation was distributed within an adipocyte population. To control for potential cross-plate differences in staining intensity within an experiment, the average fluorescence intensity per adipocyte was normalized to the mean average fluorescence intensity for an internal plate control (cells treated with 300 nM rosiglitazone). For an independent experiment, single-cell data were grouped based on the treatment. The median lipid droplet area and normalized average fluorescence intensities per cell were calculated for each experiment. An example of the images and visualized output of the image analysis is shown in Fig. S2.

### Reporter gene assays

We performed the CALUX reporter gene assays in 384-well plates and used imaging to count nuclei per well to normalize the reporter gene response and assess cytotoxicity. Rosiglitazone was the reference compound for PPARγ and DEX for GR (Fig. S23). See supplementary text for the detailed protocol.

### Analysis of bioassay data

We used GraphPad Prism 9 (GraphPad Software, San Diego, CA) for non-linear regressions and statistical analysis, and interpolated plastic equivalents inducing 10 or 20% effect (effect concentration, EC_10_ or EC_20_) from the respective dose-response curves. The limit of detection (LOD) of each endpoint and experiment was calculated as three times the standard deviation (SD) of pooled controls. See supplementary text for details.

### Nontarget chemical analysis

We analyzed all samples, except PLA 3, using ultra-high performance liquid chromatography coupled to a quadrupole time of flight spectrometer (LC-QTOF-MS/MS) with an Acquity UPLC Waters liquid chromatography system coupled to a SYNAPT G2-S mass spectrometer (both Waters Norge, Oslo, Norway). The analytical method has been described in Zimmermann *et al.* (*22, 44*) and a brief description as well as information about the data analysis and compound identification can be found in the supplementary text.

### Comparison with chemicals known to induce adipogenesis

We built a list of 120 know adipogenic chemicals (Tab. S3) by searching Web of Science (Core Collection) for studies investigating chemicals in the adipogenesis assay and complemented the search with chemicals reviewed by Amato *et al.* (*16*). We cross-referenced the list with the tentatively identified compounds in the plastic samples to determine whether some of these compounds are MDCs (Tab. 1). See supplementary text for details.

## Acknowledgments

This study was support by internal funding of the Norwegian University of Science and Technology (NTNU, Trondheim). We thank Susana Villa Gonzales (NTNU) for UPLC-QTOF-MS/MS training and support.

## Author contributions

J.V., F.A. and M.W. conceived the study, J.V. and Å.V. performed the experiments, L.Z. performed the chemical analysis, J.V., F.A., and M.W. analyzed the data and wrote the manuscript, and all authors provided comments on the manuscript.

## Data and materials availability

The raw mass spectral data can be accessed under DOI 10.5281/zenodo.4781257 (published after publication), bioassay data (SM Excel Tab. S3–10) and the CellProfiler pipelines can be found in the supplementary material.

