## Supplementary material for "Adipogenic activity of chemicals used in plastic consumer products"

*Johannes Völker

**This PDF file includes:**

Supplementary Materials and Methods
Figures S1 to S24
Tables S1 to S5
Legends for Datasets S1

**Other supplementary materials for this manuscript include the following:**

Datasets S1
CellProfiler pipelines (Adipogenesis_assay.cpproj, Nuclear_counts.cpproj)

### Supplementary Materials and Methods

**Plastic extraction.** We used the original samples stored in glassware, except for PLA 3. As there was not enough material available for the old PLA 3 sample and we were not able to obtain more of the same product, we replaced the sample with a PLA coffee lid sample. To avoid sample contamination, glass or polytetrafluorethylene consumables were used for the sample extraction and all material was rinsed twice with acetone and annealed at 200 °C for ≥ 3 h. The samples were cut into 0.5–0.8 × 2 cm pieces. Foamy products were cut to a thickness of 0.5 cm. We weighed three grams each into 1 or 2 transparent glass vials depending on the sample volume, added 20 mL methanol (≥ 99.9%, Sigma Aldrich), and extracted the samples by sonication in an ultrasonic bath for 1 h at room temperature. We then transferred the methanol into clean glass vials, added 200 µL dimethyl sulfoxide (≥ 99.5%, Sigma Aldrich) as a keeper and evaporated the samples under a gentle stream of nitrogen to a final volume of 200 µL. Further, we treated three procedural blanks (PB 1–3) not containing any sample identically to control for contamination and stored the final extracts at -20 °C prior to analysis.

**Cell culture conditions**. 3T3-L1 cells were cultured in preadipocyte medium (PAM: DMEM-high supplemented with 10% bovine calf serum and 1% penicillin/streptomycin). Culturing 3T3-L1 cells over multiple passages can cause a decline in differentiation efficiency due to the loss of contact inhibition (*45*). Thus, cryo-cultures of passage 9 were thawed, subcultured once upon reaching 60-80% confluency, and sub-confluent cells of passage 10 were used for all experiments to ensure comparability and preserve differentiation capability. CALUX cells were maintained in growth medium (DMEM/F-12 supplemented with 7.5% fetal bovine serum, 1% penicillin/streptomycin, and 0.2 mg $mL^{-1}$ G418 and non-essential amino acids) and subcultured twice weekly and used until passage 20.

**Optimization of the adipogenesis assay.** We conducted optimization experiments to identify a suitable concentration of dexamethasone (DEX) to initiate adipocyte differentiation. Moreover, we applied high-content fluorescence imaging combined with an automated image processing in addition to the fluorescence readout well$^{-1}$ at the end of the experiment and compared both methods with regards to sensitivity.

Following the growth arrest window, the medium was replaced with 200 µL differentiation medium well$^{-1}$ (DM: DMEM-high supplemented with 10% FBS, 1% penicillin/streptomycin, 20 mM HEPES, 1 µg mL$^{-1}$ insulin and 0.5 mM 3-isobutyl-1-methylxanthine (IBMX)) containing either none or six concentrations of dexamethasone (6.25–250 nM) and eight concentrations of the reference compound rosiglitazone (300 pM – 1 µM). After the 48-h differentiation window, we replaced the medium with 200 µL adipocyte maintenance medium well$^{-1}$ (DM without IBMX and DEX) containing the eight rosiglitazone concentrations and changed the medium every other day during the maintenance period.

Counting nuclei based on NucBlue staining using the imaging approach was more sensitive than the standard fluorescence readout for detecting the proliferative effect of rosiglitazone (Fig. S20 A) with an $EC_{50}$ of 16.3 and 41.2 nM for nuclei counts and fluorescence readout, respectively. In contrast, quantification of adipogenesis had comparable sensitivity (Fig. S20 B) with an $EC_{50}$ of 10.6 and 11.4 nM for lipid droplet intensity and total NileRed fluorescence readout, respectively. However, the dynamic range of the assay was greatly enhanced using the imaging approach with an 11.7-fold increase in the lipid droplet count at the highest rosiglitazone concentration versus 4.69-fold increase in the fluorescence readout. In addition, the imaging-based approach provides more information including the characterization of the differentiation stage of cells in the population, and single-cell measurements to quantify the size of adipocytes and triglyceride accumulation (e.g., number and size of lipid droplets). Thus, high-content imaging with automated image processing can greatly extend our capabilities fir screening of MDCs *in vitro*.

Glucocorticoids in the differentiation medium are essential to prime the preadipocytes for adipogenesis and the differentiation success is variable and weak when DEX is absent (Fig. S24). In contrast, an excess of DEX (>25 nM) results in a significant stimulation of adipogenesis without an additional inducer and, thus, reduces the capability of the assay to detect adipogenic responses (Fig. S20 C). This is in line with a previous study reporting up to 40% of adipocytes in the vehicle controls using 250 nM DEX in the differentiation medium (*46*). Accordingly, the use of DEX concentrations varying from 0 (*36, 42, 47*) up to 1 µM (*45, 46, 48, 49*) might contribute to the poor reproducibility and comparability of 3T3-L1 studies. Based on our experiments, we recommend using a rather low DEX concentration of 6.25 nM which was sufficient to initiate adipocyte differentiation without increasing the assay baseline (Fig. S20 D).

Based on these results, we analyzed the effects of the plastic extracts in using 6.25 nM DEX during the differentiation window and the automated imaging approach.

**Fixation and staining**. After 11 d, the medium was removed, and cells were rinsed with PBS and fixed with 2% paraformaldehyde for 10 min on ice. The fixative was removed, and cells were rinsed twice with PBS and stored at 4 ℃ prior to staining. Cells were co-stained with 100 µL NileRed solution $well^{-1}$ (19.5 mL PBS + 500 µL AdipoRed (N3013, Lonza) and 1 drop $mL^{-1}$ NucBlue (R37605, Thermo, Hoechst 3342 staining)). Plates were incubated for 40 min in the dark at room temperature. Stained cells were washed twice with PBS and stored at 4 ℃ prior to analysis.

Fluorescence per well was measured using a Cytation 5 Cell Imaging Multimode reader (BioTek with excitation at 485 nm and emission at 572 nm for NileRed, and excitation at 360 nm and emission at 460 nm for NucBlue). Imaging was carried out on the same instrument using a 10× Plan Fluorite objective (WD10, NA 0.3). Image-based autofocusing of NucBlue fluorescence was used to select the image plane, and three images per field were captured (Brightfield, NucBlue and NileRed). A 365 LED with DAPI filter cube (Ex 377/50, Em 447/60) was used to detect the NucBlue staining, and a 523 LED with RFP filter cube (Ex 531/40, Em 593/40) for NileRed. Nine fields were captured per well.

**Cell profiler analysis**. For the adipogenesis assay, NucBlue and NileRed staining imaged at x10 magnification were analyzed using the following protocol to generate the assay measurements described.

*1. Cell identification*: nuclei (primary objects) were identified using an Otsu thresholding method based on NucBlue staining and used as the seed objects to identify cells (secondary objects). NileRed images were smoothed by gaussian filtration and used to guide the propagation algorithm for secondary object identification with a minimal threshold factor to limit the foreground.

*2. Lipid droplet identification* (adapted from Adomshick *et al*. (*50*)): lipid droplets were identified using a minimum cross entropy thresholding method applied to the NileRed images followed by a filtration step based on the mean intensity per droplet.

*3. Image-based measurements:* the number of cells and the number of lipid droplets were counted in each image. We additionally measured the total area occupied by lipid droplets, and the intensity of the Nile Red staining in this region.

*4. Single cell analysis:* to measure the lipid content per cell, lipid droplets were assigned to a given parent and merged such that the total area occupied by lipid, and the average intensity of the NileRed staining in this region could be calculated.

5. *Data processing:* filtration steps were subsequently applied to identify adipocytes (any cell containing at least one lipid droplet), and mature adipocytes (a cell having a lipid droplet area ≥ 1000 pixels, equivalent to ≥ 8 average size lipid droplets.)

For the nuclear counts (reporter gene assays), NucBlue staining imaged at x4 magnification was analyzed using the following protocol to quantify the number of nuclei in a given field. Nuclear counts were used for normalization and calculation of cytotoxicity.

*1. Image correction:* we applied the background method, with smoothing based on a gaussian filter to calculate an illumination function which was applied to NucBlue images to correct for the uneven illumination resulting from imaging the 384 well plates at ×4 magnification.

*2. Identification of nuclei*: nuclei were identified using an Otsu thresholding method based on the corrected images and we filtered the resulting objects based on their shape (form factor) to obtain final nuclear counts.

The cell profiler pipelines are attached:

- Adipogenesis_assay.cpproj
- Nuclear_counts.cpproj

**Reporter gene assays**. We performed the CALUX reporter gene assays in white clear polystyrene CellStar 384-well plates (781098, Greiner Bio-One). Trypsinized cells were resuspended in assay medium (DMEM/F-12 without phenol red supplemented with 5% charcoal-stripped FBS, 1% penicillin/streptomycin, non-essential amino acids). 3000 cells well$^{-1}$ were seeded in 25 µL and plates were incubated at 37 ℃ and 5% $CO_2$. Samples and reference compounds were prepared in assay medium (2-fold higher than the final assay concentration) in six concentrations per sample serially diluted 1:2 or eight concentrations of the reference compound (rosiglitazone for PPAR$\gamma$ and dexamethasone for GR; Fig. S23). After 24 h of incubation, 25 µL sample was added to the 25 µL assay medium well$^{-1}$ (1-fold), resulting in final sample concentrations of 0.05–1.5 mg plastic well$^{-1}$ (equivalent to 0.09–30 mg plastic mL$^{-1}$). After 23 h of exposure, the medium was replaced with 25 µL NucBlue staining solution well$^{-1}$ (1 drop NucBlue per mL PBS, Thermo Fisher Scientific) and incubated for 30 min in the dark at room temperature. Imaging was performed on the Cytation 5 Cell Imaging Multimode reader (BioTek) with a 4× Plan Fluorite objective (WD 17 NA 0.13) using a 365 LED with DAPI filter cube (Ex 377/50, Em 447/60) to detect NucBlue staining. A single field was captured per well. Following the imaging, a white sticker was placed on the transparent bottom of the plates, and the staining solution was replaced with 20 µL cell lysis buffer (25 mM pH 7.8 TRIS, 2 mM DDT, 2 mM CDTA, 10% glycerol and 1% Tritron-X100), and cells were lysed by linear shaking for 3 min. Luminescence was measured (Cytation 5) for one second after injection of 30 µL illuminate mix (20 mM Tricine, 1.07 mM $C_4H_2Mg_5O_{14}$, 2.67 mM $MgSO_4$ ´ $7H_20$, 0.1 mM EDTA, 1.5 mM DDT, 539 µM D-Luciferine, 5.49 mM ATP) followed by quenching of the reaction with 30 µL 0.1 M NaOH.

**Analysis of bioassay data**. We used GraphPad Prism 9 (GraphPad Software, San Diego, CA) for non-linear regressions and statistical analysis. To express cytotoxicity, we normalized the nuclei count to the vehicle controls (0% cytotoxicity) and a value of zero (100% cytotoxicity). We used 20% as cytotoxicity threshold. When the value of an individual image was over 20%, all channels of that image were excluded from further analysis (cytotoxic or out of focus). If the mean value of a replicate exceeded 20%, the replicate was excluded. When more than one replicate per concentration exceeded the threshold, the concentration was defined as cytotoxic. Fluorescence and luminescence readouts were corrected for background (well without cells). Percentage increase (proliferative effects) or fold induction over the corresponding vehicle control were calculated for each endpoint of the adipogenesis assay to compare both methods in the optimization experiments. To express agonistic activity in the reporter gene assays, luminescence data were normalized to the maximal assay response (100% activity: upper plateau of the dose-response relationship) of the corresponding reference compound and the mean value of the vehicle control (0% activity). The limit of detection (LOD) of each endpoint and experiment was calculated as three times the standard deviation (SD) of pooled controls. Dose-response relationships for all investigated endpoints were calculated using a four-parameter logistic function constrained to the bottom level of zero (0% activity). The respective plastic equivalents inducing 10 or 20% effect (effect concentration, $EC_{10,}$ or $EC_{20}$) were interpolated from the dose-response curves.

**Nontarget chemical analysis**. We analyzed all samples, except PLA 3, using ultra-high performance liquid chromatography coupled to a quadrupole time of flight spectrometer (LC-QTOF-MS/MS) with an Acquity UPLC Waters liquid chromatography system coupled to a SYNAPT G2-S mass spectrometer (both Waters Norge, Oslo, Norway). The analytical method has been described in (*22*) and (*44*). In brief, we injected 2 µL sample, equivalent to the chemicals extracted from 1.5 mg plastic, and performed the chromatographic separation on an Acquity UPLC BEH C18 column equipped with a C18 guard column (both from Waters). The mass spectrometer equipped with an electron-spray ionization source was operated in positive ionization mode with a mass range of 50-1200 Da at a resolution of 20,000. MS data were recorded from 2–35.5 min with a data-dependent acquisition (triggered when the threshold of an individual ion intensity exceeded 25,000 counts, maximum 15 precursor ions per survey scan) using a collision energy ramp (8–35 eV in the low mass region and 30–70 eV in the high mass region). After every $7^{th}$–$8^{th}$ sample, we analyzed a solvent blank (mobile phase, methanol or DMSO, n = 15) and a quality control sample (containing an aliquot of each sample). Two procedural blanks from the extraction were also analyzed. The raw mass spectral data can be accessed under DOI 10.5281/zenodo.4781257 (published after publication).

**Chemical data analysis and compound identification**. We imported the data for the 15 blanks, two PBs, six quality controls and 33 samples to Progenesis QI (version 3.0, Nonlinear Dynamics) and corrected for the lock mass of leucine enkephalin. We automatically aligned the retention times of all blanks and samples using the quality controls. Peak picking was performed on the samples using common adducts (M+H, M+2H, M+H-H2O, M+H-2$H_2O$, 2M+H, M+Na, M+2Na, M+H+Na, M+2H+Na, M+2Na+H, M+2Na-H), an automatic sensitivity, a minimum peak width of 0.02 min and a fragment sensitivity of 0.2% of the base peak.

We generated a list of chemical features which had MS/MS data and performed the further data analysis as described before (*22*). Basically, we filtered for features that were not detected in the solvent and procedural blanks or present in the samples with an at least 10-fold higher raw abundance than the maximum abundance of that feature in any of the blanks.

To tentatively identify the remaining features, we compared their mass spectra with the empirical spectra in MassBank (14,788 compounds, release version 2021.03, https://github.com/MassBank/MassBank-data/releases/tag/2021.03) and with three databases covering chemicals present in plastic packaging (2680 compounds), registered under the REACH regulation in 2020 (7092 compounds) and (pre)registered under REACH in 2017 (65,738 compounds) using the Metascope algorithm in Progenesis QI with a precursor tolerance of 5 ppm and a fragment tolerance of 10 ppm. The compounds in the latter three databases were *in silico* fragmented using the Metascope algorithm. The resulting identification was accepted if the match score was > 40. When a feature had multiple identifications with a score > 40, the first hit with the highest score was accepted. The identification corresponds to confidence level 3 according to Schymanski *et al.* (*51*).

**Comparison with chemicals known to induce adipogenesis**. We built a list of know adipogenic chemicals by searching Web of Science (Core Collection) for studies investigating chemicals in the adipogenesis assay using the following search strings: “(3T3L1 OR 3T3-L1) AND toxic* AND chemical*” (58 hits) as well as “(3T3L1 OR 3T3-L1) AND obesogen* OR metabolic disruptor* AND in vitro” (241 hits). The search was conducted on March 22, 2021. We removed duplicates and reviews and screened the remaining 254 full text articles for studies that investigated the adipogenic activity of chemicals in 3T3-L1 adipocytes. We decided not to perform a quality assessment to keep the list broad and included from 47 suitable studies all chemicals which were reported to be adipogenic. We further complemented the list with the chemicals reviewed by Amato *et al.* (*16*) and ended up with a list of 120 adipogenic chemicals (Tab. S3). For comparison with our results, we added the associated PubChem CIDs.

To cross-reference this list with the compounds we tentatively identified in plastics, we built a joint compound list based on our previous GC-QTOF-MS/MS analysis (*15*) and the present LC-QTOF-MS/MS analysis. For the former, we translated the available CAS numbers of all tentatively identified compounds to SMILES using the US EPA’s CompTox Dashboard (https://comptox.epa.gov/dashboard) and then translated the SMILES to PubChem CIDs using the PubChem Identifier Exchange Service (https://pubchem.ncbi.nlm.nih.gov/idexchange). If CAS numbers were invalid or unavailable, we searched the compound name in PubChem and manually annotated the CID. For the LC-QTOF-MS/MS data, we used the PubChem CIDs provided by Progenesis QI or manually annotated the compound names provided by MassBank. The combined list from the GC- and LC-QFOT/MS/MS data contained 803 unique chemicals with CIDs (Tab. S2 and Excel Tab. S1). To determine whether some of these compounds are MDCs, we cross-referenced both CID lists (Tab. 1).

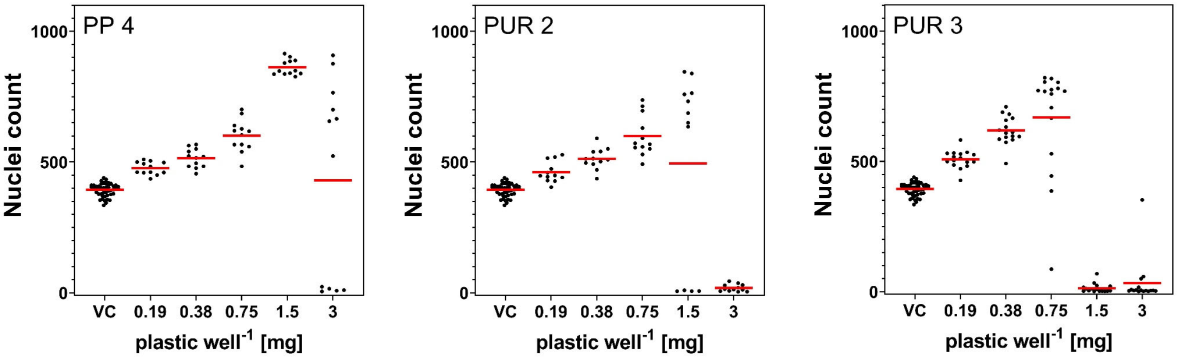

**Fig. S1. Nuclei count of the cytotoxic plastic extracts (PP 4, PUR 2, PUR 3) in the adipogenesis assay.** Data is presented as mean count per field (red lines) from three to four independent experiments performed with four replicates each (dots, n ≥ 12). VC = vehicle control.

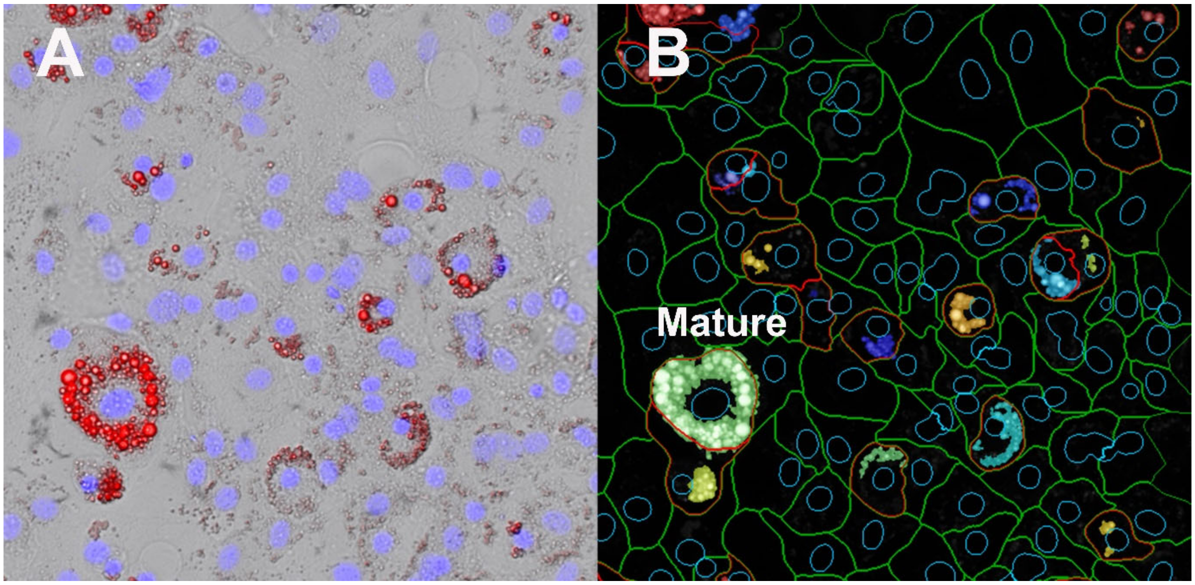

**Fig. S2. Image analysis example.** Differentiated 3T3-L1 cells exposed to rosiglitazone (4.69 nM). (A) Merged brightfield and fluorescence images. Nuclei are stained with NucBlue (blue) and lipid with NileRed (red). (B) Corresponding object identification performed with CellProfiler. Nuclei are outlined in blue, cell boundaries in green, and adipocytes in red. Identified lipid droplets are shown as a solid color and all lipid droplets associated with a given adipocyte are displayed in the same color.

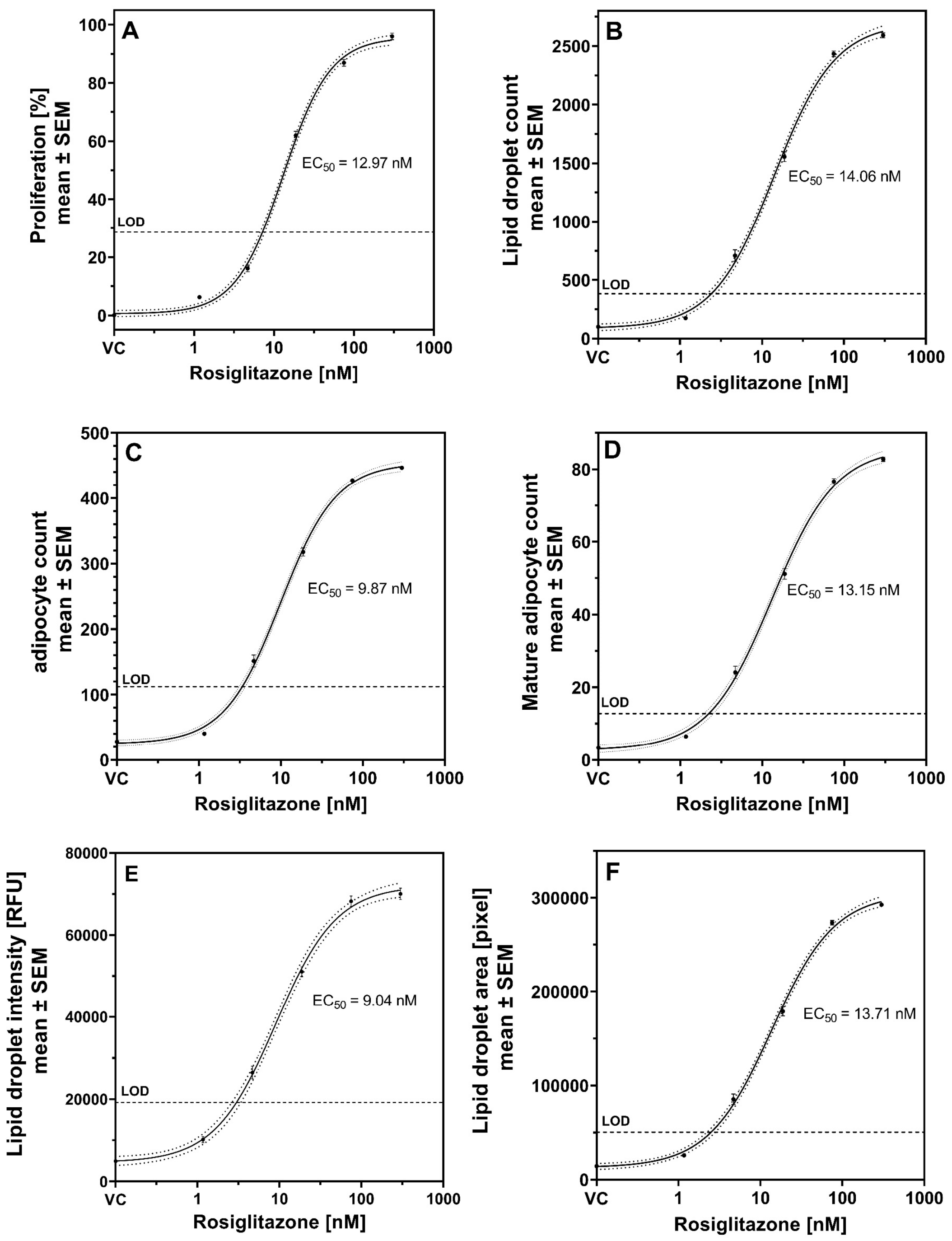

**Fig. S3. Dose-response relationship for the reference compound rosiglitazone in the adipogenesis assay with 6.25 nM dexamethasone in the differentiation medium.** (A) proliferation normalized on the mean of the vehicle control, (B) lipid droplet count per field, (C) adipocyte count per field, (D) mature adipocyte count per field, (E) total intensity of the NileRed staining within the lipid droplet mask per field and (F) total area occupied by lipid droplets per field. 160 or more replicates per concentration (n ≥ 160). VC = vehicle control, LOD = limit of detection, RFU = relative fluorescence unit.

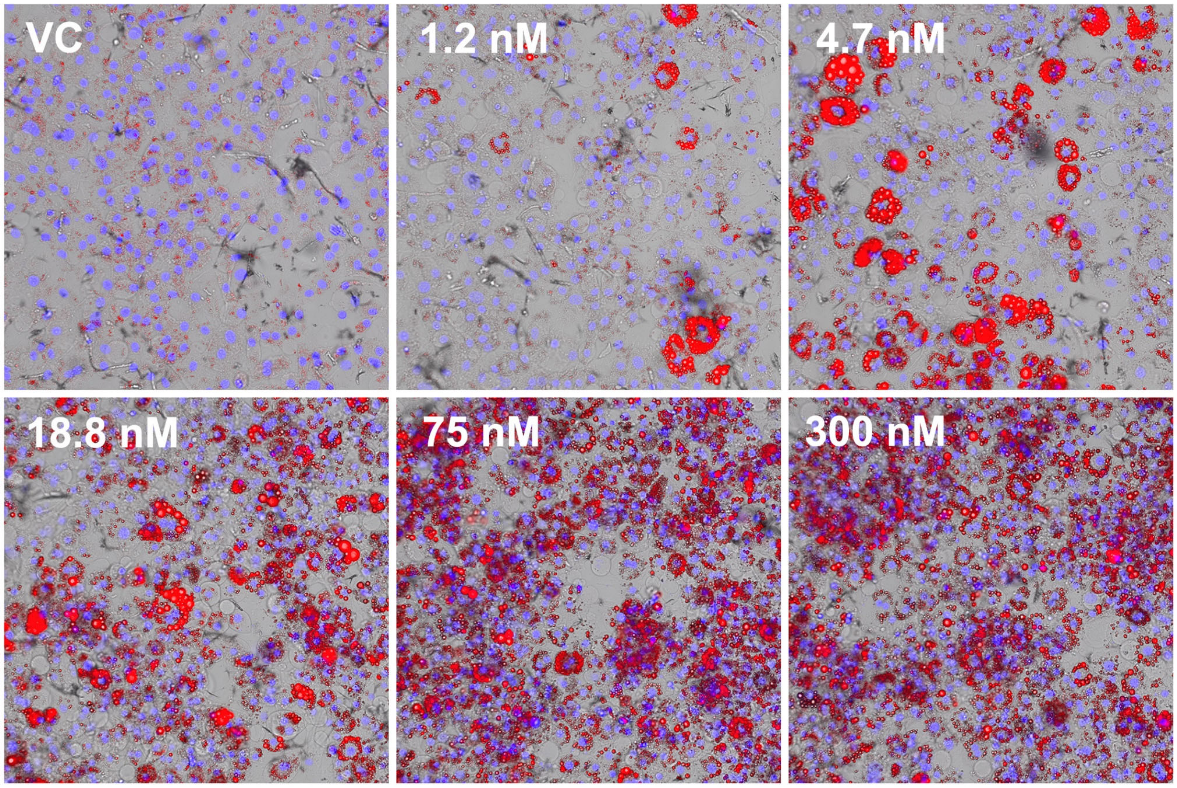

**Fig. S4. Dose-dependent induction of adipogenesis in 3T3-L1 cells exposed to the reference compound rosiglitazone.** Merged brightfield and fluorescence images. Nuclei are stained with NucBlue (blue) and triglycerides with NileRed (red). VC = vehicle control. Raw pictures were processed in the same manner for visualization.

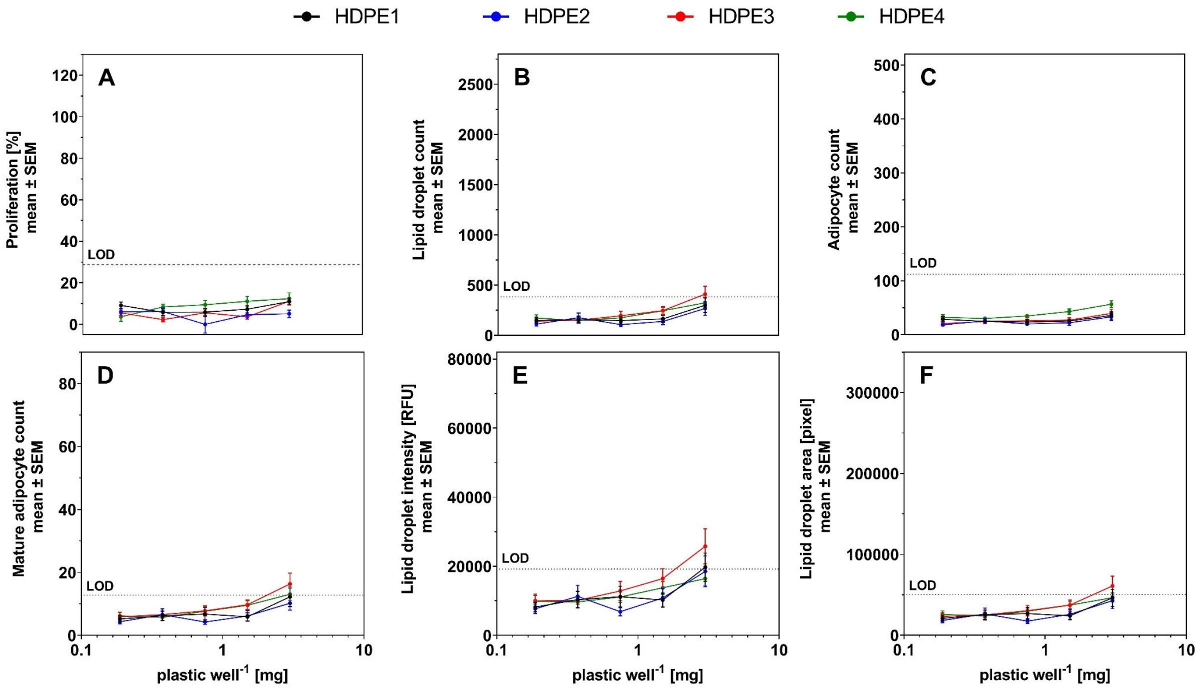

**Fig. S5. Dose-response relationship for the HDPE plastic extracts (HDPE 1–4) in the adipogenesis assay.** (A) proliferation normalized on the mean of the vehicle control, (B) lipid droplet count per field, (C) adipocyte count per field, (D) mature adipocyte count per field, (E) total intensity of the NileRed staining within the lipid droplet mask per field and (F) total area occupied by lipid droplets per field. Twelve or more replicates per concentration ($n \geq 12$). LOD = limit of detection, RFU = relative fluorescence unit.

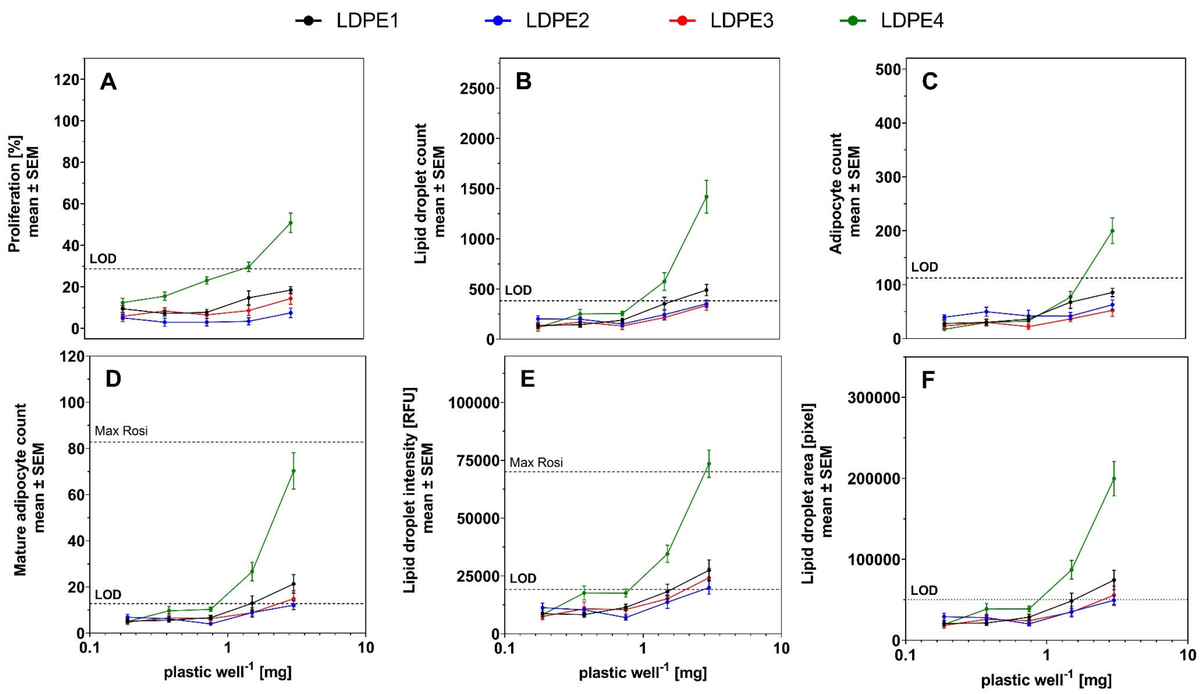

**Fig. S6. Dose-response relationship for the LDPE plastic extracts (LDPE 1–4) in the adipogenesis assay.** (A) proliferation normalized on the mean of the vehicle control, (B) lipid droplet count per field, (C) adipocyte count per field, (D) mature adipocyte count per field, (E) total intensity of the NileRed staining within the lipid droplet mask per field and (F) total area occupied by lipid droplets per field. Twelve or more replicates per concentration (n ≥ 12). LOD = limit of detection, Max Rosi = rosiglitazone maximal response, RFU = relative fluorescence unit.

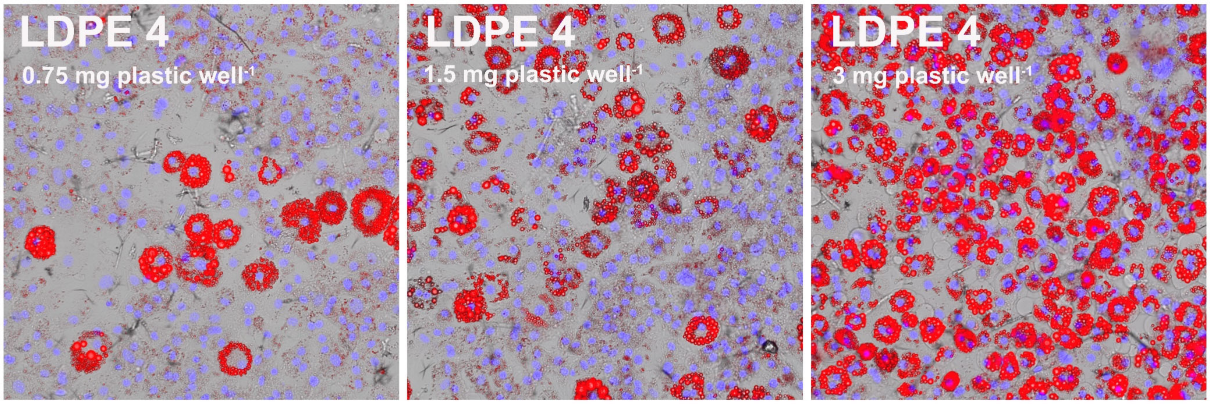

**Fig. S7. Dose-dependent induction of adipogenesis in 3T3-L1 cells exposed to the active plastic extract LDPE 4.** Merged brightfield and fluorescence images. Nuclei are stained with NucBlue (blue) and triglycerides with NileRed (red). Raw pictures were processed in the same manner for visualization.

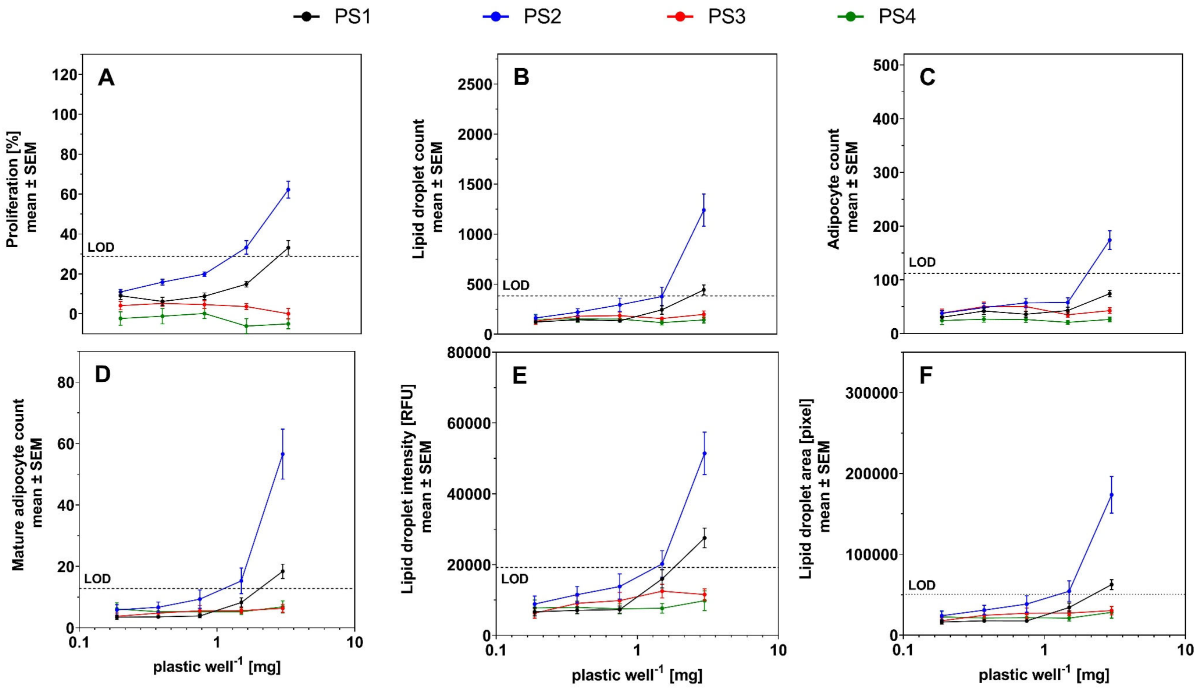

**Fig. S8. Dose-response relationships for the PS plastic extracts (PS 1–4) in the adipogenesis assay.** (A) proliferation normalized on the mean of the vehicle control, (B) lipid droplet count per field, (C) adipocyte count per field, (D) mature adipocyte count per field, (E) total intensity of the NileRed staining within the lipid droplet mask per field and (F) total area occupied by lipid droplets per field. Twelve or more replicates per concentration (n ≥ 12). LOD = limit of detection, RFU = relative fluorescence unit.

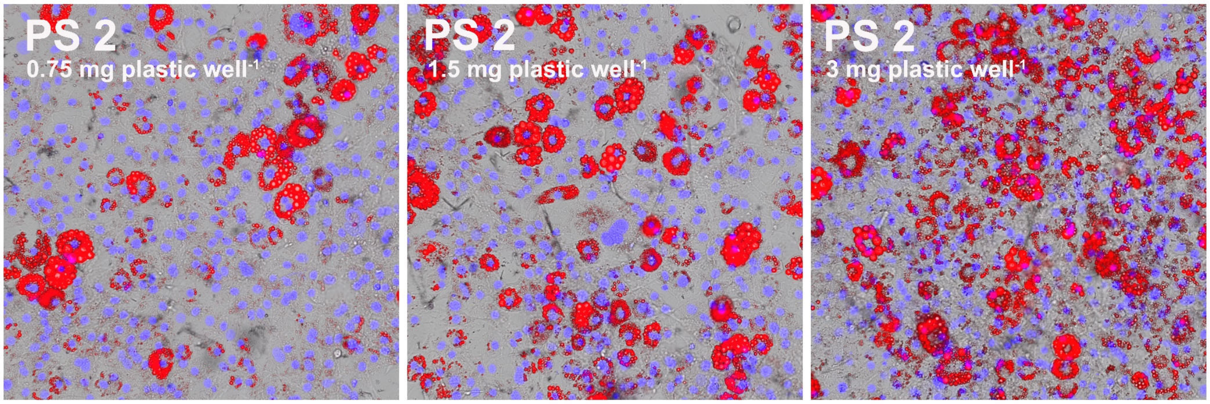

**Fig. S9. Dose-dependent induction of adipogenesis in 3T3-L1 cells exposed to the active plastic extract PS 2.** Merged brightfield and fluorescence images. Nuclei are stained with NucBlue (blue) and triglycerides with NileRed (red). Raw pictures were processed in the same manner for visualization.

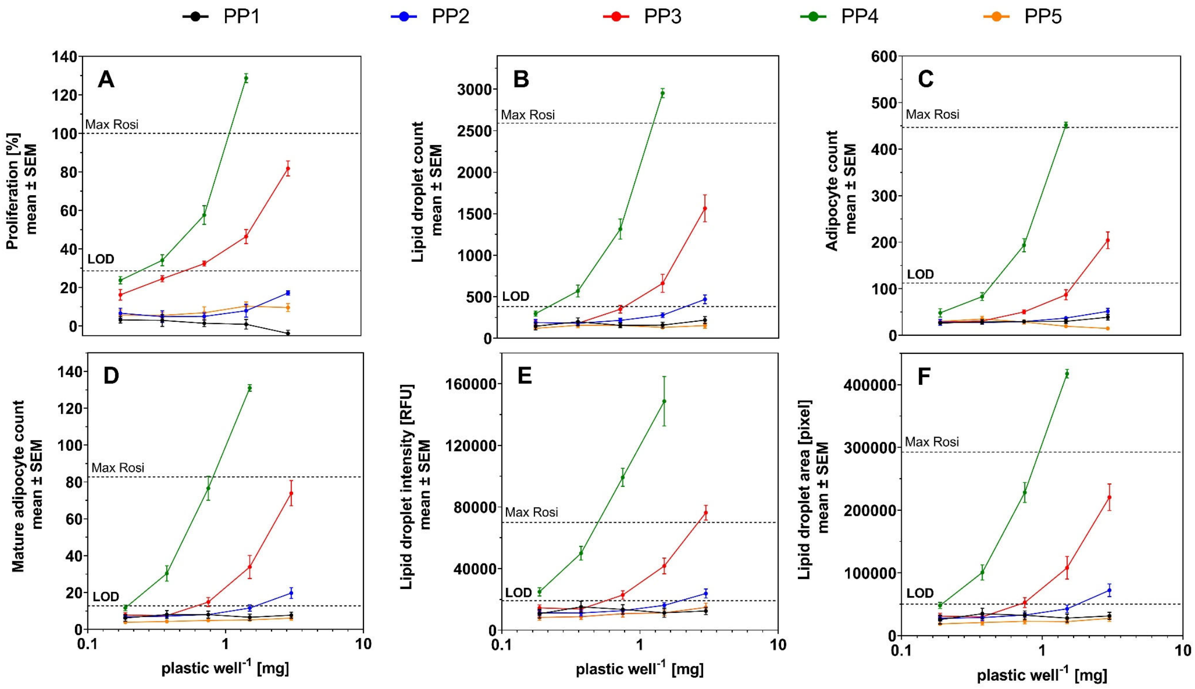

**Fig. S10. Dose-response relationship for the PP plastic extracts (PP 1–5) in the adipogenesis assay.** (A) proliferation normalized on the mean of the vehicle control, (B) lipid droplet count per field, (C) adipocyte count per field, (D) mature adipocyte count per field, (E) total intensity of the NileRed staining within the lipid droplet mask per field and (F) total area occupied by lipid droplets per field. Twelve or more replicates per concentration ($n \geq 12$). LOD = limit of detection, Max Rosi = rosiglitazone maximal response, RFU = relative fluorescence unit.

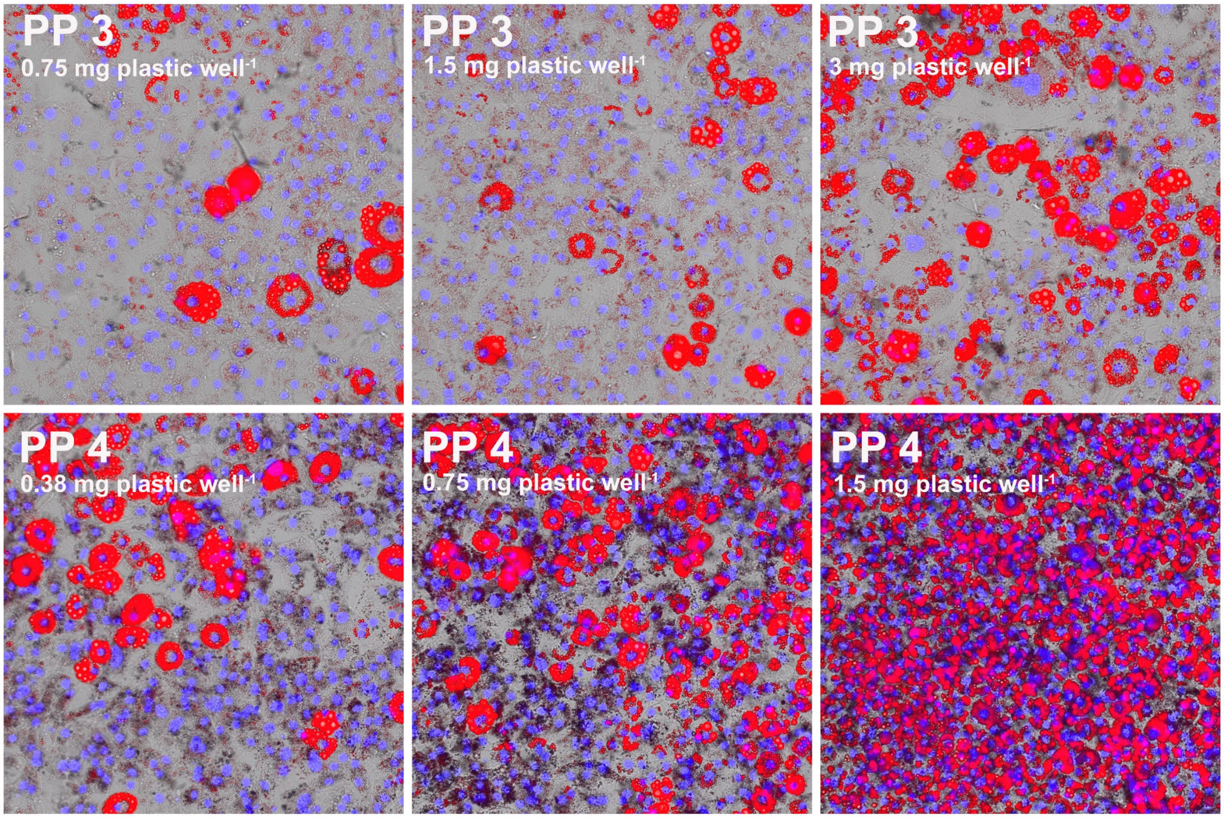

**Fig. S11. Dose-dependent induction of adipogenesis in 3T3-L1 cells exposed to the active plastic extracts PP 3 and PP 4.** Merged brightfield and fluorescence images. Nuclei are stained with NucBlue (blue) and triglycerides with NileRed (red). Raw pictures were processed in the same manner for visualization.

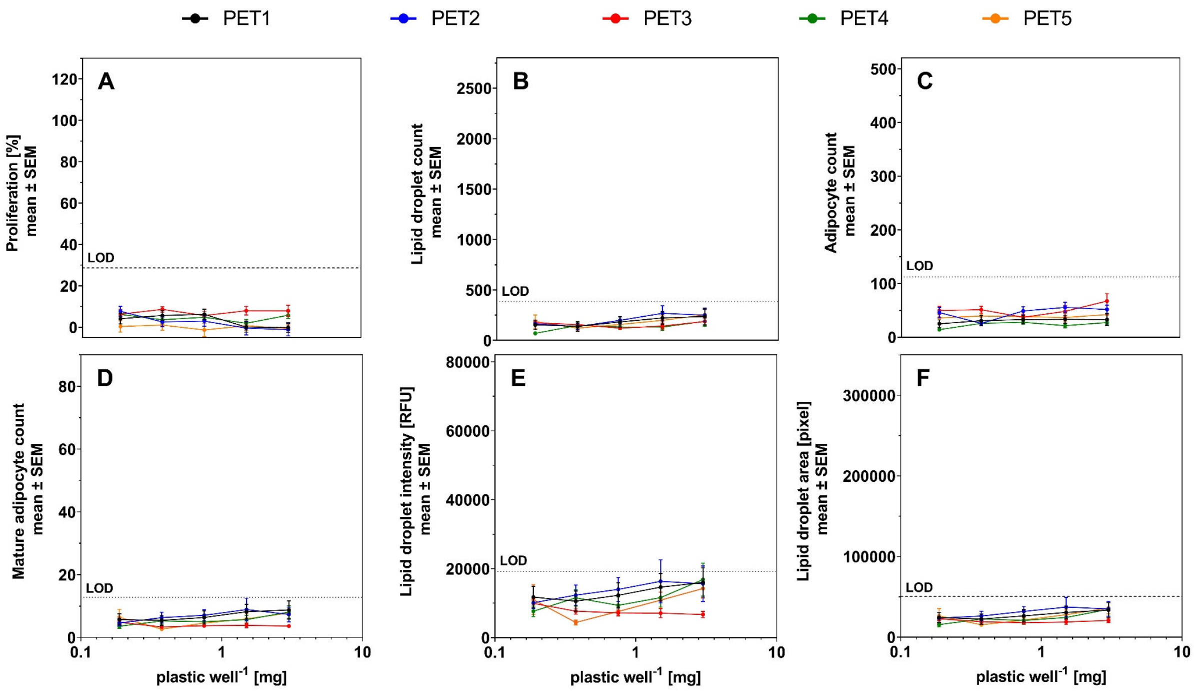

**Fig. S12. Dose-response relationship for the PET plastic extracts (PET 1–5) in the adipogenesis assay.** (A) proliferation normalized on the mean of the vehicle control, (B) lipid droplet count per field, (C) adipocyte count per field, (D) mature adipocyte count per field, (E) total intensity of the NileRed staining within the lipid droplet mask per field and (F) total area occupied by lipid droplets per field. Twelve or more replicates per concentration (n ≥ 12). LOD = limit of detection, RFU = relative fluorescence unit.

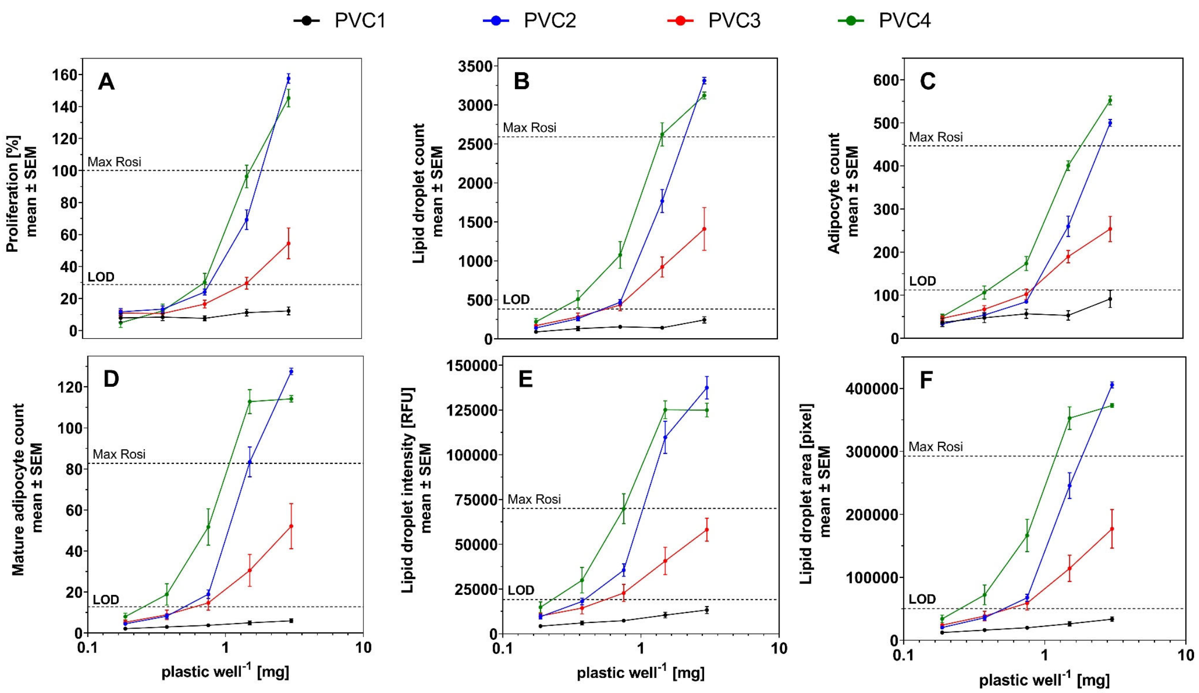

**Fig. S13. Dose-response relationship for the PVC plastic extracts (PVC 1–4) in the adipogenesis assay.** (A) proliferation normalized on the mean of the vehicle control, (B) lipid droplet count per field, (C) adipocyte count per field, (D) mature adipocyte count per field, (E) total intensity of the NileRed staining within the lipid droplet mask per field and (F) total area occupied by lipid droplets per field. Twelve or more replicates per concentration ($n \geq 12$). LOD = limit of detection, Max Rosi = rosiglitazone maximal response, RFU = relative fluorescence unit.

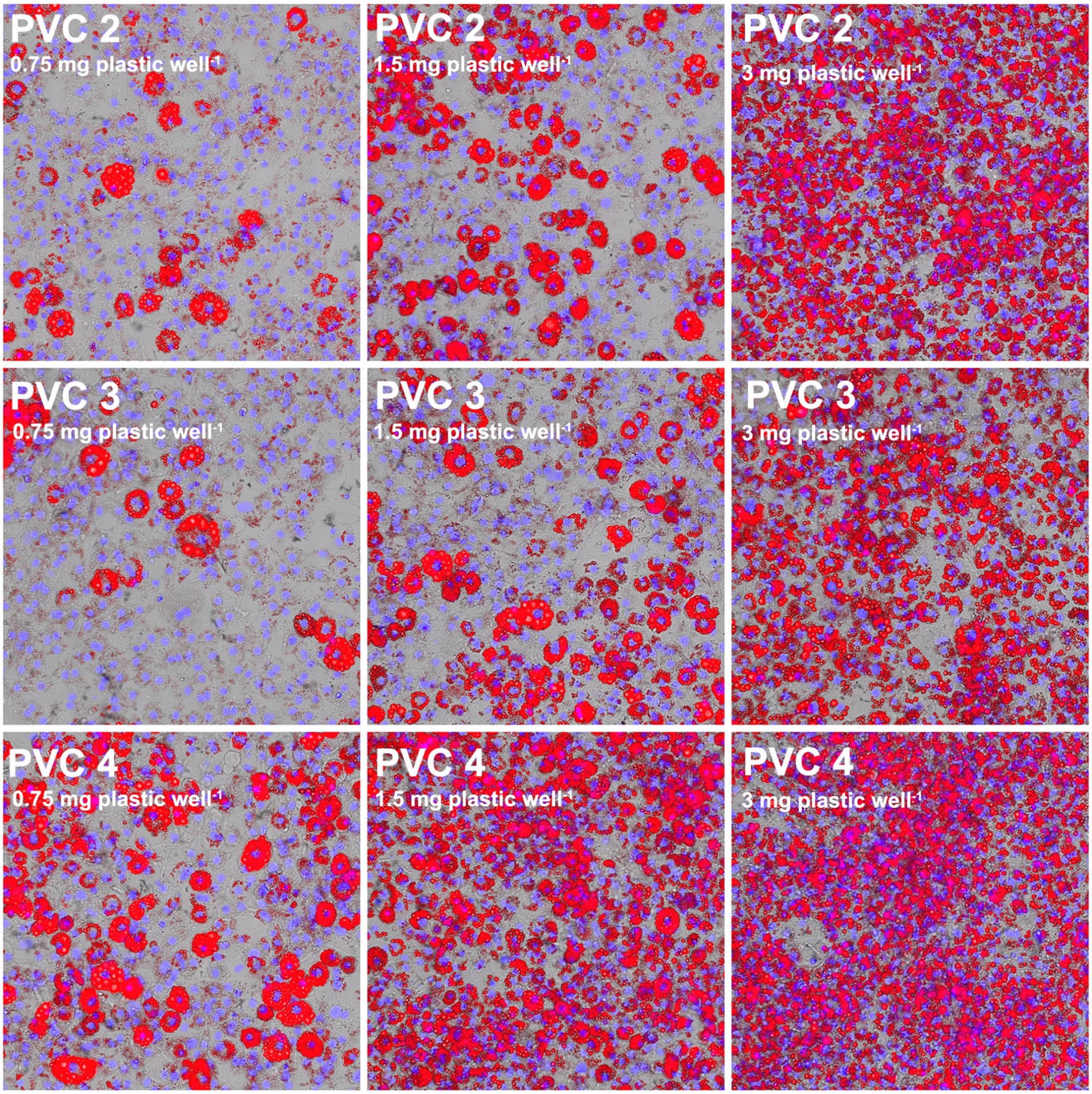

**Fig. S14. Dose-dependent induction of adipogenesis in 3T3-L1 cells exposed to the active plastic extracts PVC 2– 4.** Merged brightfield and fluorescence images. Nuclei are stained with NucBlue (blue) and triglycerides with NileRed (red). Raw pictures were processed in the same manner for visualization.

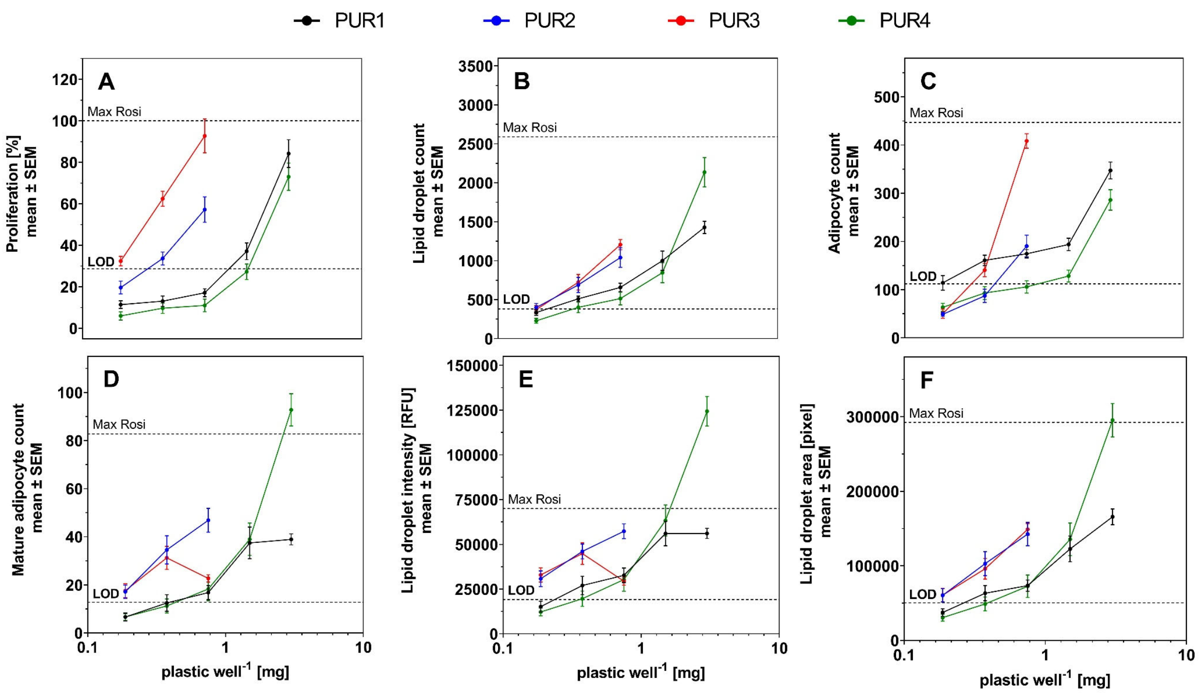

**Fig. S15. Dose-response relationship for the PUR plastic extracts (PUR 1–4) in the adipogenesis assay.** (A) proliferation normalized on the mean of the vehicle control, (B) lipid droplet count per field, (C) adipocyte count per field, (D) mature adipocyte count per field, (E) total intensity of the NileRed staining within the lipid droplet mask per field and (F) total area occupied by lipid droplets per field. Twelve or more replicates per concentration (n ≥ 12). LOD = limit of detection, Max Rosi = rosiglitazone maximal response, RFU = relative fluorescence unit.

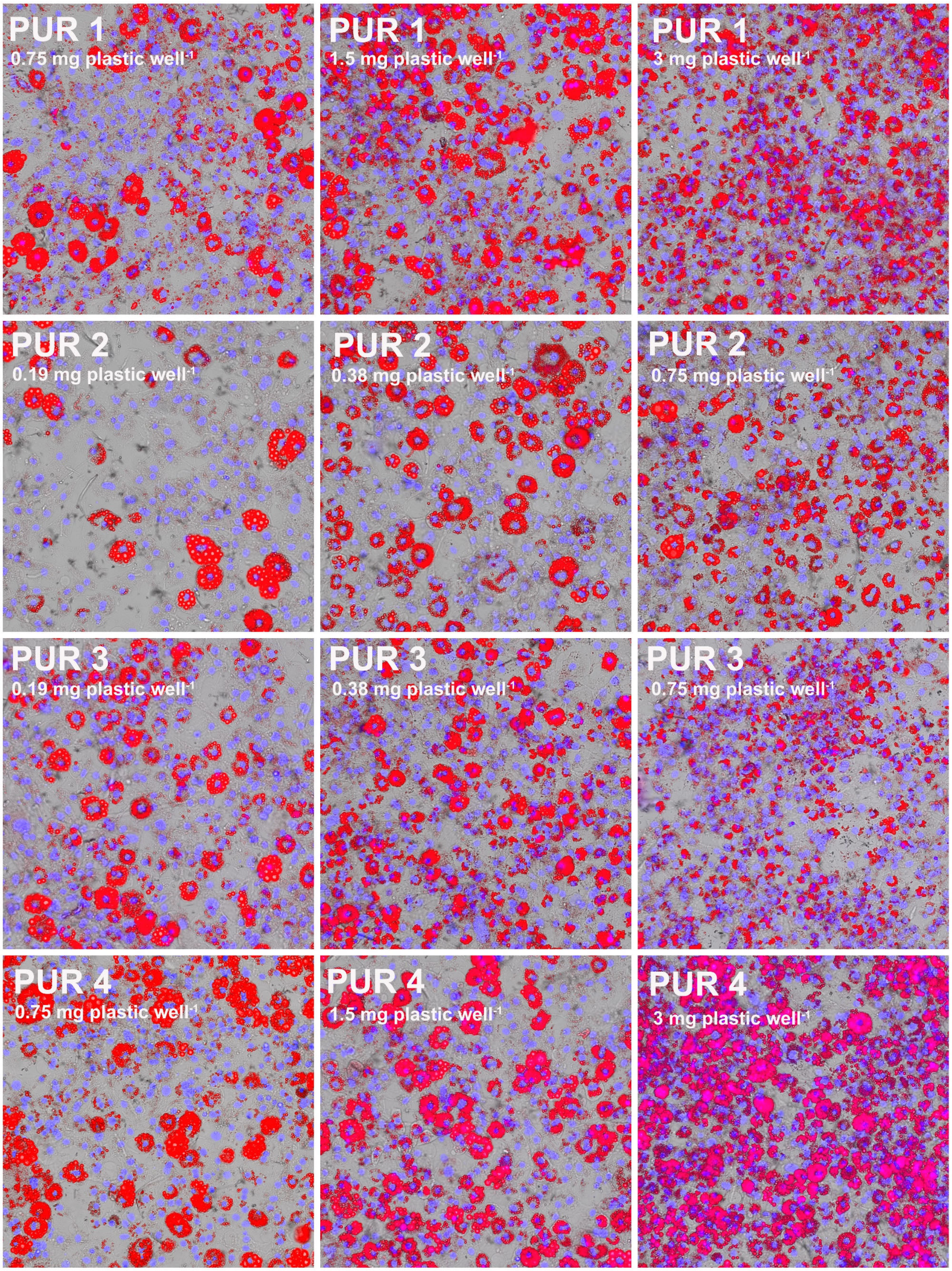

**Fig. S16. Dose-dependent induction of adipogenesis in 3T3-L1 cells exposed to the active plastic extracts PUR 1-4.** Merged brightfield and fluorescence images. Nuclei are stained with NucBlue (blue) and triglycerides with NileRed (red). Raw pictures were processed in the same manner for visualization.

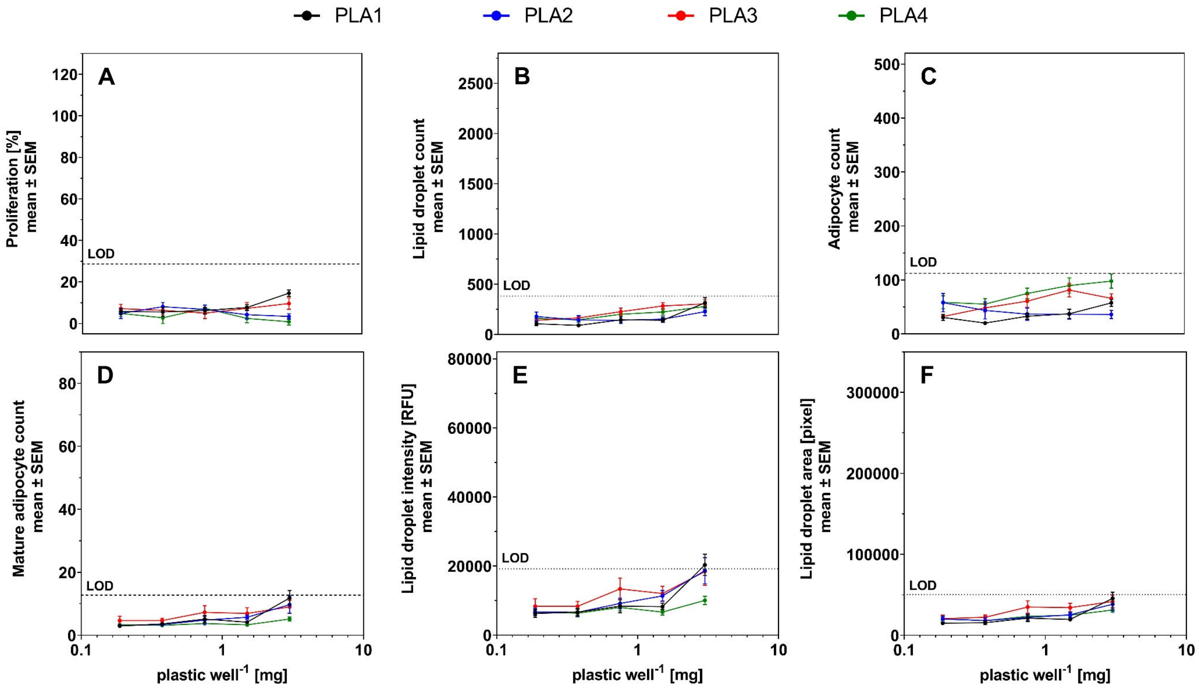

**Fig. S17. Dose-response relationship for the PLA plastic extracts (PLA 1–4) in the adipogenesis assay.** (A) proliferation normalized on the mean of the vehicle control, (B) lipid droplet count per field, (C) adipocyte count per field, (D) mature adipocyte count per field, (E) total intensity of the NileRed staining within the lipid droplet mask per field and (F) total area occupied by lipid droplets per field. Twelve or more replicates per concentration ($n \geq 12$). LOD = limit of detection, Max Rosi = rosiglitazone maximal response, RFU = relative fluorescence unit.

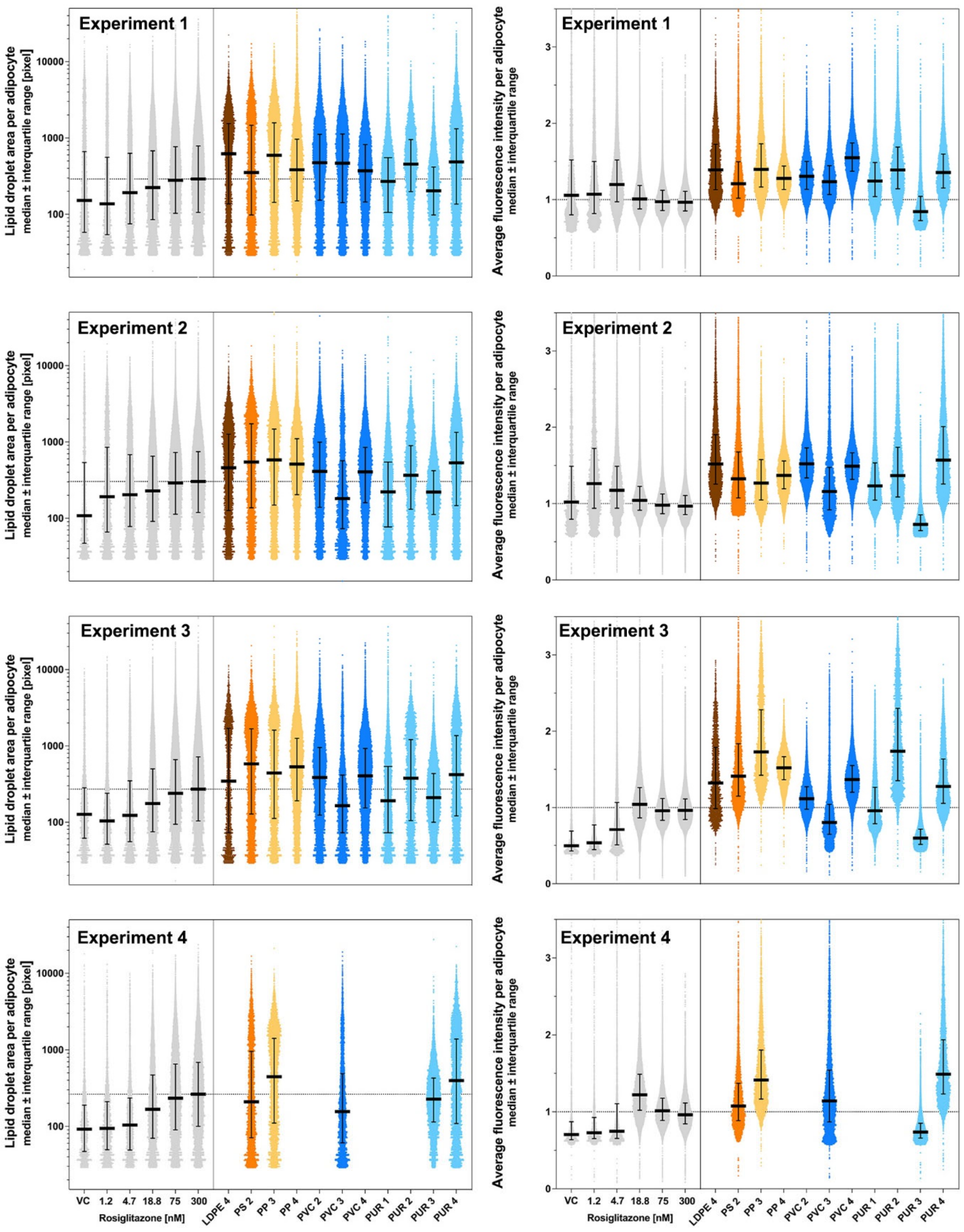

**Fig. S18. Size distribution of adipocyte population (left) and accumulation of triglyceride per adipocyte in cells exposed to rosiglitazone or the highest noncytotoxic concentration of the eleven active plastic extracts.** Single-cell data from 4 independent experiments. Intensity data is normalized to the mean of the highest rosiglitazone concentration (300 nM). VC = vehicle control.

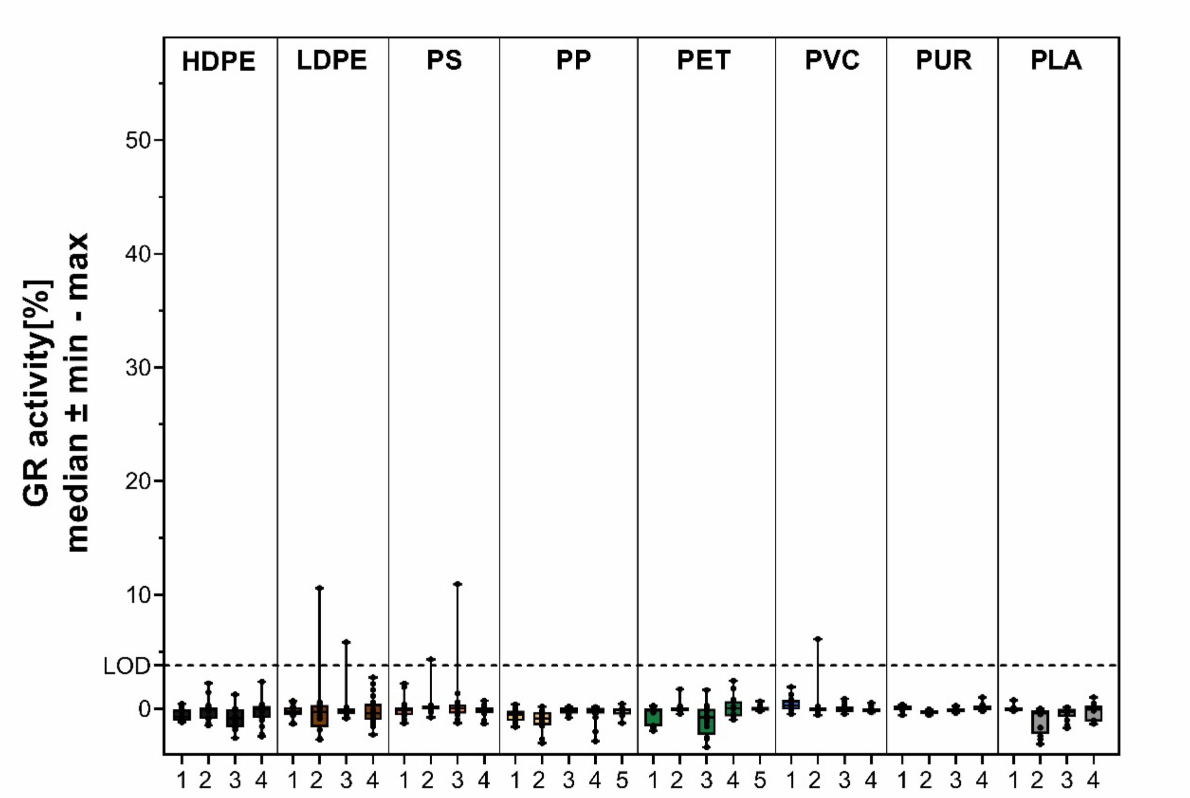

**Fig. S19. GR activity of plastic extracts at the highest noncytotoxic concentration.** Highest noncytotoxic concentration was 1.5 mg plastic $well^{-1}$ except for PP 4 (0.19 mg plastic $well^{-1}$), PS 2 and PP 3 (0.38 mg plastic $well^{-1}$), PLA 1, PVC 2 and PVC 4 (0.75 mg plastic $well^{-1}$). GR = glucocorticoid receptor, LOD = limit of detection. n ≥ 12.

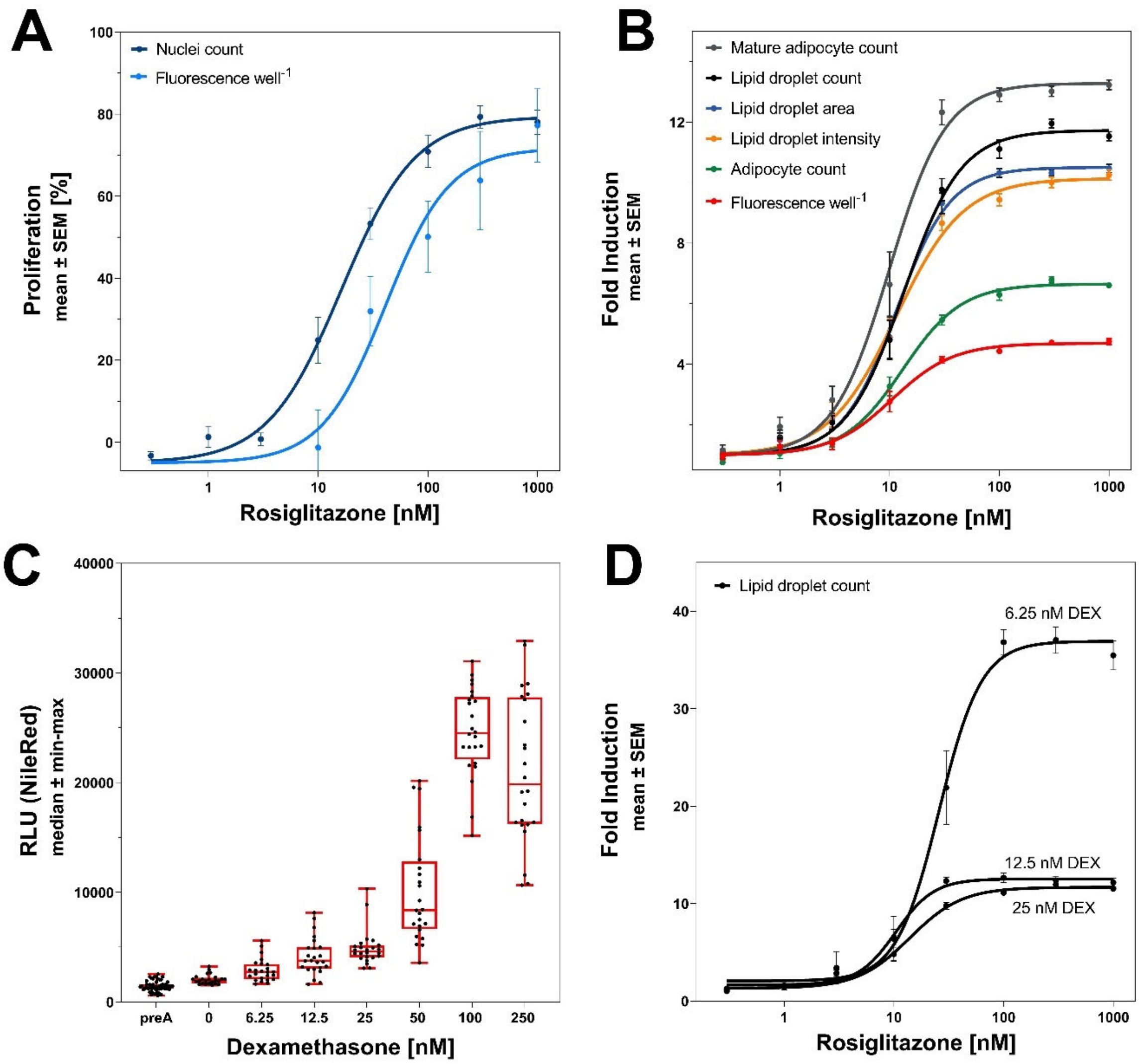

**Fig. S20. Optimization of the adipogenesis assay.** (A) Proliferative effects induced by rosiglitazone analyzed by imaging (DAPI filter) and total fluorescence readout (NucBlue) with 25 nM dexamethasone (DEX) in the differentiation medium (DM). (B) Multiple adipogenic endpoints analyzed by imaging (RFP filter) compared to total fluorescence readout (NileRed) with 25 nM DEX in DM. (C) Total fluorescence readout (NileRed) for the preadipocyte control (preA, undifferentiated) and the differentiated vehicle controls with increasing dexamethasone concentrations in DM. (D) Lipid droplet count induction by rosiglitazone with 6.25, 12.5, 25 nM DEX in DM.

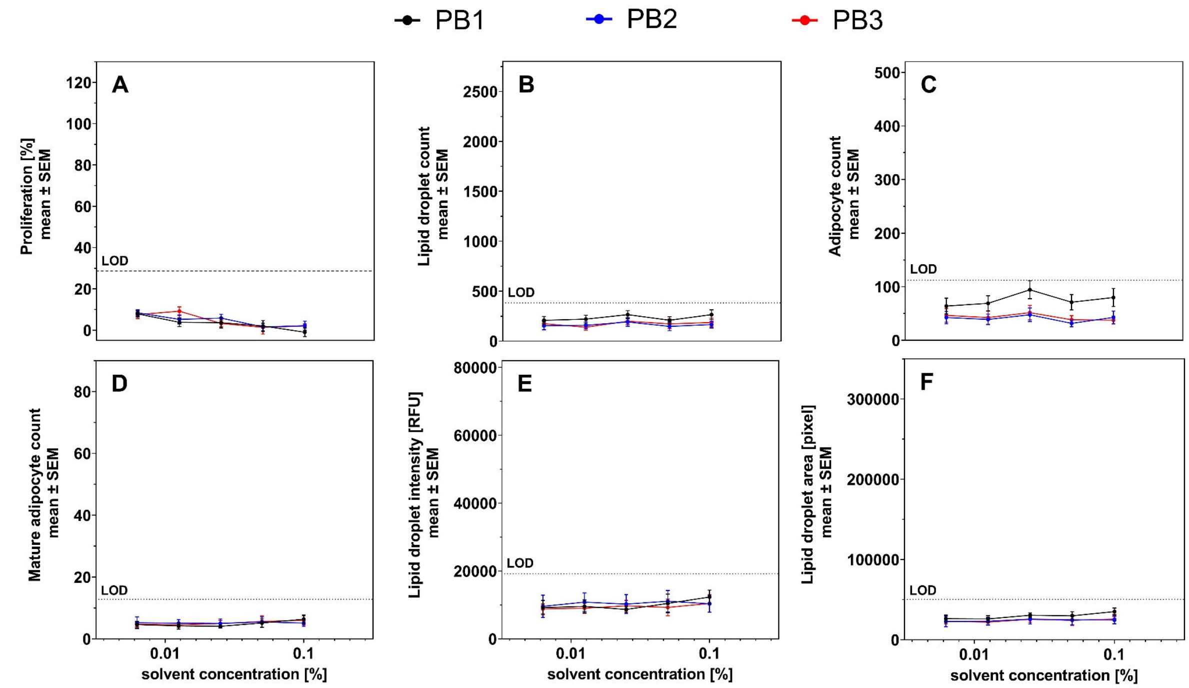

**Fig. S21. Dose-response relationships for the adipogenesis assay endpoints of the three procedure blanks (PB 1–3).** (A) proliferation normalized on the mean of the vehicle control, (B) lipid droplet count per field, (C) adipocyte count per field, (D) mature adipocyte count per field, (E) total intensity of the NileRed staining within the lipid droplet mask per field and (F) total area occupied by lipid droplets per field. Twelve replicates per concentration (n = 12). LOD = limit of detection, RFU = relative fluorescence unit.

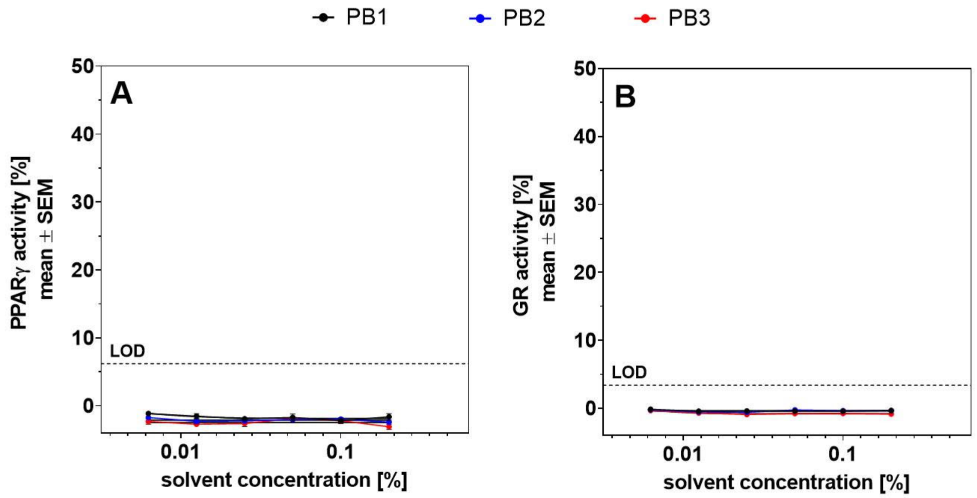

**Fig. S22. Dose-response relationships for (A) PPARγ and (B) GR activity of the three procedure blanks (PB 1–3).** Twelve or more replicates per concentration (n ≥ 12). PPARγ = peroxisome proliferator receptor gamma, GR = glucocorticoid receptor, LOD = limit of detection.

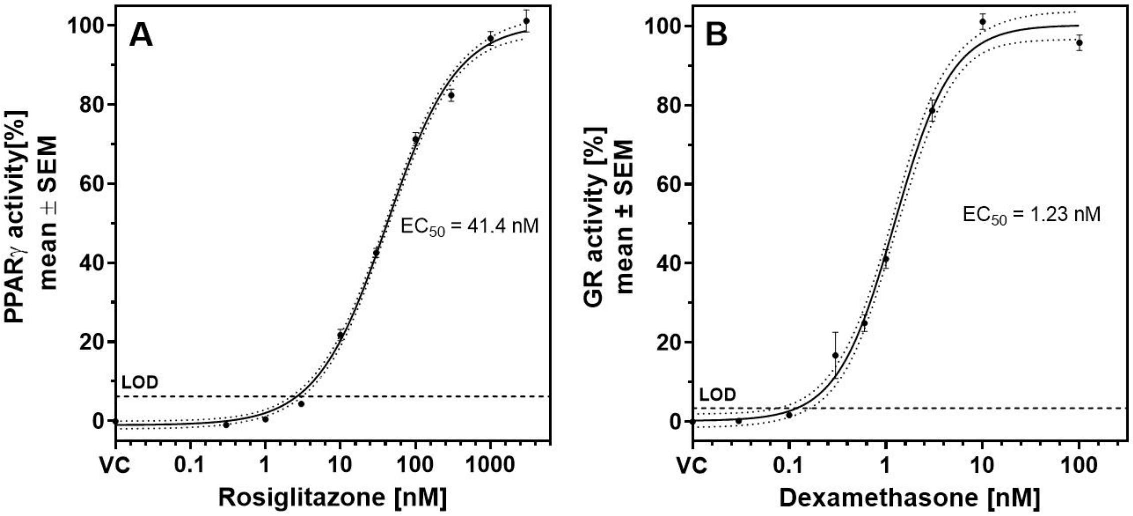

**Fig. S23. Dose-response relationships for (A) PPARγ and (B) GR activity of the reference compound rosiglitazone and dexamethasone.** 48 or more replicates per concentration ($n \geq 48$). PPARγ = peroxisome proliferator receptor gamma, GR = glucocorticoid receptor, LOD = limit of detection, VC = vehicle control.

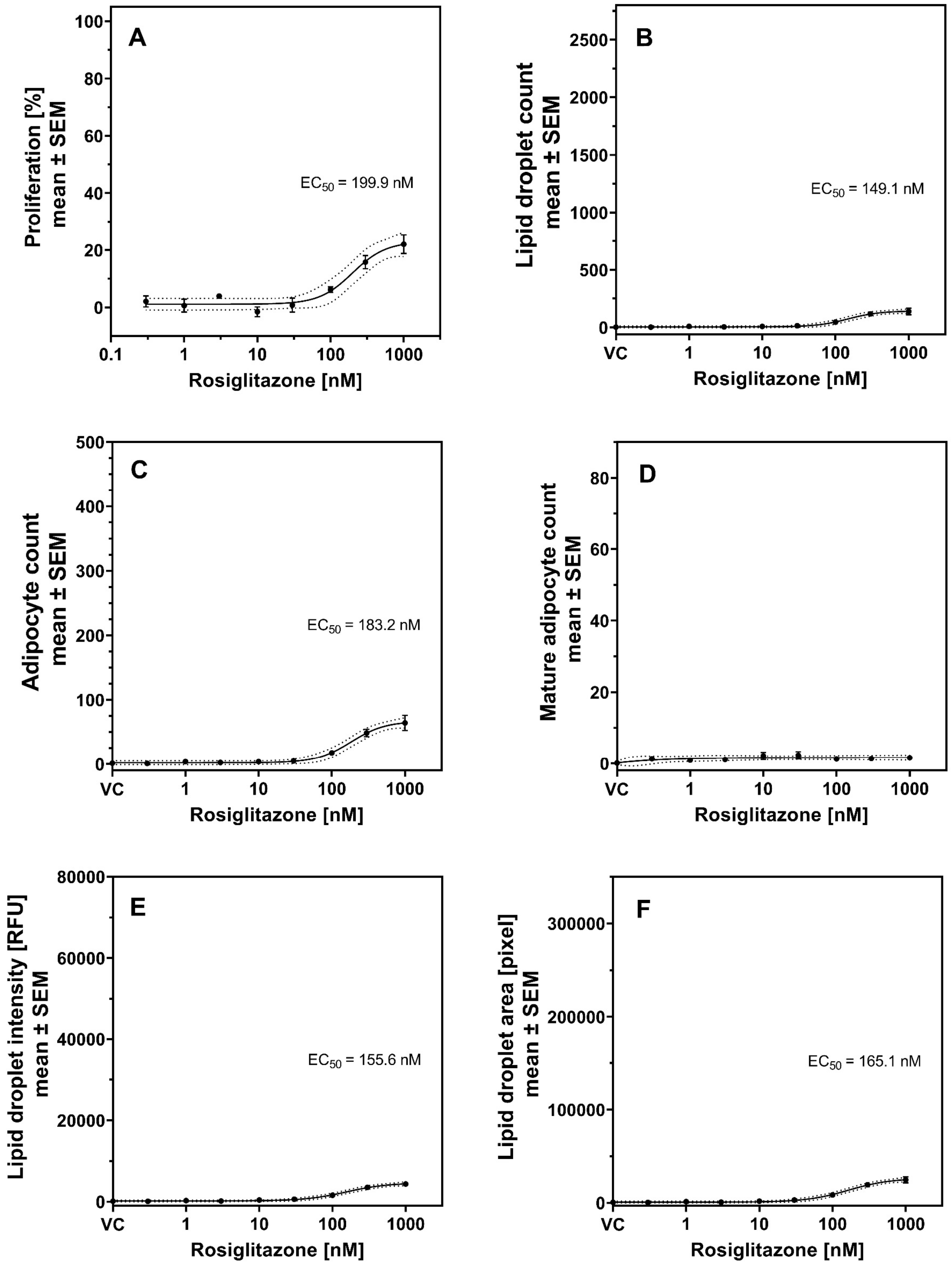

**Fig. S24. Dose-response relationship for the reference compound rosiglitazone in the adipogenesis assay without dexamethasone in the differentiation medium.** (A) proliferation normalized on the mean of the vehicle control, (B) lipid droplet count per field, (C) adipocyte count per field, (D) mature adipocyte count per field, (E) total intensity of the NileRed staining within the lipid droplet mask per field and (F) total area occupied by lipid droplets per field. Eight or more replicates per concentration (n ≥ 8). VC = vehicle control, RFU = relative fluorescence unit.

**Table S1. Cell viability (%) of the cytotoxic samples in the reporter gene assay experiments.** PPARγ = peroxisome proliferator receptor gamma, GR = glucocorticoid receptor, SD = standard deviation.

**PPARγ CALUX**

| mg plastic well$^{-1}$ | PS 2 | | | PP 3 | | | PP 4 | | | PVC 2 | | | PLA1 | | | PVC 4 | | |
|---|---|---|---|---|---|---|---|---|---|---|---|---|---|---|---|---|---|---|
| | mean | SD | n | mean | SD | n | mean | SD | n | mean | SD | n | mean | SD | n | mean | SD | n |
| 1.5 | **9.7** | **5** | 16 | **13.9** | **7.6** | 12 | **22.6** | **13.2** | 16 | **44.9** | **21.6** | 20 | **25.7** | **7.9** | **12** | 107.9 | 14.9 | 20 |
| 0.75 | **68.4** | **14.2** | 16 | **41** | **21** | 12 | **14.9** | **3.8** | 16 | 106.8 | 15.4 | 20 | 103.2 | 15.1 | 12 | 120.9 | 20.3 | 20 |
| 0.38 | 108.2 | 10.6 | 16 | 86.9 | 20.8 | 12 | **19.9** | **9.8** | 16 | 117.5 | 16.3 | 20 | 111.6 | 15.5 | 12 | 126.7 | 20.4 | 20 |
| 0.19 | 120.5 | 7.1 | 16 | 106 | 16.6 | 12 | 97.7 | 25.4 | 16 | 123.9 | 19.9 | 20 | 118.4 | 21 | 12 | 126.6 | 30.5 | 20 |
| 0.09 | 121.7 | 6.3 | 16 | 123 | 20.4 | 12 | 109.8 | 11.8 | 16 | 125.5 | 25.1 | 20 | 128.6 | 24.5 | 12 | 121.8 | 34.6 | 20 |
| 0.04 | 116.8 | 15.7 | 16 | 104 | 26.4 | 12 | 101.8 | 10.9 | 16 | 115.4 | 34 | 20 | 131.9 | 21.5 | 12 | 101.8 | 27.2 | 20 |

**GR CALUX**

| mg plastic well$^{-1}$ | PS 2 | | | PP 3 | | | PP 4 | | | PVC 2 | | | PLA 1 | | | PVC 4 | | |
|---|---|---|---|---|---|---|---|---|---|---|---|---|---|---|---|---|---|---|
| | mean | SD | n | mean | SD | n | mean | SD | n | mean | SD | n | mean | SD | n | mean | SD | n |
| 1.5 | **7** | **7.1** | 12 | **43.2** | **13** | 12 | **8.8** | **8.6** | 16 | **52** | **38.7** | 16 | **45.1** | **42.1** | 12 | **71** | **20** | 12 |
| 0.75 | **76.1** | **45.9** | 12 | **57.3** | **13.4** | 12 | **13** | **9.1** | 16 | 109.8 | 10.6 | 16 | 104.3 | 9.3 | 12 | 96 | 9 | 12 |
| 0.38 | 101 | 38.4 | 12 | 88.9 | 15.3 | 12 | **17.5** | **25.2** | 16 | 118.3 | 7.6 | 16 | 112.7 | 6.9 | 12 | 110 | 10 | 12 |
| 0.19 | 117 | 23.7 | 12 | 110.4 | 5.8 | 12 | 96.8 | 18.4 | 16 | 117.2 | 8.5 | 16 | 112.2 | 6 | 12 | 116 | 6 | 12 |
| 0.09 | 113.3 | 19.6 | 12 | 115.2 | 7.5 | 12 | 118.4 | 17.8 | 16 | 118.9 | 8 | 16 | 113.9 | 6.9 | 12 | 113 | 12 | 12 |
| 0.04 | 111.2 | 31.7 | 12 | 103.8 | 10.4 | 12 | 105.8 | 13.4 | 16 | 104.5 | 6.8 | 16 | 107.5 | 9.6 | 12 | 102 | 11 | 12 |

**Table S2. The chemicals tentatively identified using GC-QTOF-MS/MS (data from Zimmermann *et al.* (*15*)).**

| Sample | CAS | PubChem CID | Name according to NIST | Mass | Formula | Score |
|---|---|---|---|---|---|---|
| HDPE 1 | 1000374-06-1[a] | 3023652 | 1,7-di-iso-propylnaphthalene | 212.16 | C16H20 | 71.58 |
| | 10233-13-3 | 25068 | Dodecanoic acid, 1-methylethyl ester | 242.23 | C15H30O2 | 72.54 |
| | 138345-00-3 | 605776 | 7,9-Di-tertbutyl-1-oxaspiro[4,5]deca-6,9-dien-8-one | 262.19 | C17H26O2 | 70.98 |
| | 24169-43-5 | 595960 | 1,3-Dioxolane, 2-(bromomethyl)-2-(2-methylphenyl)- | 256.01 | C11H13BrO2 | 72.61 |
| | 87-97-8 | 6911 | Phenol, 2,6-bis(1,1-dimethylethyl)-4-(methoxymethyl)- | 250.19 | C16H26O2 | 70.81 |
| HDPE 2 | 4337-65-9 | 20342 | Hexanedioic acid, mono(2-ethylhexyl)ester | 258.18 | C14H26O4 | 82.22 |
| | 6386-38-5 | 62603 | Benzenepropanoic acid, 3,5-bis(1,1-dimethylethyl)-4-hydroxy-, methyl ester | 292.20 | C18H28O3 | 71.02 |
| HDPE 3 | 96-76-4 | 7311 | 2,4-Di-tert-butylphenol | 206.17 | C14H22O | 80.64 |
| HDPE 4 | 1000192-65-0[a] | 543423 | 1,2-15,16-Diepoxyhexadecane | 254.23 | C16H30O2 | 72.96 |
| | 1000315-44-3[a] | 6423518 | Phthalic acid, isobutyl tridec-2-yn-1-yl ester | 400.26 | C25H36O4 | 72.55 |
| | 1000336-60-4[a] | 44515574 | i-Propyl 12-methyl-tridecanoate | 270.26 | C17H34O2 | 73.73 |
| | 1000374-06-1[a] | 3023652 | 1,7-di-iso-propylnaphthalene | 212.16 | C16H20 | 79.77 |
| | 101-86-0 | 7585 | Octanal, 2-(phenylmethylene)- | 216.15 | C15H20O | 85.79 |
| | 105794-58-9 | 537071 | 1-Heptatriacotanol | 536.59 | C37H76O | 70.03 |
| | 13466-78-9 | 26049 | 3-Carene | 136.13 | C10H16 | 92.31 |
| | 16507-61-2 | 5367784 | cis-1-Chloro-9-octadecene | 286.24 | C18H35Cl | 74.15 |
| | 242794-76-9 | 564746 | Bicyclo[5.2.0]nonane, 2-methylene-4,8,8-trimethyl-4-vinyl- | 204.19 | C15H24 | 85.25 |
| | 294-62-2 | 9268 | Cyclododecane | 168.19 | C12H24 | 74.42 |
| | 499-97-8 | 68140 | Cyclohexane, 1-methylene-4-(1-methylethenyl)- | 136.13 | C10H16 | 88.01 |
| | 535-77-3 | 10812 | Benzene, 1-methyl-3-(1-methylethyl)- | 134.11 | C10H14 | 84.43 |
| | 546-80-5 | 261491 | Bicyclo[3.1.0]hex-2-ene, 2-methyl-5-(1-methylethyl)- | 136.13 | C10H16 | 89.31 |
| | 586-63-0 | 102443 | Cyclohexene, 3-methyl-6-(1-methylethylidene)- | 136.13 | C10H16 | 85.52 |
| | 5989-27-5 | 440917 | D-Limonene | 136.13 | C10H16 | 91.24 |
| LDPE 1 | 1000333-58-3[a] | 12541027 | cis-13-Octadecenoic acid, methyl ester | 296.27 | C19H36O2 | 75.47 |
| | 1000336-50-5[a] | 13908974 | Methyl 9-eicosenoate | 324.30 | C21H40O2 | 72.97 |
| | 1000364-37-1[a] | 91738699 | 2-Methyl-3,5-dinitrobenzyl alcohol, TBDMS derivative | 326.13 | C14H22N2O5Si | 71.01 |
| | 1000374-06-1[a] | 3023652 | 1,7-di-iso-propylnaphthalene | 212.16 | C16H20 | 74.02 |
| | 1000374-17-9[a] | 6428435 | 7-epi-cis-sesquisabinene hydrate | 222.20 | C15H26O | 72.79 |
| | 10436-08-5 | 547891 | cis-11-Eicosenamide | 309.30 | C20H39NO | 77.21 |
| | 10436-09-6 | 5365369 | trans-13-Docosenamide | 337.33 | C22H43NO | 84.90 |
| | 10482-56-1 | 443162 | L-.alpha.-Terpineol | 154.14 | C10H18O | 87.36 |
| | 105794-58-9 | 537071 | 1-Heptatriacotanol | 536.59 | C37H76O | 70.34 |
| | 1120-25-8 | 643801 | 9-Hexadecenoic acid, methyl ester, (Z)- | 268.24 | C17H32O2 | 74.59 |
| | 112-63-0 | 5284421 | 9,12-Octadecadienoic acid (Z,Z)-, methyl ester | 294.26 | C19H34O2 | 89.17 |
| | 112-80-1 | 445639 | Oleic Acid | 282.26 | C18H34O2 | 70.26 |
| | 115-99-1 | 61040 | 1,6-Octadien-3-ol, 3,7-dimethyl-, formate | 182.13 | C11H18O2 | 81.34 |
| | 13466-78-9 | 26049 | 3-Carene | 136.13 | C10H16 | 82.42 |
| | 1632-73-1 | 15406 | Fenchol | 154.14 | C10H18O | 80.75 |
| | 1732-10-1 | 15612 | Nonanedioic acid, dimethyl ester | 216.14 | C11H20O4 | 76.33 |
| | 17367-08-7 | 549041 | Ethanol, 2-(9,12-octadecadienyloxy)-, (Z,Z)- | 310.29 | C20H38O2 | 71.76 |

| Sample | CAS | PubChem CID | Name according to NIST | Mass | Formula | Score |
|---|---|---|---|---|---|---|
| | 2091-29-4 | 4668 | 9-Hexadecenoic acid | 254.23 | C16H30O2 | 72.15 |
| | 562-74-3 | 11230 | Terpinen-4-ol | 154.14 | C10H18O | 82.74 |
| | 56554-30-4 | 556196 | 7,10,13-Hexadecatrienoic acid, methyl ester | 264.21 | C17H28O2 | 83.82 |
| | 56630-69-4 | 61268 | 13-Docosenoic acid, methyl ester | 352.33 | C23H44O2 | 90.77 |
| | 57156-97-5 | 5365571 | 12,15-Octadecadienoic acid, methyl ester | 294.26 | C19H34O2 | 89.69 |
| | 5989-27-5 | 440917 | D-Limonene | 136.13 | C10H16 | 82.93 |
| | 60-33-3 | 5280450 | 9,12-Octadecadienoic acid (Z,Z)- | 280.24 | C18H32O2 | 72.77 |
| | 81601-03-8 | 91694967 | Geranyl oleate | 418.38 | C28H50O2 | 75.52 |
| LDPE 2 | 112-62-9 | 5364509 | 9-Octadecenoic acid (Z)-, methyl ester | 296.27 | C19H36O2 | 87.08 |
| | 5129-56-6 | 554144 | Undecanoic acid, 10-methyl-, methyl ester | 214.19 | C13H26O2 | 83.16 |
| | 5129-58-8 | 21204 | Tridecanoic acid, 12-methyl-, methyl ester | 242.23 | C15H30O2 | 73.08 |
| | 5129-61-3 | 110444 | Heptadecanoic acid, 16-methyl-, methyl ester | 298.29 | C19H38O2 | 74.37 |
| | 57156-97-5 | 5365571 | 12,15-Octadecadienoic acid, methyl ester | 294.26 | C19H34O2 | 88.65 |
| | 6386-38-5 | 62603 | Benzenepropanoic acid, 3,5-bis(1,1-dimethylethyl)-4-hydroxy-, methyl ester | 292.20 | C18H28O3 | 73.29 |
| | 96-76-4 | 7311 | 2,4-Di-tert-butylphenol | 206.17 | C14H22O | 82.88 |
| LDPE 3 | 6386-38-5 | 62603 | Benzenepropanoic acid, 3,5-bis(1,1-dimethylethyl)-4-hydroxy-, methyl ester | 292.20 | C18H28O3 | 89.02 |
| | 82304-66-3 | 545303 | 7,9-Di-tert-butyl-1-oxaspiro(4,5)deca-6,9-diene-2,8-dione | 276.17 | C17H24O3 | 82.82 |
| | 96-76-4 | 7311 | 2,4-Di-tert-butylphenol | 206.17 | C14H22O | 81.84 |
| LDPE 4 | 1000131-11-6[a] | 5362676 | Z,Z,Z-1,4,6,9-Nonadecatetraene | 260.25 | C19H32 | 73.26 |
| | 1000131-33-2[a] | 5363633 | Z-(13,14-Epoxy)tetradec-11-en-1-ol acetate | 268.20 | C16H28O3 | 70.02 |
| | 1000142-34-3[a] | 591964 | 2-Adamantanol, 2-(bromomethyl)- | 244.05 | C11H17BrO | 73.95 |
| | 1000192-65-0[a] | 543423 | 1,2-15,16-Diepoxyhexadecane | 254.23 | C16H30O2 | 73.92 |
| | 1000314-35-6[a] | 6421281 | (E,Z,Z)-2,4,7-Tridecatrienal | 192.15 | C13H20O | 70.48 |
| | 1000336-36-7[a] | 14122946 | Methyl 4,7,10,13-hexadecatetraenoate | 262.19 | C17H26O2 | 72.54 |
| | 1000336-38-4[a] | 91694372 | Methyl 3-cis,9-cis,12-cis-octadecatrienoate | 292.24 | C19H32O2 | 71.26 |
| | 1000336-60-4[a] | 44515574 | i-Propyl 12-methyl-tridecanoate | 270.26 | C17H34O2 | 73.43 |
| | 1000352-68-4[a] | 88368751 | Oleyl alcohol, trifluoroacetate | 364.26 | C20H35F3O2 | 75.37 |
| | 1000374-06-1[a] | 3023652 | 1,7-di-iso-propylnaphthalene | 212.16 | C16H20 | 79.77 |
| | 1000374-18-0[a] | 11972555 | .alpha.-acorenol | 222.20 | C15H26O | 86.08 |
| | 1000405-59-5[a] | 91701181 | Fumaric acid, 2-octyl tridec-2-yn-1-yl ester | 406.31 | C25H42O4 | 71.10 |
| | 1000406-96-9[a] | 91697642 | Undec-10-ynoic acid, tridec-2-yn-1-yl ester | 360.30 | C24H40O2 | 74.05 |
| | 1000414-43-3[a] | 91694966 | Geranyl palmitoleate | 390.35 | C26H46O2 | 75.56 |
| | 10287-53-3 | 25127 | Parbenate | 193.11 | C11H15NO2 | 83.57 |
| | 10482-56-1 | 443162 | L-.alpha.-Terpineol | 154.14 | C10H18O | 87.52 |
| | 105794-58-9 | 537071 | 1-Heptatriacotanol | 536.59 | C37H76O | 78.17 |
| | 115-99-1 | 61040 | 1,6-Octadien-3-ol, 3,7-dimethyl-, formate | 182.13 | C11H18O2 | 83.17 |
| | 117066-77-0 | none | 2-((2S,4aR)-4a,8-Dimethyl-1,2,3,4,4a,5,6,7-octahydronaphthalen-2-yl)propan-2-ol | 222.20 | C15H26O | 85.50 |
| | 120-51-4 | 2345 | Benzyl Benzoate | 212.08 | C14H12O2 | 70.07 |
| | 1209-71-8 | 6432005 | 2-Naphthalenemethanol, 1,2,3,4,4a,5,6,7-octahydro-.alpha.,.alpha.,4a,8-tetramethyl-, (2R-cis)- | 222.20 | C15H26O | 87.40 |

| Sample | CAS | PubChem CID | Name according to NIST | Mass | Formula | Score |
|---|---|---|---|---|---|---|
| | 13466-78-9 | 26049 | 3-Carene | 136.13 | C10H16 | 89.92 |
| | 177205-54-8 | 622707 | 1,2,3,4-Tetrahydro-1,4-ethanoanthracene, 9,10-dimethoxy- | 268.15 | C18H20O2 | 81.18 |
| | 17735-94-3 | 5312518 | cis-13-Eicosenoic acid | 310.29 | C20H38O2 | 70.56 |
| | 1845-30-3 | 164888 | cis-Verbenol | 152.12 | C10H16O | 82.59 |
| | 194607-96-0 | 527206 | 2-((4aS,8R,8aR)-4a,8-Dimethyl-3,4,4a,5,6,7,8,8a-octahydronaphthalen-2-yl)propan-2-ol | 222.20 | C15H26O | 85.06 |
| | 21747-46-6 | 10910653 | 1H-Cycloprop[e]azulene, 1a,2,3,5,6,7,7a,7b-octahydro-1,1,4,7-tetramethyl-, [1aR-(1a.alpha.,7.alpha.,7a.beta.,7b.alpha.)]- | 204.19 | C15H24 | 82.27 |
| | 22117-09-5 | 5367371 | 5,8,11-Heptadecatrien-1-ol | 250.23 | C17H30O | 72.18 |
| | 2244-16-8 | 16724 | D-Carvone | 150.10 | C10H14O | 78.01 |
| | 29141-10-4 | 6553885 | (1R,2R,5S)-5-Methyl-2-(prop-1-en-2-yl)cyclohexanol | 154.14 | C10H18O | 80.60 |
| | 294-62-2 | 9268 | Cyclododecane | 168.19 | C12H24 | 73.79 |
| | 29803-82-5 | 122485 | 2-Cyclohexen-1-ol, 1-methyl-4-(1-methylethyl)-, cis- | 154.14 | C10H18O | 70.29 |
| | 353313[a] | 17868 | Bicyclo[3.1.0]hex-2-ene, 2-methyl-5-(1-methylethyl)- | 136.13 | C10H16 | 87.39 |
| | 463-40-1 | 5280934 | 9,12,15-Octadecatrienoic acid, (Z,Z,Z)- | 278.23 | C18H30O2 | 73.20 |
| | 470-40-6 | 11401461 | cis-Thujopsene | 204.19 | C15H24 | 79.33 |
| | 499-97-8 | 68140 | Cyclohexane, 1-methylene-4-(1-methylethenyl)- | 136.13 | C10H16 | 89.08 |
| | 506-26-3 | 5280933 | Gamolenic acid | 278.23 | C18H30O2 | 73.07 |
| | 514-95-4 | 578237 | 1,5,5-Trimethyl-6-methylene-cyclohexene | 136.13 | C10H16 | 76.22 |
| | 535-77-3 | 10812 | Benzene, 1-methyl-3-(1-methylethyl)- | 134.11 | C10H14 | 84.97 |
| | 5392-40-5 | 8843 | Citral | 152.12 | C10H16O | 80.01 |
| | 562-74-3 | 11230 | Terpinen-4-ol | 154.14 | C10H18O | 83.75 |
| | 56666-38-7 | 41961 | 2H-Pyran, tetrahydro-2-(12-pentadecynyloxy)- | 308.27 | C20H36O2 | 72.69 |
| | 584-79-2 | 11442 | Bioallethrin | 302.19 | C19H26O3 | 71.11 |
| | 5989-27-5 | 440917 | D-Limonene | 136.13 | C10H16 | 83.74 |
| | 61465-23-4 | 577045 | (+-)-trans-1-Isopropenyl-4-methyl-1,4-cyclohexanediol | 170.13 | C10H18O2 | 71.12 |
| | 7212-40-0 | 12618691 | 2-Cyclohexen-1-ol, 1-methyl-4-(1-methylethenyl)-, trans- | 152.12 | C10H16O | 72.18 |
| | 7452-79-1 | 24020 | Butanoic acid, 2-methyl-, ethyl ester | 130.10 | C7H14O2 | 76.76 |
| | 77-53-2 | 65575 | Cedrol | 222.20 | C15H26O | 90.06 |
| | 7785-70-8 | 82227 | (1R)-2,6,6-Trimethylbicyclo[3.1.1]hept-2-ene | 136.13 | C10H16 | 82.07 |
| | 7786-67-6 | 24585 | Cyclohexanol, 5-methyl-2-(1-methylethenyl)- | 154.14 | C10H18O | 81.53 |
| | 99-87-6 | 7463 | p-Cymene | 134.11 | C10H14 | 84.23 |
| PS 1 | 100-42-5 | 7501 | Styrene | 104.06 | C8H8 | 92.48 |
| | 131758-71-9 | 562543 | (2,3-Diphenylcyclopropyl)methyl phenyl sulfoxide, trans- | 332.12 | C22H20OS | 76.37 |
| | 20071-09-4 | 11954175 | Benzene, 1,1'-(1,2-cyclobutanediyl)bis-, trans- | 208.13 | C16H16 | 83.78 |
| | 25558-23-0 | 568889 | Cyclobutane, 1,3-diphenyl-, trans- | 208.13 | C16H16 | 71.07 |
| | 538-81-8 | 641683 | 1,3-Butadiene, 1,4-diphenyl-, (E,E)- | 206.11 | C16H14 | 78.93 |
| PS 2 | 1000192-89-2[a] | 5375831 | Thiocarbamic acid, N,N-dimethyl, S-1,3-diphenyl-2-butenyl ester | 311.13 | C19H21NOS | 72.30 |
| | 1000336-60-4[a] | 44515574 | i-Propyl 12-methyl-tridecanoate | 270.26 | C17H34O2 | 81.81 |
| | 100-42-5 | 7501 | Styrene | 104.06 | C8H8 | 86.69 |
| | 10436-08-5 | 547891 | cis-11-Eicosenamide | 309.30 | C20H39NO | 75.46 |

| Sample | CAS | PubChem CID | Name according to NIST | Mass | Formula | Score |
|---|---|---|---|---|---|---|
| | 120-51-4 | 2345 | Benzyl Benzoate | 212.08 | C14H12O2 | 73.31 |
| | 20071-09-4 | 11954175 | Benzene, 1,1'-(1,2-cyclobutanediyl)bis-, trans- | 208.13 | C16H16 | 74.04 |
| | 23470-00-0 | 123409 | Hexadecanoic acid, 2-hydroxy-1-(hydroxymethyl)ethyl ester | 330.28 | C19H38O4 | 81.59 |
| | 29422-13-7 | 34581 | Naphthalene, 1,2,3,4-tetrahydro-2-phenyl- | 208.13 | C16H16 | 72.68 |
| | 301-02-0 | 5283387 | 9-Octadecenamide, (Z)- | 281.27 | C18H35NO | 87.13 |
| | 56728-02-0 | 609923 | Benzene, 1,1'-[2-methyl-2-(phenylthio)cyclopropylidene]bis- | 316.13 | C22H20S | 70.10 |
| PS 3 | 100-42-5 | 7501 | Styrene | 104.06 | C8H8 | 89.70 |
| | 131758-71-9 | 562543 | (2,3-Diphenylcyclopropyl)methyl phenyl sulfoxide, trans- | 332.12 | C22H20OS | 76.42 |
| | 20071-09-4 | 11954175 | Benzene, 1,1'-(1,2-cyclobutanediyl)bis-, trans- | 208.13 | C16H16 | 86.62 |
| | 538-81-8 | 641683 | 1,3-Butadiene, 1,4-diphenyl-, (E,E)- | 206.11 | C16H14 | 80.62 |
| | 56728-02-0 | 609923 | Benzene, 1,1'-[2-methyl-2-(phenylthio)cyclopropylidene]bis- | 316.13 | C22H20S | 77.61 |
| PS 4 | 1000130-80-7[a] | 5363617 | E-11-Methyl-12-tetradecen-1-ol acetate | 268.24 | C17H32O2 | 77.16 |
| | 1000130-81-0[a] | 549821 | 11,13-Dimethyl-12-tetradecen-1-ol acetate | 282.26 | C18H34O2 | 78.57 |
| | 1000192-65-0[a] | 543423 | 1,2-15,16-Diepoxyhexadecane | 254.23 | C16H30O2 | 73.64 |
| | 1000374-06-1[a] | 3023652 | 1,7-di-iso-propylnaphthalene | 212.16 | C16H20 | 72.97 |
| | 1000382-54-3[a] | 91693137 | Carbonic acid, eicosyl vinyl ester | 368.33 | C23H44O3 | 75.68 |
| | 1000406-16-9[a] | 91692473 | Undec-10-ynoic acid, hexadecyl ester | 406.38 | C27H50O2 | 75.59 |
| | 100-41-4 | 7500 | Ethylbenzene | 106.08 | C8H10 | 89.77 |
| | 100-42-5 | 7501 | Styrene | 104.06 | C8H8 | 89.77 |
| | 150-86-7 | 5280435 | Phytol | 296.31 | C20H40O | 77.15 |
| | 17634-51-4 | 561243 | 1,3,5-Cycloheptatriene, 7-ethyl- | 120.09 | C9H12 | 72.65 |
| | 20071-09-4 | 11954175 | Benzene, 1,1'-(1,2-cyclobutanediyl)bis-, trans- | 208.13 | C16H16 | 86.94 |
| | 56728-02-0 | 609923 | Benzene, 1,1'-[2-methyl-2-(phenylthio)cyclopropylidene]bis- | 316.13 | C22H20S | 80.56 |
| | 5989-27-5 | 440917 | D-Limonene | 136.13 | C10H16 | 78.77 |
| | 98-82-8 | 7406 | Benzene, (1-methylethyl)- | 120.09 | C9H12 | 81.26 |
| PP 1 | n.d. | | | | | |
| PP 2 | n.d. | | | | | |
| PP 3 | 1000336-43-6[a] | 20619411 | Methyl 8-methyl-nonanoate | 186.16 | C11H22O2 | 77.31 |
| | 1000339-14-5[a] | 91695412 | Fumaric acid, 2-ethylhexyl undecyl ester | 382.31 | C23H42O4 | 70.28 |
| | 1000368-53-5[a] | 91205583 | Ethyl stearate, 9,12-diepoxy | 340.26 | C20H36O4 | 72.53 |
| | 1000381-53-1[a] | 91726212 | Succinic acid, 2-(2-chlorophenoxy)ethyl ethyl ester | 300.08 | C14H17ClO5 | 71.07 |
| | 109-43-3 | 7986 | Decanedioic acid, dibutyl ester | 314.25 | C18H34O4 | 82.40 |
| | 111-11-5 | 8091 | Octanoic acid, methyl ester | 158.13 | C9H18O2 | 82.15 |
| | 128-37-0 | 31404 | Butylated Hydroxytoluene | 220.18 | C15H24O | 87.82 |
| | 24560-98-3 | 119250 | Oxiraneoctanoic acid, 3-octyl-, cis- | 298.25 | C18H34O3 | 71.24 |
| | 33368-86-4 | 10186592 | 2-(Octanoyloxy)propane-1,3-diyl bis(decanoate) | 526.42 | C31H58O6 | 73.10 |
| | 33368-87-5 | 10436013 | 2-(Decanoyloxy)propane-1,3-diyl dioctanoate | 498.39 | C29H54O6 | 73.20 |
| | 4098-71-9 | 169132 | Isophorone diisocyanate | 222.14 | C12H18N2O2 | 80.96 |
| | 5129-61-3 | 110444 | Heptadecanoic acid, 16-methyl-, methyl ester | 298.29 | C19H38O2 | 80.72 |
| | 628-97-7 | 12366 | Hexadecanoic acid, ethyl ester | 284.27 | C18H36O2 | 75.72 |
| | 6386-38-5 | 62603 | Benzenepropanoic acid, 3,5-bis(1,1-dimethylethyl)-4-hydroxy-, methyl ester | 292.20 | C18H28O3 | 74.23 |

| Sample | CAS | PubChem CID | Name according to NIST | Mass | Formula | Score |
|---|---|---|---|---|---|---|
| | 77-90-7 | 6505 | Tributyl acetylcitrate | 402.23 | C20H34O8 | 76.77 |
| | 87-97-8 | 6911 | Phenol, 2,6-bis(1,1-dimethylethyl)-4-(methoxymethyl)- | 250.19 | C16H26O2 | 77.23 |
| PP 4 | 1000336-62-4[a] | 53745103 | i-Propyl 14-methyl-pentadecanoate | 298.29 | C19H38O2 | 79.06 |
| | 101-68-8 | 7570 | Benzene, 1,1'-methylenebis[4-isocyanato- | 250.07 | C15H10N2O2 | 90.58 |
| | 10436-09-6 | 5365369 | trans-13-Docosenamide | 337.33 | C22H43NO | 72.31 |
| | 301-02-0 | 5283387 | 9-Octadecenamide, (Z)- | 281.27 | C18H35NO | 85.30 |
| | 77-90-7 | 6505 | Tributyl acetylcitrate | 402.23 | C20H34O8 | 84.50 |
| PP 5 | 1000333-58-3[a] | 12541027 | cis-13-Octadecenoic acid, methyl ester | 296.27 | C19H36O2 | 74.59 |
| | 1000336-60-4[a] | 44515574 | i-Propyl 12-methyl-tridecanoate | 270.26 | C17H34O2 | 86.61 |
| | 1000351-75-2[a] | 14574254 | Eicosyl trifluoroacetate | 394.31 | C22H41F3O2 | 80.57 |
| | 1000368-56-5[a] | 91691599 | 2-Hexyldodecyl isobutyrate | 340.33 | C22H44O2 | 80.81 |
| | 1000374-06-1[a] | 3023652 | 1,7-di-iso-propylnaphthalene | 212.16 | C16H20 | 71.86 |
| | 1000382-54-3[a] | 91693137 | Carbonic acid, eicosyl vinyl ester | 368.33 | C23H44O3 | 70.61 |
| | 103-95-7 | 517827 | 3-(4-Isopropylphenyl)-2-methylpropionaldehyde | 190.14 | C13H18O | 87.10 |
| | 112-39-0 | 8181 | Hexadecanoic acid, methyl ester | 270.26 | C17H34O2 | 85.30 |
| | 119-61-9 | 3102 | Benzophenone | 182.07 | C13H10O | 82.23 |
| | 1222-05-5 | 91497 | Cyclopenta[g]-2-benzopyran, 1,3,4,6,7,8-hexahydro-4,6,6,7,8,8-hexamethyl- | 258.20 | C18H26O | 94.81 |
| | 128-37-0 | 31404 | Butylated Hydroxytoluene | 220.18 | C15H24O | 89.19 |
| | 13491-79-7 | 26068 | Cyclohexanol, 2-(1,1-dimethylethyl)- | 156.15 | C10H20O | 78.36 |
| | 2425-77-6 | 95337 | 1-Decanol, 2-hexyl- | 242.26 | C16H34O | 79.62 |
| | 5129-56-6 | 554144 | Undecanoic acid, 10-methyl-, methyl ester | 214.19 | C13H26O2 | 76.80 |
| | 5129-58-8 | 21204 | Tridecanoic acid, 12-methyl-, methyl ester | 242.23 | C15H30O2 | 79.44 |
| | 5129-61-3 | 110444 | Heptadecanoic acid, 16-methyl-, methyl ester | 298.29 | C19H38O2 | 89.70 |
| | 5348-82-3 | 79299 | Acetic acid, chloro-, octadecyl ester | 346.26 | C20H39ClO2 | 79.89 |
| | 55741-10-1 | 615306 | Naphthalene, 6,7-diethyl-1,2,3,4-tetrahydro-1,1,4,4-tetramethyl- | 244.22 | C18H28 | 78.47 |
| | 6386-38-5 | 62603 | Benzenepropanoic acid, 3,5-bis(1,1-dimethylethyl)-4-hydroxy-, methyl ester | 292.20 | C18H28O3 | 76.72 |
| | 7460-74-4 | 81964 | Pentanoic acid, 2-phenylethyl ester | 206.13 | C13H18O2 | 75.64 |
| | 80-54-6 | 228987 | Lilial | 204.15 | C14H20O | 81.43 |
| | 88-29-9 | 6930 | 7-Acetyl-6-ethyl-1,1,4,4-tetramethyltetralin | 258.20 | C18H26O | 78.53 |
| PET 1 | 5989-27-5 | 440917 | D-Limonene | 136.13 | C10H16 | 86.85 |
| PET 2 | n.d. | | | | | |
| PET 3 | n.d. | | | | | |
| PET 4 | n.d. | | | | | |
| PET 5 | n.d. | | | | | |
| PVC 1 | 1000043-05-3[a] | 6452096 | Ethyl iso-allocholate | 436.32 | C26H44O5 | 70.14 |
| | 1000368-53-5[a] | 91205583 | Ethyl stearate, 9,12-diepoxy | 340.26 | C20H36O4 | 82.01 |
| | 1000383-37-7[a] | 5354568 | Glycidyl oleate | 338.28 | C21H38O3 | 75.46 |
| | 111-03-5 | 5283468 | 9-Octadecenoic acid (Z)-, 2,3-dihydroxypropyl ester | 356.29 | C21H40O4 | 70.61 |
| | 119-61-9 | 3102 | Benzophenone | 182.07 | C13H10O | 76.29 |
| | 14290-23-4 | 539937 | Myristin, 1,3-diaceto-2- | 386.27 | C21H38O6 | 79.34 |
| | 17598-94-6 | 33979 | Dodecanoic acid, 1-(hydroxymethyl)-1,2-ethanediyl ester | 456.38 | C27H52O5 | 72.34 |

| Sample | CAS | PubChem CID | Name according to NIST | Mass | Formula | Score |
|---|---|---|---|---|---|---|
| | 26719-54-0 | 537376 | Dodecanoic acid 3-dodecanoyloxy-propyl ester | 440.39 | C27H52O4 | 71.92 |
| | 4337-65-9 | 20342 | Hexanedioic acid, mono(2-ethylhexyl)ester | 258.18 | C14H26O4 | 76.61 |
| | 5129-56-6 | 554144 | Undecanoic acid, 10-methyl-, methyl ester | 214.19 | C13H26O2 | 81.77 |
| | 5129-61-3 | 110444 | Heptadecanoic acid, 16-methyl-, methyl ester | 298.29 | C19H38O2 | 81.70 |
| | 52380-33-3 | 5364432 | 11-Octadecenoic acid, methyl ester | 296.27 | C19H36O2 | 82.24 |
| | 55268-70-7 | 191981 | Hexadecanoic acid, 2,3-bis(acetyloxy)propyl ester | 414.30 | C23H42O6 | 72.01 |
| | 55429-68-0 | 539928 | Eicosanoic acid, 2-(acetyloxy)-1-[(acetyloxy)methyl]ethyl ester | 470.36 | C27H50O6 | 73.40 |
| | 645-66-9 | 69527 | Lauric anhydride | 382.35 | C24H46O3 | 73.38 |
| | 761-35-3 | 99931 | Hexadecanoic acid, 1-(hydroxymethyl)-1,2-ethanediyl ester | 568.51 | C35H68O5 | 71.87 |
| PVC 2 | 1000324-52-2[a] | 91718008 | Adipic acid, isohexyl methyl ester | 244.17 | C13H24O4 | 74.23 |
| | 1000339-40-5[a] | 12151622 | 1,2-Cyclohexanedicarboxylic acid, dinonyl ester | 424.36 | C26H48O4 | 80.59 |
| | 1000339-74-3[a] | 91721826 | 1,2-Cyclohexanedicarboxylic acid, cyclohexylmethyl nonyl ester | 394.31 | C24H42O4 | 78.08 |
| | 1000339-85-1[a] | 91721974 | 1,2-Cyclohexanedicarboxylic acid, 3,5-dimethylcyclohexyl nonyl ester | 408.32 | C25H44O4 | 75.36 |
| | 109-39-7 | 66959 | 2-Butoxyethyl oleate | 382.35 | C24H46O3 | 78.43 |
| | 112-62-9 | 5364509 | 9-Octadecenoic acid (Z)-, methyl ester | 296.27 | C19H36O2 | 80.99 |
| | 117-81-7 | 8343 | Bis(2-ethylhexyl) phthalate | 390.28 | C24H38O4 | 72.28 |
| | 128-37-0 | 31404 | Butylated Hydroxytoluene | 220.18 | C15H24O | 86.70 |
| | 29761-21-5 | 34697 | Phosphoric acid, isodecyl diphenyl ester | 390.20 | C22H31O4P | 74.21 |
| | 301-02-0 | 5283387 | 9-Octadecenamide, (Z)- | 281.27 | C18H35NO | 81.87 |
| | 4337-65-9 | 20342 | Hexanedioic acid, mono(2-ethylhexyl)ester | 258.18 | C14H26O4 | 76.59 |
| PVC 3 | 1000115-60-4[a] | 5369409 | 5-Hexadecenoic acid, 2-methoxy-, methyl ester | 298.25 | C18H34O3 | 71.33 |
| | 1000127-49-8[a] | 590850 | Phen-1,4-diol, 2,3-dimethyl-5-trifluoromethyl- | 206.06 | C9H9F3O2 | 72.01 |
| | 1000131-33-2[a] | 5363633 | Z-(13,14-Epoxy)tetradec-11-en-1-ol acetate | 268.20 | C16H28O3 | 71.95 |
| | 1000215-67-6[a] | none | trans-2,4-Dimethylthiane, S,S-dioxide | 162.07 | C7H14O2S | 73.76 |
| | 1000215-75-3[a] | none | trans-2-methyl-4-n-pentylthiane, S,S-dioxide | 218.13 | C11H22O2S | 82.26 |
| | 1000253-26-1[a] | 569846 | Octanediamide, N,N'-di-benzoyloxy- | 412.16 | C22H24N2O6 | 71.67 |
| | 1000270-36-9[a] | 569440 | Benzamide, N-(1,3-dihydro-2-oxo-4-isobenzofuryl)- | 253.07 | C15H11NO3 | 79.15 |
| | 1000324-49-0[a] | 91713297 | Adipic acid, 2-ethylhexyl tetradecyl ester | 454.40 | C28H54O4 | 70.35 |
| | 1000333-54-0[a] | 15717634 | 17-Octadecynoic acid, methyl ester | 294.26 | C19H34O2 | 70.63 |
| | 1000333-58-3[a] | 12541027 | cis-13-Octadecenoic acid, methyl ester | 296.27 | C19H36O2 | 86.84 |
| | 1000340-22-6[a] | 9814973 | Benzoic acid, tridecyl ester | 304.24 | C20H32O2 | 76.91 |
| | 1000340-22-7[a] | 64671 | Benzoic acid, tetradecyl ester | 318.26 | C21H34O2 | 81.9 |
| | 1000352-68-4[a] | 88368751 | Oleyl alcohol, trifluoroacetate | 364.26 | C20H35F3O2 | 79.42 |
| | 1000356-41-5[a] | 91724604 | Isophthalic acid, butyl 10-chlorodecyl ester | 396.21 | C22H33ClO4 | 71.83 |
| | 1000367-89-7[a] | 90471467 | Benzoic acid, dec-2-yl ester | 262.19 | C17H26O2 | 71.93 |
| | 1000368-75-3[a] | 91711800 | Benzoic acid, 10-chlorodecyl ester | 296.15 | C17H25ClO2 | 72.25 |
| | 1000371-47-7[a] | 21262075 | 1-Nonylcycloheptane | 224.25 | C16H32 | 74.28 |
| | 1000377-71-8[a] | 91719631 | Phthalic acid, nonyl oct-3-yl ester | 404.29 | C25H40O4 | 82.23 |
| | 1000406-16-5[a] | 91692432 | Undec-10-ynoic acid, dodecyl ester | 350.32 | C23H42O2 | 71.23 |
| | 10417-94-4 | 446284 | cis-5,8,11,14,17-Eicosapentaenoic acid | 302.23 | C20H30O2 | 73.46 |
| | 105794-58-9 | 537071 | 1-Heptatriacotanol | 536.59 | C37H76O | 78.32 |

| Sample | CAS | PubChem CID | Name according to NIST | Mass | Formula | Score |
|---|---|---|---|---|---|---|
| | 108511-83-7 | 569871 | 2-Benzoyloxy-1,1,10-trimethyl-6,9-epidioxydecalin | 330.18 | C20H26O4 | 71.18 |
| | 1129-41-5 | 14322 | Carbamic acid, methyl-, 3-methylphenyl ester | 165.08 | C9H11NO2 | 80.66 |
| | 115-89-9 | 8291 | Diphenyl methyl phosphate | 264.06 | C13H13O4P | 78.24 |
| | 119-61-9 | 3102 | Benzophenone | 182.07 | C13H10O | 80.07 |
| | 120-46-7 | 8433 | Dibenzoylmethane | 224.08 | C15H12O2 | 73.54 |
| | 128-37-0 | 31404 | Butylated Hydroxytoluene | 220.18 | C15H24O | 82.89 |
| | 139776-09-3 | 569530 | S-Benzoyl-N-(p-nitrobenzylidene)thiohydroxylamine | 286.04 | C14H10N2O3S | 74.31 |
| | 143-07-7 | 3893 | Dodecanoic acid | 200.18 | C12H24O2 | 78.34 |
| | 149180-87-0 | 624073 | Butylaldehyde, 4-benzyloxy-4-[2,2,-dimethyl-4-dioxolanyl]- | 278.15 | C16H22O4 | 72.53 |
| | 150-86-7 | 5280435 | Phytol | 296.31 | C20H40O | 78.92 |
| | 177746-99-5 | 91692548 | Methyl 15-hydroxy-9,12-octadecadienoate | 310.25 | C19H34O3 | 76.43 |
| | 20548-62-3 | 590836 | Phthalic acid, bis(7-methyloctyl) ester | 418.31 | C26H42O4 | 78.82 |
| | 22599-96-8 | 22213932 | Cholestan-3-ol, 2-methylene-, (3.beta.,5.alpha.)- | 400.37 | C28H48O | 73.41 |
| | 24560-98-3 | 119250 | Oxiraneoctanoic acid, 3-octyl-, cis- | 298.25 | C18H34O3 | 77.99 |
| | 25360-09-2 | 19107815 | tert-Hexadecanethiol | 258.24 | C16H34S | 72.32 |
| | 2676-41-7 | 146287 | 6,9,12-Octadecatrienoic acid, methyl ester | 292.24 | C19H32O2 | 74.8 |
| | 26896-20-8 | 62838 | Neodecanoic acid | 172.15 | C10H20O2 | 72.5 |
| | 28108-99-8 | 34148 | Phosphoric acid, (1-methylethyl)phenyl diphenyl ester | 368.12 | C21H21O4P | 78.89 |
| | 29761-21-5 | 34697 | Phosphoric acid, isodecyl diphenyl ester | 390.20 | C22H31O4P | 73.13 |
| | 334-68-9 | 9548 | Dodecane, 1-fluoro- | 188.19 | C12H25F | 74.35 |
| | 33795-18-5 | 214694 | Phosphonic acid, (p-hydroxyphenyl)- | 174.01 | C6H7O4P | 76.11 |
| | 3443-82-1 | 5365676 | 9,12-Octadecadienoic acid (Z,Z)-, 2-hydroxy-1-(hydroxymethyl)ethyl ester | 354.28 | C21H38O4 | 79.87 |
| | 34909-69-8 | 631942 | Phosphoric acid, bis(4-methylphenyl) phenyl ester | 354.10 | C20H19O4P | 83.97 |
| | 373-49-9 | 445638 | Palmitoleic acid | 254.24 | C16H30O2 | 77.31 |
| | 56051-53-7 | 554084 | Cyclopropanebutanoic acid, 2-[[2-[[2-[(2-pentylcyclopropyl)methyl]cyclopropyl]methyl]cyclopropyl]methyl]-, methyl ester | 374.32 | C25H42O2 | 75.58 |
| | 57-10-3 | 985 | n-Hexadecanoic acid | 256.24 | C16H32O2 | 74.41 |
| | 57156-91-9 | 42151 | 2,5-Octadecadiynoic acid, methyl ester | 290.23 | C19H30O2 | 71.56 |
| | 60-33-3 | 5280450 | 9,12-Octadecadienoic acid (Z,Z)- | 280.24 | C18H32O2 | 74.4 |
| | 60609-53-2 | 5364688 | 8-Hexadecenal, 14-methyl-, (Z)- | 252.25 | C17H32O | 77.49 |
| | 74685-30-6 | 5364600 | 5-Eicosene, (E)- | 280.32 | C20H40 | 80.86 |
| | 76841-70-8 | 71403428 | E-2-Hexenyl benzoate | 204.12 | C13H16O2 | 73.99 |
| | 77-90-7 | 6505 | Tributyl acetylcitrate | 402.23 | C20H34O8 | 71.72 |
| | 78-31-9 | 6528 | Phosphoric acid, 4-methylphenyl diphenyl ester | 340.09 | C19H17O4P | 79.33 |
| | 816-19-3 | 102491 | Hexanoic acid, 2-ethyl-, methyl ester | 158.13 | C9H18O2 | 71.83 |
| | 82304-66-3 | 545303 | 7,9-Di-tert-butyl-1-oxaspiro(4,5)deca-6,9-diene-2,8-dione | 276.17 | C17H24O3 | 71.69 |
| | 84-77-5 | 6788 | Didecyl phthalate | 446.34 | C28H46O4 | 83.9 |
| | 85763-57-1 | 33865 | 11-Methyldodecanol | 200.21 | C13H28O | 78.32 |
| PVC 4 | 1225365[a] | 520263 | Phosphoric acid, 2-methylphenyl diphenyl ester | 340.09 | C19H17O4P | 86.56 |
| | 1919690[a] | 81591 | Benzoic acid, heptyl ester | 220.15 | C14H20O2 | 84.53 |
| | 1000308-89-8[a] | 3024584 | Phthalic acid, decyl nonyl ester | 432.32 | C27H44O4 | 70.78 |

| Sample | CAS | PubChem CID | Name according to NIST | Mass | Formula | Score |
|---|---|---|---|---|---|---|
| | 1000339-40-5[a] | 12151622 | 1,2-Cyclohexanedicarboxylic acid, dinonyl ester | 424.36 | C26H48O4 | 74.42 |
| | 1000339-74-3[a] | 91721826 | 1,2-Cyclohexanedicarboxylic acid, cyclohexylmethyl nonyl ester | 394.31 | C24H42O4 | 77.87 |
| | 1000340-22-6[a] | 9814973 | Benzoic acid, tridecyl ester | 304.24 | C20H32O2 | 80.46 |
| | 1000353-65-9[a] | 91714177 | Adipic acid, 3-heptyl tetradecyl ester | 440.39 | C27H52O4 | 76.38 |
| | 1000367-91-3[a] | 103653 | Benzoic acid, 2-methylbutyl ester | 192.12 | C12H16O2 | 77.5 |
| | 1000368-69-4[a] | 243678 | Benzoic acid, hept-2-yl ester | 220.15 | C14H20O2 | 76.04 |
| | 1000371-07-7[a] | 91719575 | Phthalic acid, 5-methylhex-2-yl heptadecyl ester | 502.40 | C32H54O4 | 75.96 |
| | 18699-48-4 | 29218 | 1,4-Benzenedicarboxylic acid, bis(2-methylpropyl) ester | 278.15 | C16H22O4 | 90.68 |
| | 20548-62-3 | 590836 | Phthalic acid, bis(7-methyloctyl) ester | 418.31 | C26H42O4 | 81.45 |
| | 29761-21-5 | 34697 | Phosphoric acid, isodecyl diphenyl ester | 390.20 | C22H31O4P | 80.26 |
| | 34909-69-8 | 631942 | Phosphoric acid, bis(4-methylphenyl) phenyl ester | 354.10 | C20H19O4P | 88.72 |
| | 59736-57-1 | 570433 | Benzoic acid 2-methylpentyl ester | 206.13 | C13H18O2 | 83.13 |
| | 76841-70-8 | 71403428 | E-2-Hexenyl benzoate | 204.12 | C13H16O2 | 73.45 |
| | 77-90-7 | 6505 | Tributyl acetylcitrate | 402.23 | C20H34O8 | 83.62 |
| | 78-31-9 | 6528 | Phosphoric acid, 4-methylphenyl diphenyl ester | 340.09 | C19H17O4P | 83.33 |
| | 84-64-0 | 6779 | 1,2-Benzenedicarboxylic acid, butyl cyclohexyl ester | 304.17 | C18H24O4 | 75.60 |
| | 84-76-4 | 6787 | 1,2-Benzenedicarboxylic acid, dinonyl ester | 418.31 | C26H42O4 | 83.79 |
| | 85-68-7 | 2347 | Benzyl butyl phthalate | 312.14 | C19H20O4 | 71.83 |
| | 94-50-8 | 66751 | Benzoic acid, octyl ester | 234.16 | C15H22O2 | 85.23 |
| PUR 1 | 112-61-8 | 8201 | Methyl stearate | 298.29 | C19H38O2 | 75.92 |
| | 128-37-0 | 31404 | Butylated Hydroxytoluene | 220.18 | C15H24O | 89.33 |
| PUR 2 | 1000128-20-5[a] | none | (+,-)-Epi-perhydrohistrionicotoxin | 295.29 | C19H37NO | 71.38 |
| | 1000195-87-0[a] | 550132 | 8,14-Seco-3,19-epoxyandrostane-8,14-dione, 17-acetoxy-3.beta.-methoxy-4,4-dimethyl- | 420.25 | C24H36O6 | 70.01 |
| | 1000303-02-6[a] | 6423312 | 7-Amino-1,3-dihydro-indol-2-one | 148.06 | C8H8N2O | 75.44 |
| | 128-37-0 | 31404 | Butylated Hydroxytoluene | 220.18 | C15H24O | 92.86 |
| | 149-57-5 | 8697 | Hexanoic acid, 2-ethyl- | 144.12 | C8H16O2 | 71.15 |
| | 2456-81-7 | 75567 | Pyridine, 4-(1-pyrrolidinyl)- | 148.10 | C9H12N2 | 81.91 |
| | 2566-91-8 | 6451414 | Oxiraneoctanoic acid, 3-octyl-, methyl ester, cis- | 312.27 | C19H36O3 | 76.43 |
| | 301-02-0 | 5283387 | 9-Octadecenamide, (Z)- | 281.27 | C18H35NO | 80.39 |
| | 5129-61-3 | 110444 | Heptadecanoic acid, 16-methyl-, methyl ester | 298.29 | C19H38O2 | 72.06 |
| | 584-84-9 | 11443 | Benzene, 2,4-diisocyanato-1-methyl- | 174.04 | C9H6N2O2 | 85.39 |
| | 61338-98-5 | 547892 | Benzeneethanamine, 2-fluoro-.beta.,3,4-trihydroxy-N-isopropyl- | 229.11 | C11H16FNO3 | 74.85 |
| | 76-25-5 | 6436 | Triamcinolone Acetonide | 434.21 | C24H31FO6 | 70.57 |
| | 823-40-5 | 13205 | 1,3-Benzenediamine, 2-methyl- | 122.08 | C7H10N2 | 71.37 |
| PUR 3 | 1000370-31-1[a] | 458684 | 4,4'-Di-tert-butyl-diphenylamine | 281.21 | C20H27N | 70.45 |
| | 1000370-31-3[a] | 117942 | Tert-octyldiphenylamine | 281.21 | C20H27N | 79.04 |
| | 1000400-90-8[a] | 291360 | benzaldehyde, 4-(ethylphenylamino)- | 225.12 | C15H15NO | 75.37 |
| | 1000408-33-2[a] | 91739821 | 2-[2-Methoxy-5-(1,1,3,3-tetramethylbutyl)phenyl]-2H-benzotriazole | 337.22 | C21H27N3O | 77.39 |
| | 6386-38-5 | 62603 | Benzenepropanoic acid, 3,5-bis(1,1-dimethylethyl)-4-hydroxy-, methyl ester | 292.20 | C18H28O3 | 70.30 |
| | 78-40-0 | 6535 | Triethyl phosphate | 182.07 | C6H15O4P | 74.93 |

| Sample | CAS | PubChem CID | Name according to NIST | Mass | Formula | Score |
|---|---|---|---|---|---|---|
| PUR 4 | 128-37-0 | 31404 | Butylated Hydroxytoluene | 220.18 | C15H24O | 77.47 |
| | 729-43-1 | 5484329 | Ethanone, 1-phenyl-, (1-phenylethylidene)hydrazone | 236.13 | C16H16N2 | 81.99 |
| PLA 1 | 1000382-54-3[a] | 91693137 | Carbonic acid, eicosyl vinyl ester | 368.33 | C23H44O3 | 78.36 |
| | 554-12-1 | 11124 | Methyl propionate | 88.05 | C4H8O2 | 74.34 |
| PLA 2 | 554-12-1 | 11124 | Methyl propionate | 88.05 | C4H8O2 | 72.33 |
| PLA 3 | not analyzed via GC-QTOF-MS/MS | | | | | |
| PLA 4 | 112-67-4 | 8206 | Palmitoyl chloride | 274.21 | C16H31ClO | 70.25 |
| | 112-80-1 | 445639 | Oleic Acid | 282.26 | C18H34O2 | 74.11 |
| | 143-07-7 | 3893 | Dodecanoic acid | 200.18 | C12H24O2 | 74.32 |
| | 1673-08-1 | 74288 | Hexadecanoic acid, cyclohexyl ester | 338.32 | C22H42O2 | 70.45 |
| | 5129-61-3 | 110444 | Heptadecanoic acid, 16-methyl-, methyl ester | 298.29 | C19H38O2 | 82.05 |
| | 57-10-3 | 985 | n-Hexadecanoic acid | 256.24 | C16H32O2 | 84.18 |
| | 57-11-4 | 5281 | Octadecanoic acid | 284.27 | C18H36O2 | 71.32 |

**Table S3. List of adipogenic chemicals based on the published literature.**

| Name | CAS | PubChem CID | References |
|---|---|---|---|
| 1-850 | 251310 -57-3 | 2765122 | (*42*) |
| 2,4,6-tribromophenol | 118-79-6 | 1483 | (*23*) |
| 2-ethylhexyl diphenyl phosphate | 1241-94-7 | 14716 | (*52*) |
| 3,5,6-trichloro-2-pyridinol | 6515-38-4 | 23017 | (*53*) |
| 3-tert-butyl-4-hydroxanisole | 121-00-6 | 8456 | (*16*) |
| 4-hexyl phenol | 2446-69-7 | 17132 | (*54*) |
| 4-n-octylphenol | 1806-26-4 | 15730 | (*55*) |
| 4-nonylphenol | 104-40-5 | 1752 | (*55, 56*) |
| 8:2 FTAcr | 27905-45-9 | 119747 | (*23*) |
| Acetamiprid | 135410-20-7 | 213021 | (*57*) |
| Acrylamide | 79-06-1 | 6579 | (*16*) |
| Allethrin | 584-79-2 | 11442 | (*58*) |
| Alpha naphthoflavone | 604-59-1 | 11790 | (*16*) |
| Azoxystrobin | 131860-33-8 | 3034285 | (*23*) |
| BDE-47 | 5436-43-1 | 95170 | (*23, 59*) |
| Benzyl butyl phthalate | 85-68-7 | 2347 | (*60-62*) |
| Biphenthrin | 82657-04-03 | 6442842 | (*63*) |
| Bisphenol A | 80-05-07 | 6623 | (*28, 57, 59-62, 64-68*) |
| Bisphenol A diglycidyl ether | 1675-54-3 | 2286 | (*67*) |
| Bisphenol F | 620-92-8 | 12111 | (*64, 65*) |
| Bisphenol S | 80-09-1 | 6626 | (*64-66, 68*) |
| Butylparaben | 94-26-8 | 7184 | (*62*) |
| Carboxymethylcellulose | 9000-11-7 | 24748 | (*16*) |
| Cetyl alcohol ethoxylate | 5274-61-3 | 4303686 | (*55*) |
| Chlorpyrifos | 2921-88-2 | 2730 | (*23, 53*) |
| Cypermethrin | 52315-07-8 | 2912 | (*23*) |
| Daidzein | 486-66-8 | 5281708 | (*69*) |
| DDE | 72-55-9 | 3035 | (*16*) |
| DDT | 50-29-3 | 3036 | (*16*) |
| Dechlorane plus | 13560-89-9 | 26111 | (*70*) |
| DEHP | 117-81-7 | 8343 | (*23, 71*) |
| Dexamethasone | 50-02-2 | 5743 | (*42, 72, 73*) |
| Diazinon | 333-41-5 | 3017 | (*74*) |

| Name | CAS | PubChem CID | References |
|---|---|---|---|
| Diethylene glycol dibenzoate | 120-55-8 | 8437 | (*48*) |
| Dibutyl phthalate | 84-74-2 | 3026 | (*23, 62*) |
| Dibutyltin | 1002-53-5 | 6484 | (*16*) |
| Dicamba | 1918-00-9 | 3030 | (*72*) |
| Diclofop-methyl | 51338-27-3 | 39985 | (*16*) |
| Di-iso-butyl phthalate | 84-69-5 | 6782 | (*62*) |
| Di-iso-decyl phthalate | 89-16-7 | 33599 | (*48*) |
| Di-iso-nonyl phthalate | 28553-12-0 | 590836 | (*48*) |
| Dioctyl sodium sulfosuccinate | 577-11-7 | 23673837 | (*75, 76*) |
| Diphenyl phosphate | 838-85-7 | 13282 | (*77*) |
| Ethylparaben | 120-47-8 | 8434 | (*62*) |
| Fenthion | 55-38-9 | 3346 | (*58*) |
| Fludioxonil | 131341-86-1 | 86398 | (*57*) |
| Fluoxastrobin | 361377-29-9 | 11048796 | (*23*) |
| Flusilazole | 85509-19-9 | 73675 | (*57*) |
| Flutamide | 13311-84-7 | 3397 | (*42*) |
| Forchlorfenuron | 68157-60-8 | 93379 | (*57*) |
| Genistein | 446-72-0 | 5280961 | (*69*) |
| Glyphosate | 1071-83-6 | 3496 | (*16*) |
| GW3965 | 405911-17-3 | 16078973 | (*42*) |
| Halosulfuron-methyl | 100784-20-1 | 91763 | (*16*) |
| HBCD | 3194-55-6 | 18529 | (*78*) |
| Hexafluorobisphenol A | 1478-61-1 | 73864 | (*42*) |
| Imidacloprid | 138261-41-3 | 86287518 | (*73*) |
| Isopropylated triphenyl phosphate | 78-30-8 | 6527 | (*79*) |
| Isoxaflutole | 141112-29-0 | 84098 | (*72*) |
| Lactofen | 77501-63-4 | 62276 | (*16*) |
| Lauryl alcohol ethoxylate (4) | 4536-30-5 | 24750 | (*55*) |
| LG100268 | 153559-76-3 | 3922 | (*42*) |
| MEHP | 4376-20-9 | 20393 | (*71*) |
| Methylparaben | 99-76-3 | 7456 | (*62, 80*) |
| Mono(2-ethylhexyl) phthalate | 4376-20-9 | 20393 | (*81*) |
| Monosodium glutamate | 142-47-2 | 23672308 | (*16*) |
| Musk xylene | 81-15-2 | 62329 | (*62*) |
| Nonylphenol ethoxylate (1-2) | 9016-45-9 | 24773 | (*55*) |

| Name | CAS | PubChem CID | References |
|---|---|---|---|
| Nonylphenol ethoxylate (20) | N/A | N/A | (*55*) |
| Nonylphenol ethoxylate (4) | N/A | N/A | (*55*) |
| Nonylphenol ethoxylate (6) | N/A | N/A | (*55*) |
| Nonylphenol ethoxylate (9-10) | N/A | N/A | (*55*) |
| Octylphenol ethoxylate (3) | 2315-67-5 | 5590 | (*55*) |
| P-80 | 9005-65-6 | 5284448 | (*16*) |
| PBDE 99 | 60348-60-9 | 36159 | (*82*) |
| PCB 153 | 35065-27-1 | 37034 | (*59*) |
| Permethrin | 52645-53-1 | 40326 | (*23*) |
| PFHxS | 3871-99-6 | 23678874 | (*83*) |
| PFNA | 375-95-1 | 67821 | (*83*) |
| PFOA | 335-67-1 | 9554 | (*59, 83, 84*) |
| PFOS | 1763-23-1 | 74483 | (*83*) |
| Pioglitazone | 112529-15-4 | 60560 | (*58*) |
| Pirinixic acid | 50892-23-4 | 5694 | (*83*) |
| Prallethrin | 23031-36-9 | 9839306 | (*58*) |
| Propylparaben | 94-13-3 | 7175 | (*62, 80*) |
| Pymetrozine | 123312-89-0 | 9576037 | (*57*) |
| Pyraclostrobin | 175013-18-0 | 6422843 | (*23*) |
| Pyrimethanil | 53112-28-0 | 91650 | (*58*) |
| Quinoxyfen | 124495-18-7 | 3391107 | (*57, 58*) |
| Quizalofop-p-ethyl | 100646-51-3 | 1617113 | (*72*) |
| Retinoic acid | 302-79-4 | 444795 | (*58*) |
| Rosiglitazone | 122320-73-4 | 77999 | positive control |
| Span 80 | 1338-43-8 | 9920342 | (*76*) |
| Spirodiclofen | 148477-71-8 | 177863 | (*57*) |
| TBEP | 78-51-3 | 6540 | (*23*) |
| TBPH | 26040-51-7 | 117291 | (*23*) |
| TDBPIC | 52434-90-9 | 103634 | (*23*) |
| Tebuconazole | 107534-96-3 | 86102 | (*58*) |
| Tebupirimfos | 96182-53-5 | 93516 | (*57*) |
| Tert-butylphenyl diphenyl phosphate | 56803-37-3 | 158333 | (*23*) |
| Tetrabromobisphenol A | 79-94-7 | 6618 | (*42, 58, 85*) |
| Tetrachlorobisphenol A | 79-95-8 | 6619 | (*42*) |
| Tolyfluanid | 731-27-1 | 12898 | (*86*) |
| Tomadol 1-9 | N/A | N/A | (*55*) |
| Tonalide | 1506-02-1 | 89440 | (*62*) |

| Name | CAS | PubChem CID | References |
|---|---|---|---|
| Tributyltin | 687-73-3 | 3032732 | (*42, 59, 62, 87-92*) |
| Triclocarban | 101-20-2 | 7547 | (*93*) |
| Tridecyl alcohol ethoxylate (9) | 78330-21-9 | N/A | (*55*) |
| Trifloxystrobin | 141517-21-7 | 11665966 | (*23*) |
| Triflumizole | 68694-11-1 | 91699 | (*57, 94*) |
| Tri-m-cresyl phosphate | 563-04-2 | 11232 | (*48*) |
| Tri-n-butyl phosphate | 126-73-8 | 31357 | (*23*) |
| Triphenyl phosphate | 115-86-6 | 8289 | (*23, 77, 79, 95*) |
| Triphenyltin | 76-87-9 | 6327657 | (*57, 58*) |
| Tris(4-tert-butylphenyl) phosphate | 78-33-1 | 6530 | (*23*) |
| Troglitazone | 97322-87-7 | 5591 | (*59, 61, 79*) |
| TTBP | 732-26-3 | 12902 | (*58*) |
| Zoxamide | 156052-68-5 | 122087 | (*57*) |

**Table S4. Comparison of the adipogenic effects of plastic extracts and the abundance of three metabolic disrupting chemicals detected in at least three samples,** diphenyl phosphate (DPP), 2-ethylhexyl diphenyl phosphate (EHDP), and triphenyl phosphate (TPP) in the LC-QTOF-MS/MS.

| | **Lipid droplet count** | | **Raw abundance** | | | | | | | | | | | | |
|---|---|---|---|---|---|---|---|---|---|---|---|---|---|---|---|
| Samples | Median | $EC_{50}$ | DPP 1 [a] | DPP 2 [a] | DPP 3 [a] | DPP 4 [a] | EHDP | TPP 1 [a] | TPP 2 [a] | TPP 3 [a] | TPP 4 [a] | TPP 5 [a] | TPP 6 [a] | TPP 7 [a] | TPP 8 [a] | TPP 9 [a] |
| HDPE 1 | 164.6 | 3.1 | 0 | 0 | <LOD | <LOD | 0 | 0 | 0 | 1542 | 0 | 0 | 0 | 0 | 0 | 0 |
| HDPE 2 | 161.4 | 3.1 | 0 | 0 | <LOD | <LOD | 0 | 0 | 0 | 0 | <LOD | 0 | 0 | 0 | 0 | 0 |
| HDPE 3 | 360.8 | 3.1 | 0 | 0 | <LOD | <LOD | 0 | 0 | 0 | 0 | 0 | 0 | 0 | 2426 | 0 | 0 |
| HDPE 4 | 369.1 | 3.1 | 0 | 0 | <LOD | <LOD | 4.7 | 0 | 0 | 0 | <LOD | 0 | 0 | 0 | 0 | 0 |
| LDPE 1 | 447.3 | 3.1 | 0 | 0 | <LOD | <LOD | 5.7 | 0 | 0 | 0 | <LOD | 0 | 0 | 0 | 0 | 0 |
| LDPE 2 | 373.2 | 3.1 | 0 | 0 | <LOD | <LOD | 0 | 0 | 0 | 24.6 | 0 | 0 | 0 | 0 | 0 | 137.4 |
| LDPE 3 | 294.4 | 3.1 | 0 | 0 | <LOD | <LOD | 0 | 0 | 0 | 0 | 0 | 0 | 0 | <LOD | 0 | 0 |
| LDPE 4 | 1352 | 1.59 | 0 | 0 | <LOD | <LOD | 0 | 0 | 0 | 0 | 974.2 | 0 | 0.1 | 0 | 0 | 0 |
| PS 1 | 419.9 | 3.1 | 0 | 0 | <LOD | 0 | 0 | 14.6 | 0 | 0 | 0 | 0 | 0 | 0 | 0 | 0 |
| PS 2 | 1333 | 2.05 | 9.8 | 0 | <LOD | <LOD | 0 | 0 | 16.7 | 0 | 0 | 0 | 0 | 0 | <LOD | 0 |
| PS 3 | 186.1 | 3.1 | 0 | 0 | <LOD | 0 | 0 | 0 | 0 | 0 | 0 | 0 | 0 | 0 | 0 | 0 |
| PS 4 | 123.7 | 3.1 | 0 | 0 | 0 | <LOD | 0 | 0 | 0 | 0 | <LOD | 0 | 0 | 0 | 0 | 0 |
| PP 1 | 197.3 | 3.1 | 0 | 0 | <LOD | <LOD | 0 | 0 | 1633 | 0 | <LOD | 0 | 0 | 0 | 0 | 984.3 |
| PP 2 | 413 | 3.1 | 0 | 0 | 0 | <LOD | 0 | 0 | 0 | 2.2 | <LOD | 0 | 0 | 0 | 0 | 0 |
| PP 3 | 1742 | 1.4 | 0 | 0 | <LOD | 0 | 0 | 1184 | 0 | 59.3 | <LOD | 0 | 0 | 0 | 0 | 0 |
| PP 4 | 2927 | 0.4 | 0 | 0 | 0 | <LOD | 0 | 0 | 0 | 0 | <LOD | 0 | 0 | 0 | 0 | 0 |
| PP 5 | 119.1 | 3.1 | 0 | 0 | 0 | <LOD | 0 | 0 | 0 | 0 | <LOD | 0 | 0 | 0 | 0 | 0 |
| PET 1 | 99.12 | 3.1 | 0 | 0 | 0 | <LOD | 0 | 0 | 0 | 0 | <LOD | 0 | 0 | 0 | 0 | 0 |
| PET 2 | 185 | 3.1 | 0 | 0 | 0 | <LOD | 0 | 0 | 0 | 0 | <LOD | 0 | 0 | 0 | 0 | 0 |
| PET 3 | 139.4 | 3.1 | 0 | 0 | 0 | <LOD | 0 | 0 | 0 | 0 | <LOD | 0 | 0 | 0 | 0 | 0 |
| PET 4 | 122.4 | 3.1 | 0 | 0 | <LOD | <LOD | 0 | 0 | 0 | 0 | 0 | 0 | 0 | 0 | 0 | 0 |
| PET 5 | 183.7 | 3.1 | 0 | 0 | <LOD | <LOD | 0 | 0 | 0 | 0 | <LOD | 0 | 0 | <LOD | 0 | 0 |
| PVC 1 | 171.4 | 3.1 | 0 | 0 | 0 | <LOD | 0 | 0 | 0 | 0 | <LOD | 0 | 0 | 0 | 0 | 0 |
| PVC 2 | 3302 | 0.85 | 4251 | 2466 | 437,900 | <LOD | 0 | 0 | 0 | 356,178 | <LOD | 0 | 0 | 0 | 0 | 0 |
| PVC 3 | 903.5 | 1 | 0 | 0 | 0 | <LOD | 0 | 0 | 0 | 0 | 554,290 | 0 | 27.1 | 0 | 0 | 0 |

| | **Lipid droplet count** | | **Raw abundance** | | | | | | | | | | | | | |
|---|---|---|---|---|---|---|---|---|---|---|---|---|---|---|---|---|
| Samples | Median | $EC_{50}$ | DPP 1 [a] | DPP 2 [a] | DPP 3 [a] | DPP 4 [a] | EHDP | TPP 1 [a] | TPP 2 [a] | TPP 3 [a] | TPP 4 [a] | TPP 5 [a] | TPP 6 [a] | TPP 7 [a] | TPP 8 [a] | TPP 9 [a] |
| PVC 4 | 3044 | 0.53 | 0 | 0 | <LOD | 315,591 | 2040 | 0 | 0 | 0 | <LOD | 4094 | 544,411 | <LOD | 4257 | 10.7 |
| PUR 1 | 1340 | 0.61 | 0 | 0 | 0 | <LOD | 1.6 | 0 | 0 | 63.9 | <LOD | 771.1 | 0 | 0 | 0 | 0 |
| PUR 2 | 969.4 | 0.33 | 0 | 0 | 0 | <LOD | <LOD | 0 | 0 | 0 | <LOD | 22.3 | 0 | 0 | 0 | 0 |
| PUR 3 | 1186 | 0.32 | 0 | 0 | <LOD | <LOD | 65.8 | 95.7 | 1.4 | 31.3 | 0 | 0 | 0 | 0 | 0 | 0 |
| PUR 4 | 2195 | 1.05 | 0 | 0 | 0 | <LOD | 438.7 | 0 | 0 | 0 | <LOD | 0 | 0.1 | 0 | 0 | 0 |
| PLA 1 | 281.7 | 3.1 | 0 | 0 | <LOD | <LOD | 0 | 0 | 0 | 0 | 0 | 0 | 33.4 | 0 | <LOD | 0 |
| PLA 2 | 203.7 | 3.1 | 0 | 0 | <LOD | <LOD | 0 | 0 | 0 | 0 | 0 | 0 | 0 | <LOD | 0 | 0 |
| PLA 3 | not analyzed via LC-QTOF-MS/MS | | | | | | | | | | | | | | | |
| PLA 4 | 271.8 | 3.1 | 0 | 0 | <LOD | <LOD | 0 | 0 | 0 | 0 | <LOD | 0 | 0 | 0 | 0 | 0 |

**Table S5. List of all used chemicals and consumables.**

| chemical/consumable | supplier | further information |
|---|---|---|
| AdipoRed assay kit (NileRed) | Lonza | N3013 |
| ATP (adenosine-5'-triphosphate disodium salt) | PanReac AppliChem | ≥98 % (HPLC), CAS: 987655 |
| Bovine calf serum iron-supplemented | Sigma | 12138C |
| Charcoal-stripped fetal bovine serum | Gibco | 12676029 |
| CDTA (1,2-cyclohexanedinitrilotetraacetic acid) | Sigma | ≥99 % (KT), CAS: 125572954 |
| Dexamethasone | Sigma | ≥98 % (HPLC), CAS: 50022 |
| DMEM/F-12 with phenol red | Gibco | 31331093 |
| DMEM/F-12 without phenol red | Gibco | 21041025 |
| DMEM high glucose | Gibco | 31966047 |
| D-Luciferin, monopotassium salt | Thermo Scientific | ≥99.7 % (HPLC), 88294 |
| DL-dithiothreitol | Sigma | ≥98 % (HPLC), CAS: 3483123 |
| DMSO (dimethyl sulfoxide) | Sigma | ≥99.5 % (GC), CAS: 67685 |
| EDTA (ethylenediaminetetraacetic acid) | Sigma | ≥99 % (titration), CAS: 60004 |
| Fetal bovine serum | Gibco | 10270-106 |
| G 418 disulfate | Sigma | ≥450 U/mg, CAS: 108321422 |
| Glycerol | Sigma | ≥99 % (GC), CAS: 56815 |
| HEPES | VWR | L1613 |
| IBMX (3-isobutyl-1-methylxanthine) | Sigma | ≥99 % (HPLC), CAS: 28822587 |
| Insulin, human recomb., zinc solution | Gibco | 12585014, 27 U/mg |
| Magnesium carbonate hydroxide pentahydrate | Sigma | M5671, CAS: 56378724 |
| Magnesium sulfate heptahydrate | Sigma | ≥98 %, CAS: 10034998 |
| Methanol | Sigma | ≥99.8 % (HPLC), 322415 |
| NucBlue live cell ready probes reagent | Life technologies | R37605 |
| Penicillin/streptomycin | VWR | L0022-100 |
| Rosiglitazone | Sigma | ≥98 % (HPLC), CAS: 122320734 |
| Sterile T75 cell culture flasks | Thermo Scientific | 156499 |
| Tris base | Sigma | ≥99 % (titration), CAS: 77861 |
| Triton™ X-100 | Sigma | laboratory grade, CAS: 9002-93-1 |
| 96-well cell culture plates (black with transparent bottom) | Greiner | Cellstar 655090 |
| 384-well cell culture plates (white with transparent bottom) | Greiner | Cellstar 738-0062 |

**Dataset S1 (separate file).** Dataset S1 – SM Excel

- Excel Table S1 contains the tentatively identified chemicals in plastics using LC-QTOF-MS/MS
- Excel Table S2 contains the identity of the 25 chemicals with the highest identification score and abundance in the plastic samples
- Excel Table S3 contains the imaging data (means of the individual replicates) for the endpoint nuclei count (adipogenesis assay)
- Excel Table S4 contains the imaging data (means of the individual replicates) for the endpoint lipid droplet (adipogenesis assay)
- Excel Table S5 contains the imaging data (means of the individual replicates) for the endpoint adipocyte count (adipogenesis assay)
- Excel Table S6 contains the imaging data (means of the individual replicates) for the endpoint mature adipocyte count (adipogenesis assay)
- Excel Table S7 contains the imaging data (means of the individual replicates) for the endpoint lipid droplet intensity [RFU] (adipogenesis assay)
- Excel Table S8 contains the imaging data (means of the individual replicates) for the endpoint lipid droplet area [pixel] ) (adipogenesis assay)
- Excel Table S9 contains the PPARγ activity (individual replicates) [%] (PPARγ CALUX)
